# Persistent Malignant Stem Cells Drive Relapse After Transplantation in the Myelodysplastic Neoplasms

**DOI:** 10.64898/2026.10.28.761431

**Authors:** Benjamin Kroger, Roni Tamari, Carmelo Gurnari, Yuanyuan Ji, Yazan Madanat, Yi Huang, Reiko Tachibana, Toby Thomas, Albert Son, Peter Carlsgaard, Clayton Jackson, Vivian Irizarry-Gatell, Alejandro Marinos, Zachary Brady, Prapti Patel, Valeria Visconte, Abhisar Cherukuri, Madhuri Vusirikala, Asmita Mishra, Amy McLemore, Jerry Yuen, Bao Duy An Hoang, Russell Vittrup, Julio Alvarenga, Robert Collins, David A. Sallman, Jaroslaw Maciejewski, Stephen S. Chung

**Affiliations:** Division of Hematology/Oncology, Department of Internal Medicine, University of Texas Southwestern Medical Center, Dallas, TX; Adult Bone Marrow Transplant Service, Memorial Sloan Kettering Cancer Center, New York, NY; Department of Translational Hematology and Oncology Research, Taussig Cancer Institute, Cleveland Clinic, Cleveland, OH; Department of Biomedicine and Prevention, University of Rome Tor Vergata, Rome, Italy; Moffitt Cancer Center, Tampa, FL

**Keywords:** myelodysplastic neoplasms, cancer stem cells, therapeutic resistance, measurable residual disease, single-cell multi-omics, lineage tracing, cell surface markers

## Abstract

The myelodysplastic neoplasms (MDS) are initiated in hematopoietic stem cells (HSCs). Allogeneic transplantation is the only curative treatment, but relapse remains common. To detect rare MDS HSCs, we developed an ultra-low-input DNA sequencing assay and applied it to a multicenter post-transplant cohort. HSC mutation detection preceded relapse by a mean of 98 days and strongly predicted relapse (95.7% vs. 30.3%; hazard ratio, 8.8; *P*=1.5×10⁻⁷). Mutations detected only in unfractionated marrow did not predict relapse. We developed methods to measure donor-host HSC chimerism, improving relapse prediction sensitivity. Single-cell protein, DNA, and RNA analyses with mitochondrial lineage tracing confirmed MDS cell enrichment within immunophenotypically- and transcriptionally-defined HSCs, identifying shared ribosomal gene downregulation and candidate mechanisms of venetoclax sensitivity. Venetoclax/azacytidine depleted detectable MDS HSCs in longitudinally evaluated patients. These findings establish that HSC mutation detection predicts post-transplant relapse, provide a rationale for preemptive intervention, and define shared biological features of transplant-resistant MDS HSCs.

**Statement of Significance:** We show that the cellular compartment harboring residual MDS-associated mutations determines their prognostic significance after allogeneic transplantation. Our ultra-low-input sequencing assay enables highly specific stem cell-based relapse prediction without requiring patient-specific customization. Integrated single-cell analyses reveal shared biological programs and candidate therapeutic vulnerabilities in persistent malignant hematopoietic stem cells.

## Introduction

The myelodysplastic neoplasms (MDS) are a heterogeneous group of bone marrow failure disorders initiated and sustained by aberrant hematopoietic stem cells (MDS HSCs)^1–4^. MDS HSCs persist despite standard therapies, including hypomethylating agents (HMAs)^5^ and lenalidomide^6^, making replacement of these cells by allogeneic hematopoietic cell transplantation (alloHCT) the only curative treatment^7^. However, relapse occurs after up to 40% of transplants^8–10^ and is associated with extremely poor outcomes^11–13^. Early detection of measurable residual disease (MRD) after transplantation could enable intervention at low disease burden and improve the curative potential of alloHCT. However, widely applicable approaches for identifying patients at highest risk of post-transplant relapse remain lacking.

Although MDS HSCs have been detected in transplanted patients who subsequently relapse^14^, whether their persistence early after alloHCT reliably identifies patients destined to relapse remains unclear. Conversely, it is unknown whether patients with durable remissions fully eradicate MDS HSCs or instead harbor them at low levels that lack clinical significance. Addressing these questions has been limited by the scarcity of HSCs and the technical challenges associated with isolating these rare cells from surveillance bone marrow specimens for genomic analysis. Previous studies using bulk^4^ and single-cell^15^ DNA sequencing have provided important insights into HSC-driven disease initiation and clonal evolution outside the transplant setting. However, these approaches relied on patient-specific sequencing probes or primers, limiting clinical scalability. In addition, the use of whole genome amplification (WGA) in these studies may introduce biases that compromise accurate mutation detection and quantification. A recent study used droplet digital PCR (ddPCR) to detect MDS HSCs after alloHCT^14^, but it similarly required patient-specific assay design.

To overcome these limitations, we developed an ultra-low-input targeted DNA sequencing assay encompassing 261 myeloid malignancy-associated genes^16^ that does not require patient-specific customization, enabling high-throughput detection of MDS-associated somatic mutations in small numbers of purified HSCs. Applying this assay to post-transplant bone marrow (BM) specimens from MDS patients across four academic centers, we found that detection of MDS-associated mutations in HSCs early after alloHCT identified patients destined to relapse. We additionally leveraged donor-and host-specific single-nucleotide variants (SNVs) to precisely track donor:host HSC chimerism after alloHCT, finding that detection of host HSCs further improved the sensitivity of relapse prediction. To orthogonally define the cellular compartment harboring persistent mutations, we integrated single-cell DNA- and RNA-sequencing with mitochondrial variant-based lineage tracing to distinguish donor and residual MDS clones after alloHCT. This approach confirmed preferential enrichment of residual MDS cells among immunophenotypically- and transcriptionally-defined HSCs and revealed molecular programs associated with disease persistence, sensitivity to venetoclax-containing salvage therapies, and other therapeutic vulnerabilities. Together, these studies establish a new paradigm for stem cell-based post-transplant disease monitoring and enable direct biological interrogation of the cells responsible for post-transplant disease persistence and relapse.

## Results

### Accurate identification and quantification of somatic mutations from limiting numbers of cells

To enable detection of mutated hematopoietic stem and progenitor cells (HSPCs) following alloHCT, we developed a method for high-throughput targeted DNA sequencing of rare cells. The accuracy of sequencing rare cells is limited by allelic drop-out and PCR amplification bias. To overcome this, we combined a method optimized to obtain high library complexity from low-quality samples with a hybridization capture-based sequencing panel covering 261 myeloid malignancy-associated genes^16^, as well as 11 HLA genes (**Supp. Table 2**). Unique dual index (UDI) primer sequences were used to enhance the specificity of PCR duplicate identification (**Fig.1A**). To evaluate assay performance, we analyzed an AML sample harboring six pathogenic somatic variants and compared their variant allele frequencies (VAFs) as detected from 10 ng and 500 ng of DNA with their clonal frequencies as determined by single-cell DNA sequencing (scDNA-seq). Detected VAFs were highly concordant with clonal frequencies (*R*² = 0.99 for 500 ng and *R*² = 0.90 for 10 ng), and for each variant, VAFs differed by <10% across both DNA input quantities and compared with scDNA-seq data (**Supp.Fig.1**).

**Figure 1.**
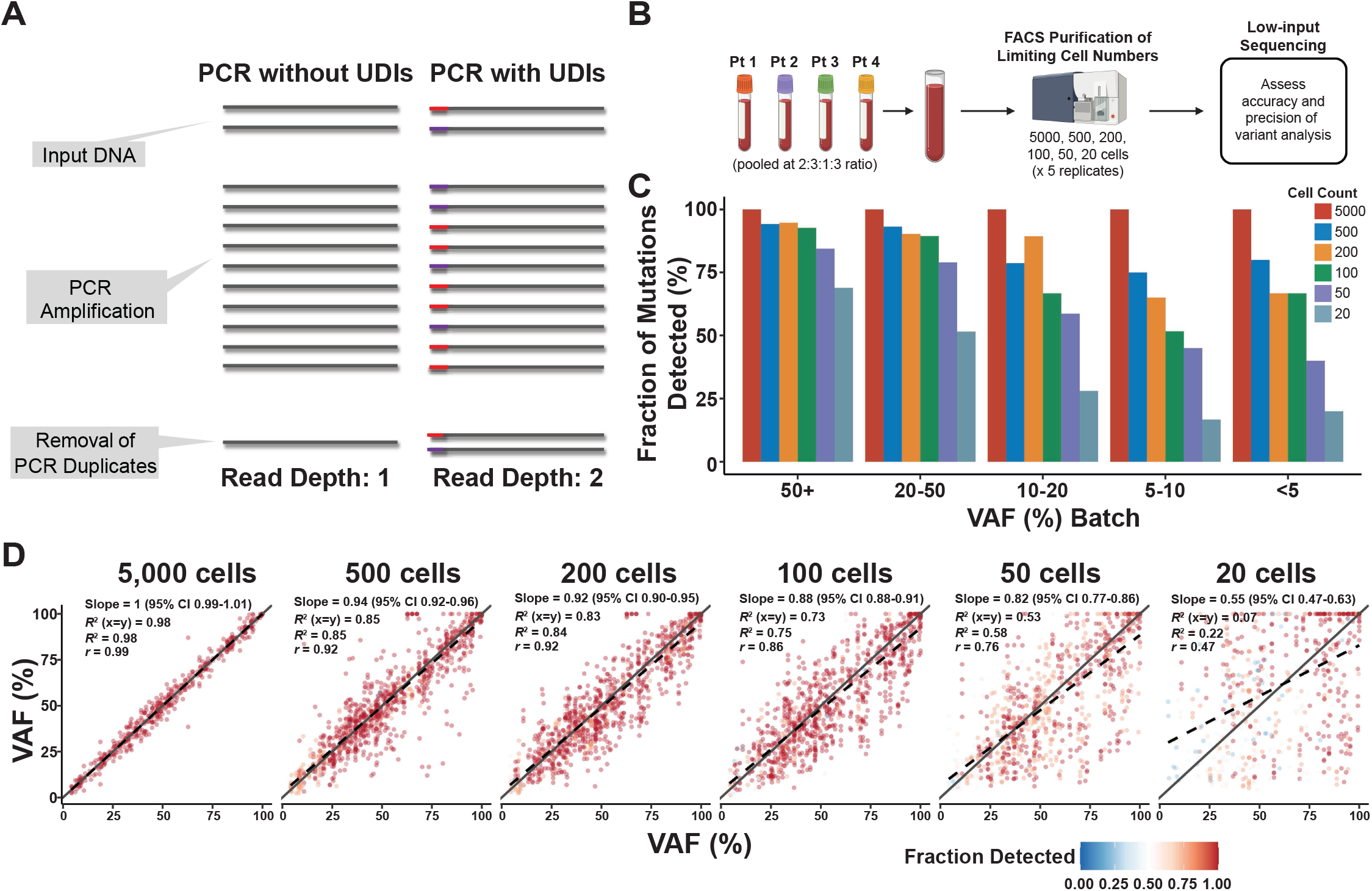
Accurate identification and quantification of somatic mutations from limiting numbers of cells. (**A**) Schematic comparison of PCR duplicate removal strategies. Conventional approaches (left) collapse reads based on identical start and end coordinates, whereas unique dual index (UDI) methods (right) use molecular barcodes to accurately identify and consolidate true PCR duplicates, increasing effective read depth. (**B**) Experimental design for low-input limit-of-detection analysis. Four patient samples were pooled at a 2:3:1:3 ratio and limiting numbers of cells (5,000, 500, 200, 100, 50, and 20 cells, n=5 per cell number) were purified for sequencing and variant detection. (**C**) Fraction of mutations detected across cell input levels stratified by variant allele frequency (VAF) bins. Detection sensitivity declines with both decreasing input and lower VAF. (**D**) Correlation of VAFs between each cell input (x-axis) and the mean VAF from 5,000-cell input (y-axis) across five replicates. Each point represents a variant; color indicates the fraction of replicates in which each variant was detected (out of 5). Solid and dashed lines indicate the x=y identity line and linear regression fit, respectively. Pearson correlation coefficient (*r*), linear model slope with 95% confidence interval, and *R²* are shown for each condition.

We next benchmarked the sensitivity of this method by applying it to detect somatic and germline variants in limiting numbers (5,000, 500, 200, 100, 50, and 20) of leukemic blasts (CD45^lo^SSC^lo^) purified from a pool of four AML patient samples, with five replicates per cell input (**Fig.1B**), obtaining approximately 8 million reads per specimen. Detection depended on both cell input and VAF (**Fig.1C**, **Supp. Table 3**). Among 229 variants with a measured VAF >10% from 5,000 cells, 172 (75%) were detected in ≥4 of 5 replicates with 50-cell input. With 20-cell input, 98 of 229 (43%) variants were detected in ≥4 of 5 replicates and 136 of 229 (59%) in ≥3 of 5 replicates. We obtained similar results when restricting our analysis to 15 pathogenic variants known to be present in the four patients at VAFs >5%. Nine of 15 (60%) variants and 11 of 15 (73%) variants were detectable in ≥3 of 5 replicates for the 20- and 50-cell inputs, respectively. With 200-cell input, all 15 variants were detectable in ≥4 of 5 replicates. We also compared the VAFs of somatic and germline variants detected at each cell number with the average VAF detected from 5,000 cells (**Fig.1D**, **Supp. Table 3**). With input of as few as 50 cells, VAF estimates correlated with measurements from 5,000 cells (Pearson *r* = 0.76; slope = 0.82, 95% CI 0.77– 0.86). With 20-cell input, concordance decreased (Pearson *r* = 0.47; slope = 0.55, 95% CI 0.47– 0.63), but many variants remained detectable. Together, these results establish a sensitive, scalable, and broadly applicable ultra-low-input targeted DNA sequencing assay well-suited for genomic interrogation of rare cell populations.

### MDS-associated mutations can be detected exclusively in HSCs well in advance of clinical relapse

We next sought to determine if MDS HSCs are present at detectable levels following alloHCT prior to clinical relapse. We thus identified four patients who relapsed after alloHCT and analyzed BMs taken at time points without clinical evidence of disease and more than three months prior to relapse (**Supp. Table 4**). To quantify disease burden across the hematopoietic hierarchy, we performed fluorescence-activated cell sorting (FACS) to purify HSCs combined with multipotent progenitors (MPPs, together referred to as HSCs/MPPs: lineage[lin]^-^CD34^+^CD38^-^CD45RA^-^), given that they sit atop the hematopoietic hierarchy as a mixed and heterogeneous population^17,18^. We separately purified lymphoid-primed multipotent progenitors (LMPPs, lin^-^CD34^+^CD38^-^CD90^-^CD45RA^+^)^19–21^, given that they may harbor incipient leukemia stem cells (LSCs)^20^, as well as restricted progenitors (referred to hereafter as progenitors, lin^-^ CD34^+^CD38^+^)^22^.

In each case, residual MDS cells, identified by the presence of driver mutations, were confined to the multipotent lin⁻CD34⁺CD38⁻ compartment, comprising HSCs, MPPs, and LMPPs. MDS cells were undetectable in progenitors or unfractionated BM (bulk) despite much higher sequencing depth. In **MSK MDS-001** (*DNMT3A* R882H and *IDH2* R140Q), both mutations were detected at D+179 within HSC/MPPs (74× from 102 cells) and LMPPs (164× from 577 cells) but were absent from progenitors and bulk BM (506× and 369× coverage, respectively, **Fig.2A**). In **UTSW 1706** (*ASXL1* E635fs) and **UTSW 2155** (*TP53* E286K), mutations were detectable exclusively within HSC/MPPs at D+84 and D+98 (64× from 190 cells and 46× from 75 cells, respectively) and were not identified in LMPPs, restricted progenitors, or bulk BM despite substantially greater coverage (**Fig.2B-C**). Similarly, in **UTSW 537** (*TP53* P75Lfs and Y220C), mutations were confined to the lin⁻CD34⁺CD38⁻ fraction at D+146 (105× from 258 cells) and remained undetectable in progenitors and bulk BM (**Fig.2D**).

**Figure 2.**
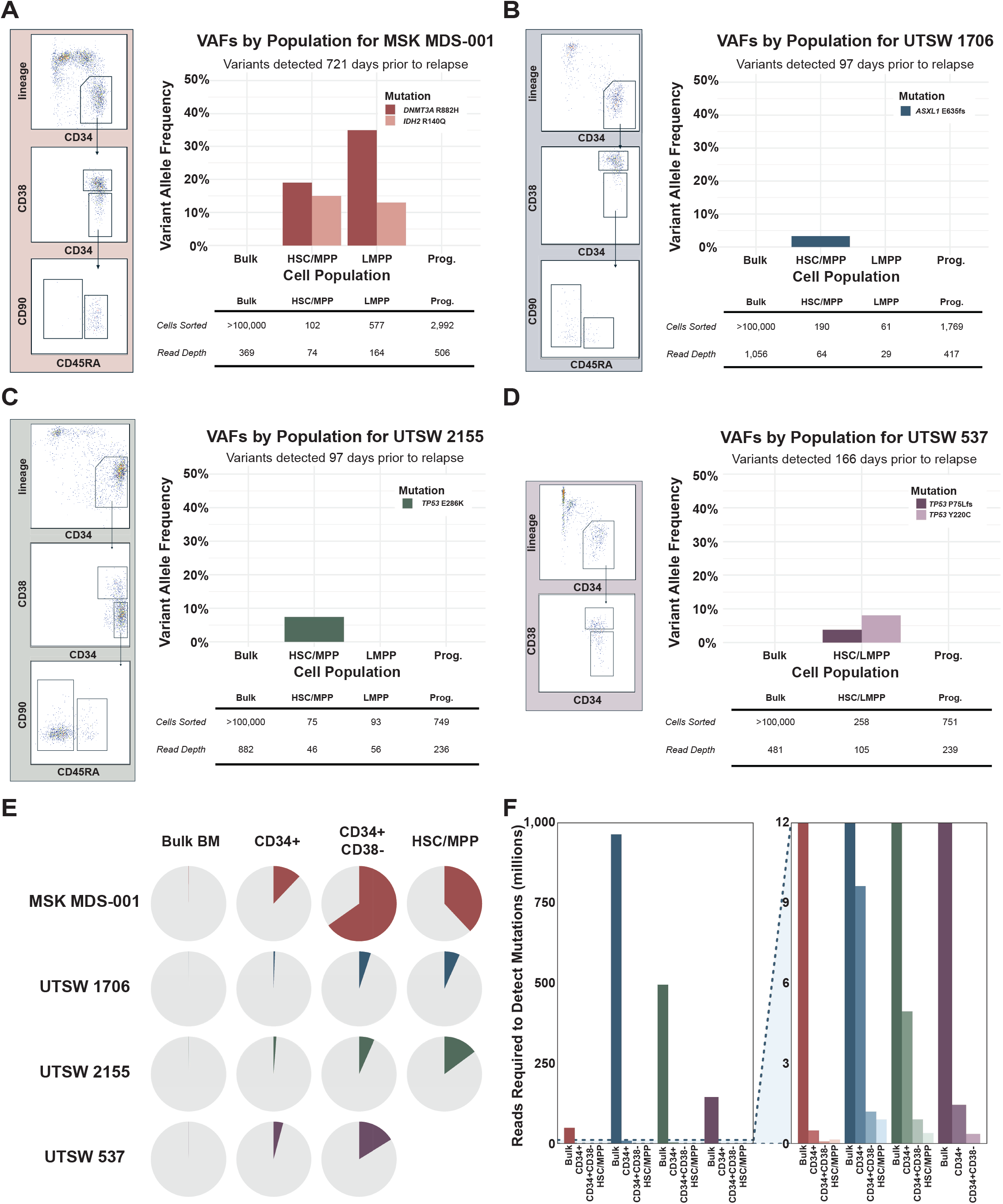
MDS-associated mutations can be detected exclusively in HSCs well in advance of clinical relapse. (**A-D**) Variant allele frequencies (VAFs) of pathogenic variants detected by low-input DNA sequencing within unfractionated bone marrow (Bulk), hematopoietic stem cells and multipotent progenitors (HSC/MPPs), lymphoid-primed multipotent progenitors (LMPPs), and restricted progenitor cells (Prog.) purified from **MSK MDS-001** (**A**), **UTSW 1706** (**B**), **UTSW 2155** (**C**), and **UTSW 537** (**D**). Flow cytometric gates used to isolate each population are shown to the left of each bar plot. Tables below each plot indicate the number of cells sorted and the mean sequencing read depth across target regions for each population. (**E**) Estimated MDS disease burden across populations, calculated as twice the VAF of the most abundant heterozygous pathogenic mutation detected in each sample. (**F**) Estimated sequencing depth required to detect the most abundant pathogenic mutation in each population, calculated as the depth needed to obtain three mutated reads.

We next combined HSC/MPP frequencies measured by flow cytometry with mutant cell fractions from our sequencing data to estimate the frequency of mutated HSC/MPPs out of bulk BM, CD34^+^, or CD34^+^CD38^-^ cells (**Fig.2E**). We estimated the read depth required to detect mutated cells from these populations (**Fig.2F**), finding that 49,758,280 to 963,587,216 reads would be required from bulk BM, 497,583 to 9,635,873 reads from CD34^+^ cells, and 92,035 to 1,197,329 reads from CD34^+^CD38^-^ cells. These results show that MDS can persist exclusively in HSC/MPPs after transplant long before clinical relapse, and that evaluation of HSC/MPPs is necessary to maximize the sensitivity of post-transplant disease assessment.

### Persistent MDS HSCs identify patients destined to relapse

To determine whether early post-transplant persistence of MDS HSCs predicts relapse, we analyzed 91 patients from four institutions who underwent alloHCT for MDS or secondary AML (**Supp.Fig.2A**). We included patients who had a post-transplant BM obtained before clinical evidence of relapse and either subsequently relapsed or had ≥1.5 years of follow-up. After excluding cases with uncertain relapse status (n=10), absence of an available BM specimen within 150 days of transplant (n=5), or no known MDS-associated driver mutation (n=9), 67 patients remained. We further restricted our analysis to samples yielding >25 purified HSC/MPP/LMPPs (n=57) and average sequencing depth >10× (n=56). Of these, 32 (57%) relapsed, whereas 24 (43%) remained progression-free at a median follow-up of 2.2 years. We performed low-input DNA sequencing on purified HSC/MPPs, LMPPs, progenitors, and bulk BM cells. In rare instances of limiting cell numbers, HSC/MPPs and LMPPs were sorted together (lin^-^CD34^+^CD38^-^). Median cell yields per patient were 152 HSC/MPPs, 130 LMPPs, and 1,552 progenitors (**Fig.3A**). UDI-adjusted read depth scaled proportionally with cell input (Spearman ρ = 0.69–0.71, all *P* < 0.001; **Fig.3B**). This relationship was maintained across a broad dynamic range, spanning approximately 16–25,000 sorted cells and 8–660× read depth (5^th^–95^th^ percentiles). Together, these metrics are indicative of accurate PCR duplicate removal and consistent sequencing library quality across sorted cell populations and specimens.

**Figure 3.**
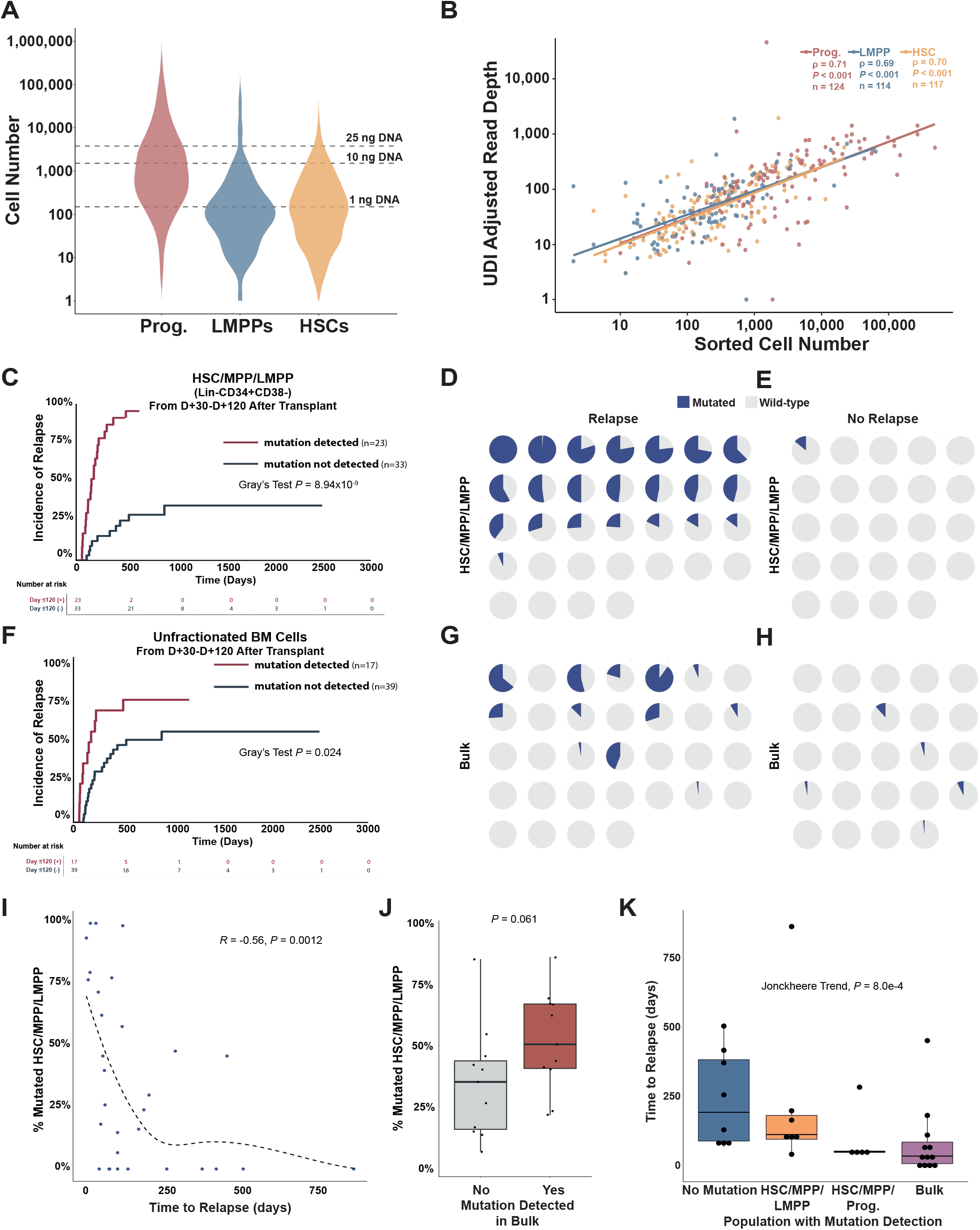
Persistent MDS HSCs identify patients destined to relapse. (**A**) Distribution of sorted cell numbers across restricted progenitor (Prog.), lymphoid-primed multipotent progenitor (LMPP), and hematopoietic stem cell (HSC) populations in the study cohort. Dashed lines indicate the approximate numbers of cells required to yield 1 ng, 10 ng, and 25 ng of genomic DNA. (**B**) Relationship between sorted cell input and unique dual index (UDI)-adjusted read depth across analyzed cell populations. Each point represents an individual sorted cell population, colored according to the respective cell type. Dashed lines indicate the fitted power-law relationship of each corresponding cell type. (**C-H**) Cumulative incidence of relapse and MDS disease burden among all patients in the cohort according to the detection of any persistent pathogenic variant in hematopoietic stem cells (HSCs), multipotent progenitors (MPPs), or lymphoid-primed multipotent progenitor cells (LMPPs; **C-E**) and unfractionated bone marrow (bulk, **F-H**). Statistical significance was assessed using Gray’s test (**C,F**). Pie charts show the relative frequencies of MDS (blue) and wild-type (WT, gray) cells in HSC/MPP/LMPPs (**D-E**) and unfractionated BM (bulk, **G-H**) in patients who relapsed (**D,G**) or did not (**E,H**). (**I**) Correlation between estimated mutated HSC/MPP/LMPP burden and time from testing to clinical relapse. Spearman correlation is shown; the dashed line indicates a LOESS-smoothed trend. (**J**) Estimated MDS burden within HSC/MPP/LMPPs in patients with or without pathogenic variants detected in bulk bone marrow. (**K**) Time from testing to relapse stratified by the most mature population in which residual disease was detected: no mutation detected, or detection of pathogenic variants in HSC/MPP/LMPPs, restricted progenitors, or bulk. Statistical significance was assessed using two-sided Spearman’s rank correlation (**B**), Gray’s test (**C,F**), a two-tailed Student’s *t*-test (**J**), and a Jonckheere trend test (**K**).

We next correlated detection of MDS driver mutations in purified cell populations with relapse (**Supp. Fig.2B, Supp. Table 9**). Detection of mutations in purified HSC/MPP/LMPPs was associated with an increased risk of relapse (22/23 [95.7%] vs. 10/33 [30.3%] when not detected; hazard ratio, 8.80; 95% CI, 3.91–19.8; *P* = 1.5×10^-7^; **Fig. 3C-E**, **Table 1**). The mean interval between detection of mutations in HSC/MPP/LMPPs and clinical relapse was 98 days, with >20% of patients relapsing more than five months after detection. Notably, the only patient with mutations detected in HSC/MPP/LMPPs who did not relapse received 12 months of maintenance therapy on a clinical trial incorporating azacytidine and the novel mutant TP53 re-activating agent APR-246^23^. Detection of mutations in bulk BM was also associated with an increased risk of relapse, but to a lesser degree (70% vs. 51% when not detected; hazard ratio, 2.25; 95% CI, 1.06–4.77; *P* = 0.034; **Fig.3F-H**, **Table 1**). In a competing-risks model including HSC/MPP/LMPPs, restricted progenitors, and bulk BM, detection of mutations in HSC/MPP/LMPPs was strongly associated with relapse (subdistribution hazard ratio, 4.71; 95% CI, 1.56–14.21; *P* = 0.006), whereas mutation detection in bulk BM was not independently associated with relapse (subdistribution hazard ratio, 0.85; 95% CI, 0.36–2.01; *P* = 0.72; **Table 1**). Consistent with the increased discriminatory potential of sequencing HSCs, of the 23 cases with detectable mutations in HSC/MPP/LMPPs, only 11 (48%) had mutations detected in bulk BM despite a median read depth of 476× (**Fig.3D, G**). Importantly, five patients had mutations detected in bulk BM (**Fig. 3H**) but not in HSC/MPP/LMPPs (**Fig. 3E**), and none of these patients relapsed.

**Table 1.** Univariate and joint-model competing risks analyses of association between clinical variables and MRD status with post-transplant relapse. Subdistribution hazard ratios (SHR) and 95% confidence intervals (CI) were calculated using competing-risks regression. Measurable residual disease (MRD) status was assessed by mutation detection in bulk bone marrow (BM), progenitor, and HSC/MPP/LMPP compartments. Abbreviations: MDS-RS, MDS with ring sideroblasts; MDS/MPN, myelodysplastic/myeloproliferative neoplasm; AML-MR, acute myeloid leukemia, myelodysplasia-related; TP53 WT, TP53 wild-type; DTA, DNMT3A/TET2/ASXL1; HMA, hypomethylating agent; Ven, venetoclax; PD, progressive disease; SD, stable disease; CRuni, complete remission with uni-lineage hematologic recovery; CRbi, complete remission with bi-lineage hematologic recovery; CRi, complete remission with incomplete hematologic recovery; CR, complete remission; Flu, fludarabine; Mel, melphalan; Bu, busulfan; Cy, cyclophosphamide; TBI, total body irradiation; RIC, reduced-intensity conditioning; NMA, non-myeloablative conditioning; MAC, myeloablative conditioning; BM, bone marrow; PBSCT, peripheral blood stem cell transplantation; MRD (donor type), matched related donor; MUD, matched unrelated donor; MMRD, mismatched related donor; Haplo, haploidentical donor; Tacro, tacrolimus; MMF, mycophenolate mofetil; PTCy, post-transplant cyclophosphamide; MTX, methotrexate; Siro, sirolimus; CSA, cyclosporine; GVHD, graft-versus-host disease; Aza, azacytidine.

|  | No relapse | Relapse |  | SHR (95% CI) | P-value |
| --- | --- | --- | --- | --- | --- |
| **Clinical characteristics** |  |  |  |  |  |
| Number of patients | 24 | 32 |  |  |  |
| Days from transplant to sequencing, per 10 days | 97 (68–100) | 84 (38–98) | • | 0.95 (0.86–1.05) | 0.34 |
| **WHO diagnosis group** |  |  |  |  |  |
| Other MDS | 8/21 (38.1%) | 3/31 (9.7%) | ◀—•—\| | 0.30 (0.09–0.96) | 0.042 |
| MDS-RS | 1/21 (4.8%) | 1/31 (3.2%) | —•—\| | 1.26 (0.22–7.11) | 0.80 |
| MDS increased blasts | 5/21 (23.8%) | 13/31 (41.9%) | —•—\| | 1.66 (0.82–3.36) | 0.16 |
| MDS-del(5q) | 0/21 (0%) | 1/31 (3.2%) | —•—\| | 6.71 (3.22–14.00) | 3.8e-07 |
| MDS/MPN | 0/21 (0%) | 6/31 (19.4%) | —•—\| | 2.90 (1.52–5.52) | 0.001 |
| AML-MR | 7/21 (33.3%) | 7/31 (22.6%) | —•—\| | 0.61 (0.27–1.37) | 0.23 |
| **TP53 mutation status** |  |  |  |  |  |
| TP53 WT | 20/24 (83.3%) | 18/32 (56.2%) | —•—\| | 0.52 (0.26–1.01) | 0.053 |
| Single TP53 Mutant | 4/24 (16.7%) | 10/32 (31.2%) | —•—\| | 1.57 (0.75–3.28) | 0.23 |
| Bi-allelic TP53 Mutant | 0/24 (0%) | 4/32 (12.5%) | —•—\| | 2.18 (1.21–3.92) | 0.010 |
| **DTA mutation status** |  |  |  |  |  |
| DTA WT | 14/24 (58.3%) | 18/32 (56.2%) | —•—\| | 1.03 (0.52–2.06) | 0.93 |
| DTA Mutant | 10/24 (41.7%) | 14/32 (43.8%) | —•—\| | 0.97 (0.49–1.93) | 0.93 |
| **Pre-HCT treatment** |  |  |  |  |  |
| None | 1/24 (4.2%) | 1/32 (3.1%) | —•—\| | 0.96 (0.13–7.28) | 0.97 |
| HMA | 9/24 (37.5%) | 19/32 (59.4%) | —•—\| | 1.96 (0.97–3.96) | 0.059 |
| HMA/Venetoclax | 4/24 (16.7%) | 9/32 (28.1%) | —•—\| | 1.65 (0.77–3.54) | 0.20 |
| Intensive chemotherapy | 6/24 (25%) | 2/32 (6.2%) | ◀—•—\| | 0.26 (0.07–1.05) | 0.058 |
| Unknown | 4/24 (16.7%) | 1/32 (3.1%) | ◀—•—\| | 0.18 (0.04–0.97) | 0.046 |
| **Response to pre-HCT treatment** |  |  |  |  |  |
| PD | 2/15 (13.3%) | 1/21 (4.8%) | ◀—•—\| | 0.43 (0.06–3.18) | 0.41 |
| SD | 3/15 (20%) | 6/21 (28.6%) | —•—\| | 1.52 (0.60–3.84) | 0.38 |
| CRUni | 4/15 (26.7%) | 0/21 (0%) | —•—\| | 0.00 (0.00–0.00) | 0e+00 |
| CRbi | 1/15 (6.7%) | 2/21 (9.5%) | —•—\| | 1.45 (0.30–7.11) | 0.64 |
| CRi | 1/15 (6.7%) | 1/21 (4.8%) | ◀—•—\| | 1.08 (0.09–12.52) | 0.95 |
| CR | 4/15 (26.7%) | 11/21 (52.4%) | —•—\| | 1.57 (0.69–3.58) | 0.28 |
| **Conditioning regimen** |  |  |  |  |  |
| Flu/Mel | 8/15 (53.3%) | 15/29 (51.7%) | —•—\| | 0.85 (0.42–1.75) | 0.67 |
| Flu/Bu | 3/15 (20%) | 11/29 (37.9%) | —•—\| | 1.91 (0.90–4.07) | 0.092 |
| Flu/Cy/TBI | 3/15 (20%) | 1/29 (3.4%) | ◀—•—\| | 0.24 (0.03–1.96) | 0.18 |
| Bu/Cy | 1/15 (6.7%) | 2/29 (6.9%) | —•—\| | 0.98 (0.27–3.57) | 0.98 |
| **Conditioning intensity** |  |  |  |  |  |
| RIC/NMA | 15/20 (75%) | 17/31 (54.8%) | —•—\| | 0.70 (0.35–1.38) | 0.30 |
| MAC | 5/20 (25%) | 14/31 (45.2%) | —•—\| | 1.43 (0.73–2.82) | 0.30 |
| **Graft source** |  |  |  |  |  |
| BM | 1/20 (5%) | 3/30 (10%) | —•—\| | 1.24 (0.45–3.41) | 0.68 |
| PBSCT | 19/20 (95%) | 27/30 (90%) | —•—\| | 0.81 (0.29–2.22) | 0.68 |
| **Donor type** |  |  |  |  |  |
| MRD | 0/19 (0%) | 5/31 (16.1%) | —•—\| | 2.21 (1.18–4.13) | 0.013 |
| MUD | 13/19 (68.4%) | 25/31 (80.6%) | —•—\| | 1.69 (0.70–4.05) | 0.24 |
| MMRD | 1/19 (5.3%) | 0/31 (0%) | —•—\| | 0.00 (0.00–0.00) | 0e+00 |
| Haplo | 5/19 (26.3%) | 1/31 (3.2%) | ◀—•—\| | 0.16 (0.02–1.26) | 0.081 |
| **GVHD prophylaxis** |  |  |  |  |  |
| Tacro/MMF/PTCy | 16/20 (80%) | 19/30 (63.3%) | —•—\| | 0.61 (0.30–1.25) | 0.18 |
| Tacro/MMF | 1/20 (5%) | 4/30 (13.3%) | —•—\| | 4.41 (1.25–15.54) | 0.021 |
| Tacro/MTX | 0/20 (0%) | 2/30 (6.7%) | —•—\| | 1.99 (0.87–4.55) | 0.10 |
| Tacro/Siro | 1/20 (5%) | 1/30 (3.3%) | —•—\| | 0.50 (0.12–2.01) | 0.33 |
| MMF/CSA | 0/20 (0%) | 4/30 (13.3%) | —•—\| | 2.44 (1.18–5.05) | 0.017 |
| MMF/PTCy/Siro | 2/20 (10%) | 0/30 (0%) | —•—\| | 0.00 (0.00–0.00) | 0e+00 |
| **GVHD outcomes** |  |  |  |  |  |
| Any acute GVHD | 12/20 (60%) | 12/31 (38.7%) | —•—\| | 0.71 (0.34–1.49) | 0.37 |
| Any chronic GVHD | 7/20 (35%) | 9/31 (29%) | —•—\| | 0.72 (0.34–1.51) | 0.38 |
| **Post-transplant maintenance** |  |  |  |  |  |
| None | 14/23 (60.9%) | 19/32 (59.4%) | —•—\| | 1.26 (0.64–2.47) | 0.50 |
| Aza-based | 6/23 (26.1%) | 3/32 (9.4%) | —•—\| | 0.39 (0.12–1.26) | 0.11 |
| Ven/Aza-based | 1/23 (4.3%) | 6/32 (18.8%) | —•—\| | 1.70 (0.92–3.17) | 0.092 |
| Aza/APR-246 | 2/23 (8.7%) | 4/32 (12.5%) | —•—\| | 0.94 (0.39–2.27) | 0.89 |
| **MRD detection, univariate** |  |  |  |  |  |
| Bulk BM MRD+ | 5/24 (20.8%) | 12/32 (37.5%) | —•—\| | 2.25 (1.06–4.77) | 0.034 |
| Progenitor MRD+ | 1/24 (4.2%) | 17/32 (53.1%) | —•—\| | 8.22 (4.11–16.43) | 2.6e-09 |
| HSC/MPP/LMPP MRD+ | 1/24 (4.2%) | 22/32 (68.8%) | —•—\| | 8.80 (3.91–19.81) | 1.5e-07 |
| **MRD detection, joint model** |  |  |  |  |  |
| Bulk MRD+ |  |  | —•—\| | 0.85 (0.36–2.01) | 0.72 |
| Progenitor MRD+ |  |  | —•—\| | 2.95 (1.02–8.48) | 0.045 |
| HSC/MPP/LMPP MRD+ |  |  | —•—\| | 4.71 (1.56–14.21) | 0.006 |

0.1 0.5 1 2 5 10 20 Subdistribution hazard ratio

Thus, persistence of MDS HSCs, rather than mutation detection in bulk BM, is the principal compartment-specific determinant of post-transplant relapse risk. Increasing disease burden within HSC/MPP/LMPPs was strongly associated with a shorter time to relapse (Spearman *R* = −0.56, *P* = 0.0012) (**Fig. 3I**). Patients with mutations detected in bulk BM, as compared with those with mutations exclusively detected in HSC/MPP/LMPPs, had significantly higher disease burden in HSC/MPP/LMPPs (**Fig. 3J**). Accordingly, the interval between mutation detection and clinical relapse decreased progressively as mutations became detectable in more differentiated cells (*P* = 8×10⁻^4^; **Fig. 3K**). These kinetics indicate that relapse is preceded by progressive expansion of mutant MDS HSCs to high frequencies, eroding donor hematopoiesis before clinical relapse; MDS cells become detectable in progenitors and bulk BM only late in this process. Finally, absence of detectable mutations across all cell populations was associated with the longest time to relapse (**Fig. 3K**), whereas detection of mutations in HSC/MPP/LMPPs was strongly predictive of relapse within one year of testing (**Supp.Fig.2C**). This demonstrates that evaluation of HSCs can estimate proximity to relapse and maximize the lead time between detection of MRD and clinical relapse.

### High-resolution HSC chimerism analysis improves the sensitivity of relapse prediction

Among patients without detectable mutations in HSC/MPP/LMPPs, 30% still relapsed, indicating that driver mutation detection alone may miss some cases with residual MDS. Because our sequencing panel captures numerous germline SNVs discordant between donor and host, we reasoned that donor:host chimerism in HSCs could be inferred using such variants to improve detection of host HSCs. To determine whether donor:host chimerism could be accurately quantified from low cell input, we mixed BM samples from two individuals at defined ratios (50:50– 5:95) and performed low-input DNA sequencing on 2,000, 1,000, or 500 sorted cells (**Fig.4A**). Germline variant calling identified 1,345 discordant SNVs, which were aggregated according to genotype discordance (e.g., HOM:WT, HET:WT, HOM:HET) (**Supp.Fig.3**, **Supp. Methods**). Summation of allele counts across all informative SNVs yielded a very high mean effective read depth (∼42,000×) from 500 sorted cells, permitting precise estimates of patient chimerism despite low cell input. Across all mixing ratios and input levels, the inferred chimerism recapitulated the known cell proportions (**Fig. 4B**).

**Figure 4.**
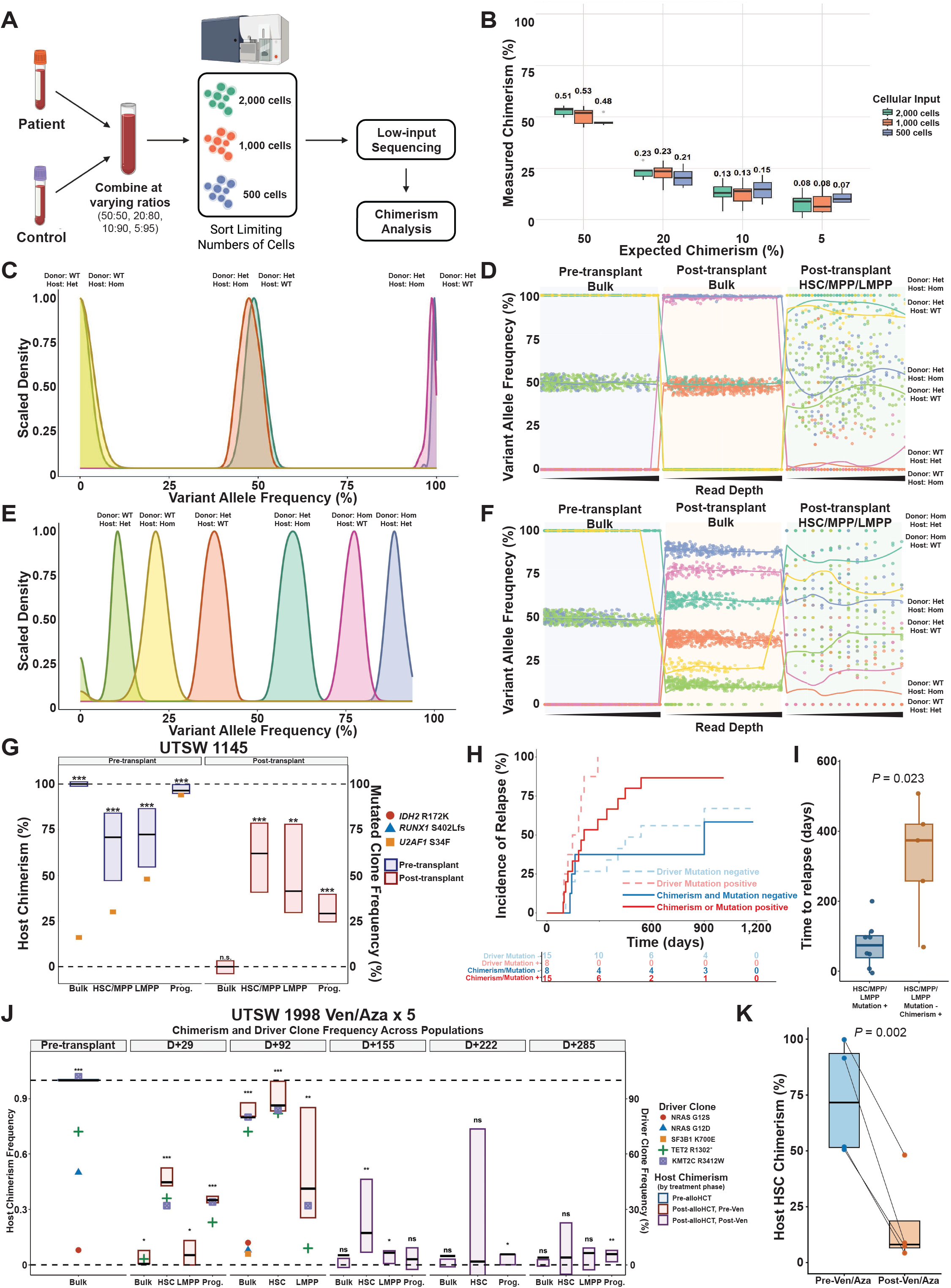
High-resolution HSC chimerism analysis improves the sensitivity of relapse prediction. (**A**) Schematic depicting a mixing experiment to assess the accuracy of chimerism estimation at limiting cell inputs. Cells from two individuals were combined at defined ratios, followed by sorting of decreasing cell numbers (2,000, 1,000, and 500 cells). (**B**) Box plots showing estimated chimerism across mixing ratios, colored by cell input. Distributions reflect chimerism estimates derived from germline variant genotype combinations between individuals (HOM:HET, HOM:WT, HET:HOM, HET:WT, WT:HOM, WT:HET; HOM = homozygous, HET = heterozygous, WT = wild-type). (**C**) Representative density plot of germline variant VAFs for variants with discordant genotypes between donor and host in a bone marrow sample with no detectable host chimerism on clinical testing. (**D**) Scatter plots of germline VAFs versus read depth in unfractionated bone marrow pre- and post-transplant (left) and in post-transplant hematopoietic stem cells (HSCs) (right). Variants are colored by genotype combination (labels on right y-axis). Lines indicate logistic regression fits, with connections linking matched variants across time points and populations. (**E**) Representative density plot of germline variant VAFs from post-transplant bulk bone marrow in a patient with 20% host chimerism on clinical testing. (**F**) Scatter plots of germline variant VAFs versus read depth in unfractionated bone marrow pre- and post-transplant (left) and in post-transplant HSCs (right). (**G**) Low-input sequencing-based chimerism estimates for **UTSW 1145** across unfractionated bone marrow (bulk), HSCs, lymphoid-primed multipotent progenitors (LMPP), and restricted progenitors (Prog.). Boxes indicate 95% confidence intervals of host chimerism, with shading denoting pre-(blue) and post-transplant (red) time points. Black lines represent chimerism estimates derived from variants with host:donor homozygous:wild-type genotypes. Symbols denote the estimated clonal fraction of cells harboring each driver mutation. (**H**) Cumulative incidence of relapse among a subset of patients (n=23) with HSC/MPP/LMPP chimerism data. Dashed lines show relapse based on detection of any pathogenic driver variants in HSC/MPP/LMPPs. Solid lines show relapse based on whether either pathogenic driver variants or host chimerism were present versus whether both pathogenic driver variants and host chimerism were absent in HSC/MPP/LMPPs. (**I**) Time from testing to relapse in patients with detectable pathogenic variants in HSC/MPP/LMPPs versus patients without detectable pathogenic variants but with detectable host chimerism in HSC/MPP/LMPPs. (**J**) Low-input sequencing-based chimerism estimates for **UTSW 1998** across bulk, HSC, LMPP, and Prog. Boxes indicate 95% confidence intervals of host chimerism. The patient received post-transplant venetoclax and azacytidine (Ven/Aza), with shading denoting pre-transplant (blue), post-transplant / pre-Ven/Aza (red), and post-transplant / post-Ven/Aza (purple) time points. Black lines represent chimerism estimates derived from variants with host:donor homozygous:wild-type genotypes. Symbols denote the estimated clonal fraction of cells harboring each driver mutation. (**K**) In four patients who received post-transplant Ven/Aza (shown in **J** and **Supp.Fig.5**), host HSC chimerism prior to the first cycle and after the last cycle of Ven/Aza at time of best clinical response. Statistical significance for elevated chimerism (**G,J**) was assessed using a one-tailed one-sample *t*-test comparing mean host chimerism to zero (*P<0.05, **P<0.01, ***P<0.001; n.s., not significant); for time to relapse (**I**), a two-sided Wilcoxon rank-sum test was used; and for paired pre/post comparisons (**K**), a two-sided paired Student’s *t*-test was used.

We next applied our chimerism analysis to BM specimens from two patients with different known post-transplant chimerism levels as established by clinical short tandem repeat (STR) testing: **UTSW 557**, with 0% host chimerism at D+69, and **UTSW 332**, with 20% host chimerism at D+102 (**Supp. Table 5**). We identified 1,963 and 1,609 putative discordant SNVs, respectively. Chimerism estimates generated from bulk BM closely matched clinical STR testing: **UTSW 557** demonstrated 0% host chimerism (95% CI, −1.4–3.5%; **Fig. 4C-D**), while **UTSW 332** demonstrated 22.0% host chimerism (95% CI, 19.8–24.1%; **Fig.4E-F**). In contrast to measurements from bulk BM, chimerism estimates generated from HSC/MPP/LMPPs from both patients (from 144 and 19 cells, respectively) demonstrated marked enrichment for host cells. In **UTSW 557**, host chimerism increased from 0% in bulk BM to 92.9% in HSC/MPP/LMPPs (95% CI, 88.7–97.0%; **Fig. 4D**). In **UTSW 332**, chimerism rose from 22% in bulk BM to 75.0% in HSC/MPPs (95% CI, 64.1–85.8%; **Fig. 4F**). Thus, bulk BM-based chimerism testing, whether by our SNV-based approach or clinical STR testing, fails to reflect host cell persistence in primitive compartments, likely because rare host cells are obscured by abundant mature donor cells.

We assessed chimerism from HSC/MPP/LMPPs and bulk BM in a subset of 23 patients within our cohort who had paired pre- and post-transplant BM specimens allowing for accurate identification of donor- and host-specific SNVs. In cases with detectable MDS driver mutations, the estimated frequency of clones harboring such mutations demonstrated strong concordance with host chimerism (*R*^2^= 0.86; **Supp.Fig.4A-C**), and the fitted regression line did not differ significantly from the line of equality (intercept = 0.04, 95% CI –0.05–0.14; slope = 1.00, 95% CI 0.77–1.24). Bland-Altman analysis showed minimal bias between overall host chimerism and mutant cell frequency (mean difference +0.045, limit of agreement −0.18–0.27), suggesting that when MDS driver mutations are detectable, host-derived hematopoiesis largely reflects the abundance of MDS cells. In contrast, several cases exhibited detectable host chimerism despite the absence of detectable MDS driver mutations (**Fig. 4G**). This discordance likely reflects both the greater analytical sensitivity of the chimerism assay, arising from the interrogation of hundreds of discordant polymorphic loci rather than individual driver mutations, as well as the persistence of wild-type host HSCs in some cases.

We next tested if detection of persistent host HSC chimerism could improve relapse prediction. We performed chimerism analyses on a subset of 23 patients, of whom 17 relapsed and six remained progression-free. While HSC/MPP/LMPPs harboring MDS driver mutations were detected in only 8 of 17 patients who relapsed (47.1%), elevated host chimerism was detected in HSC/MPP/LMPPs in 13 of 17 patients who relapsed (76.5%; McNemar *P* = 0.0625; **Fig. 4H**). Chimerism detection also preferentially identified patients who relapsed at longer intervals following testing; when host chimerism was present in the absence of detectable driver mutations in HSC/MPP/LMPPs, relapse occurred at a median of 374 days after testing as compared with 97 days when driver mutations were detectable (*P* = 0.023; **Fig. 4I**). Consistent with this, among six patients who relapsed ≥200 days after testing, elevated host chimerism was detected in five (83%), whereas detectable driver mutations were identified in only one (17%). Notably, all patients with detectable driver mutations in HSC/MPP/LMPPs ultimately relapsed, whereas four patients with elevated host HSC/MPP/LMPP chimerism did not (**Fig. 4H**). Thus, host HSC detection after alloHCT increases the sensitivity for relapse prediction at the cost of reduced specificity, likely reflecting persistence of residual wild-type host HSCs in some cases.

Collectively, these findings demonstrate that quantitative donor:host chimerism assessment in purified HSCs is feasible and increases the sensitivity of relapse prediction. More broadly, this approach provides a platform for studying human HSC engraftment dynamics and lineage output following alloHCT.

### Venetoclax and azacytidine can deplete residual MDS HSCs after transplant

Because persistence of host-derived MDS HSCs early after alloHCT was strongly associated with relapse, eradication of persistent MDS HSCs is likely essential to prevent relapse. Although no preemptive therapies have been proven to prevent post-transplant MDS relapse, prophylactic maintenance with venetoclax plus azacytidine (Ven/Aza) appears tolerable in high-risk MDS/AML patients^24^. However, the effect of Ven/Aza on MDS HSCs is unknown. To assess whether Ven/Aza can target alloHCT-resistant MDS HSCs, we performed longitudinal low-input DNA sequencing of FACS-purified HSPC populations from four patients who received Ven/Aza after alloHCT (**UTSW 2111**, **UTSW 1998**, **UTSW 1706**, and **UTSW 2640**). Treatment was initiated in response to early signs of relapse – such as declining donor chimerism or the re-emergence of disease-associated mutations on clinical NGS testing (**Fig.4J**, **Supp.Fig.5A-C**) – at days +281, +114, +193, and +321 post-transplant in the respective cases. We compared pre-Ven/Aza BM specimens with the first BM available at the time of best clinical response (achieved after one cycle for **UTSW 2111**, **1998**, and **1706** and two cycles for **UTSW 2640**). Ven/Aza eradicated detectable MDS driver mutations from HSCs in all but one case (**Supp.Fig.5D**), and it also led to a 74% relative reduction in host HSC chimerism (mean host chimerism decreasing from 73% to 22%; *P* = 0.002; **Fig. 4K**). Notably, **UTSW 2111** and **UTSW 1998** remain without evidence of disease at D+983 and D+1,004, respectively. **UTSW 2640,** who had 5% residual host HSC chimerism at day +373 after receiving Ven/Aza, relapsed at day +660, subsequently underwent intensive induction chemotherapy and a second alloHCT at day +781, and remains disease-free at day +1,072. **UTSW 1706** relapsed with isolated leukemia cutis at D+741, which was treated with radiation, and remains without disease at D+1,199. These results suggest that post-transplant Ven/Aza can reduce or eradicate MDS HSCs, potentially improving the dismal outcomes^25^ of MDS patients who relapse after alloHCT.

#### Selective persistence of immunophenotypically distinct MDS HSCs after transplant

To define the cellular distribution of residual MDS cells after alloHCT, we performed single-cell DNA sequencing (scDNA-seq) on post-transplant specimens from 12 patients and two elderly controls (**Fig.5A**). Cells were stained with a cell surface protein sequencing panel (Abseq^26^) spanning canonical HSPC and mature lineage markers, together with cell surface markers previously implicated in myeloid malignancies^27–37^ (**Supp. Table 6**). To maximize recovery from low-cellularity post-transplant specimens and to differentiate donor from host cells, we developed a multiplexing strategy in which single cells were identified by assigning them scores against patient-specific germline SNV barcodes (**Supp. Methods**). *In silico* mixing of three independently processed samples demonstrated robust demultiplexing performance, resulting in patient assignment with >99% accuracy and a 0.1% misclassification rate (**Supp.Fig.6A-G**). Cell surface protein measurements (Abseq) were normalized and rescaled to enable bivariate visualization mirroring flow cytometry (**Fig.5B**, **Supp.Fig.7A-F, Supp.Fig.8A-C**), allowing for discrimination of cell types (**Supp. Table 7**; broad cell typing, **Fig.5C**; fine cell typing, **Supp.Fig.7G**).

**Figure 5.**
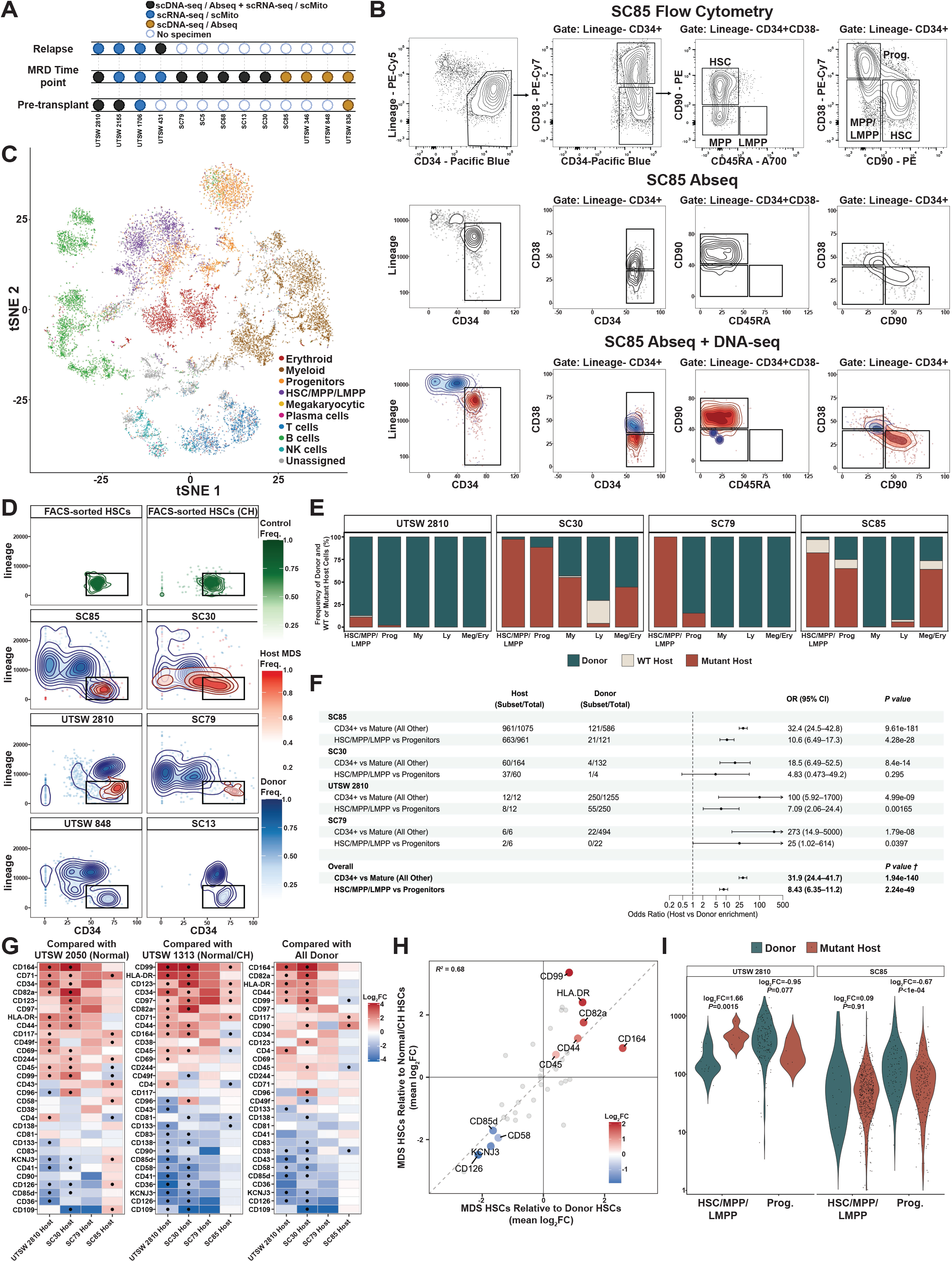
Selective persistence of immunophenotypically distinct MDS HSCs after transplant. (**A**) Schematic overview of single-cell sequencing methods applied to patient specimens spanning treatment time points (pre-transplant, MRD time point, relapse). (**B**) Biaxial gating strategy to identify hematopoietic stem and progenitor cells in a CD34+-enriched bone marrow sample from **SC85** (taken one month prior to relapse) by conventional flow cytometry (top row), by Abseq flow cytometry-like plots (middle row), and by Abseq with donor (blue) and host (red) cell densities shown (bottom row). Hematopoietic stem cells (HSC), multipotent progenitors (MPP), lymphoid-primed multipotent progenitors (LMPP), and restricted progenitors (Prog.) are indicated on flow cytometry plots. (**C**) Cell surface marker-based (Abseq) clustering of 12,880 cells from 14 individuals (12 with MDS, one elderly normal control, and one elderly clonal hematopoiesis control), colored by cell identity as determined by Abseq flow cytometry-like plots. (**D**) Abseq flow cytometry-like plots of lineage versus CD34 expression showing the distribution of donor (blue), host (red), and control hematopoietic stem cells (HSCs, green) obtained in each patient. (**E**) Frequency of donor (green), mutant host (red), and wild-type host cells (tan) among HSC/MPP/LMPP, Prog., mature myeloid (My), mature lymphoid (Ly), and megakaryocytic/erythroid (MegEry) cells. (**F**) Table and forest plot of per-sample odds ratios (±95% CI) for host cell enrichment, with an overall estimate derived from a mixed-effects logistic regression model (random intercept for sample), comparing immature versus mature compartments and HSC/MPP/LMPP versus progenitor populations. (**G-I**) Differential surface protein expression in MDS HSC/MPP/LMPP populations relative to wild-type and donor controls. Heatmaps show median log₂FC across MDS samples. Black dots indicate significant differences (Wilcoxon test, BH-adjusted FDR < 0.05 and |log₂FC| ≥ 1.5). Proteins were filtered for expression in wild-type HSCs (see **Supp.Fig.9A-B**) and selected based on recurrent differential expression across samples. (**H**) Scatter plot shows concordance of mean log₂FC comparing MDS HSCs to normal and clonal hematopoiesis controls or to donor HSCs (Pearson correlation). (**I**) Violin plots showing normalized MHC class II expression across HSC/MPP/LMPP and Prog. cells in **UTSW 2810** and **SC85**. Within cell-type donor versus host comparisons were assessed using Wilcoxon tests with BH correction, with effect sizes reported as median-based log₂ fold-change.

We profiled 12,032 single cells from CD34^+^ cell-enriched early post-alloHCT BM specimens from 12 patients, including six who later relapsed (**Fig.5A**). Multiplexed samples were deconvoluted using patient-specific germline SNVs and mitochondrial variants, enabling accurate assignment of cells to each patient (**Supp.Fig.6H-J**, **Supp. Methods**). To account for age- and clonal hematopoiesis-associated effects, we also analyzed 848 FACS-purified HSCs from two elderly controls (**UTSW 2050** and **UTSW 131**), the latter of whom had *TET2*-mutant clonal hematopoiesis (bulk BM VAF 8.4%; HSC VAF 8.1%). We identified host cells harboring MDS driver mutations in four out of six patients who subsequently relapsed (**Fig.5D**, **Supp. Table 8**), including four out of five who had detectable MDS cells by low-input DNA sequencing (**Supp.Fig.2B, Supp. Table 9**). Our findings support scDNA-seq as another potential MRD detection platform, although low capture efficiency (∼10%) and substantial input requirements (>100,000 cells) were constraints, likely limiting detection of residual disease in two of the six relapsing cases. In **UTSW 848**, only 15 HSC/MPPs were recovered by scDNA-seq, while the estimated MDS cell fraction was ∼10% as determined by low-input DNA sequencing (**Supp. Table 9**). Assuming independent binomial sampling (*P* = 0.10, n = 15), the probability of observing zero malignant cells was (0.9)^15^ ≈ 20.6%. In **SC13**, scDNA-seq capture of just 54 CD34^+^ cells, including no HSCs, precluded meaningful evaluation of the HSPC compartment. Thus, the sensitivity of scDNA-seq may be limited by sparse sampling of hypocellular post-transplant BMs.

In all four patients with residual disease, host MDS cells were markedly enriched within the Lin^-^CD34⁺ compartment relative to mature cells (**Fig.5D-F**, odds ratio [OR], **SC85**: 32.4; **SC30**: 18.5; **UTSW 2810**: 100; **SC79**: 273). Among CD34⁺ cells, residual host MDS cells were further enriched in primitive Lin^-^CD34⁺CD38⁻ HSC/MPP/LMPPs compared to Lin^-^CD34⁺CD38^+^ progenitors in three patients (OR, **SC85**: 10.6; **UTSW 2810**: 7.09; **SC79**: 25.0). A fixed-effect meta-analysis confirmed significant overall enrichment for host MDS cells in CD34⁺ versus mature cells (OR 31.9, 95% CI 24.4–41.7; *P* = 1.94×10⁻^1^^40^) and in HSC/MPP/LMPPs versus progenitors (OR 8.43, 95% CI 6.35–11.2; *P* = 2.24×10⁻^49^). Thus, MDS preferentially persists after alloHCT within the most primitive stem and multipotent progenitor cell populations. These findings help explain why bulk DNA sequencing of unfractionated BM provided no additional predictive value for relapse beyond HSC-directed sequencing.

In three of the four patients with residual disease, donor-derived HSC/MPP/LMPPs remained scarce despite high donor chimerism within mature myeloid and lymphoid cells (**Fig.5E**). In two of the four cases with TP53 mutations (**SC30** and **SC85**), donor chimerism was also reduced in erythroid progenitors, consistent with MDS cells adopting erythroid progenitor-like states, as previously described in *TP53*-mutant AML^38^. Together, these findings indicate that post-alloHCT relapse arises from selective persistence of MDS cells at the apex of the hematopoietic hierarchy, consistent with a stem cell-driven model in which persistent MDS HSCs progressively displace donor hematopoiesis. In *TP53*-mutant disease, this follows a stage- and lineage-specific pattern, with early depletion of donor HSCs and erythroid progenitors preceding overt relapse.

We next asked whether MDS HSCs persisting after alloHCT display distinct immunophenotypic features, focusing on *TP53*-mutant cases given their high risk of relapse and predominance in our cohort. We defined 32 surface proteins robustly detected by Abseq in FACS-purified healthy control HSCs (>50% positivity; scaled expression >40; **Supp.Fig.9A-B**). We then compared their expression on MDS HSCs from four *TP53*-mutant post-transplant cases with expression on healthy control and donor HSCs (**Fig.5G**). Differential expression analysis revealed a consistent immunophenotypic shift in MDS HSCs across cases, characterized by increased expression of CD164, CD71, CD34, CD82a, HLA-DR, CD44, and CD99, and reduced expression of CD109, CD126, CD85d, CD41, and KCNJ3. These changes were concordant across comparisons with matched donor HSCs (*R*² = 0.68; **Fig.5H**), as well as HSCs from both healthy controls with or without clonal hematopoiesis (*R*² = 0.68; **Supp.Fig.9C**). These results support a disease-intrinsic phenotype as opposed to a general response of HSCs to the post-transplant microenvironment. In addition to these shared changes, each patient displayed unique aberrancies previously reported in MDS or AML, including upregulation of CD56 (**SC85**), CD123 (**SC30**), and CD25 (**UTSW 2810**). In each case, biaxial gating of CD99 with the patient-specific markers distinguished mutant HSPCs from donor-derived and age-matched wild-type cells (**Supp.Fig.9D-F**). Together, these findings define a conserved *TP53*-mutant HSC immunophenotypic program, and they also suggest that optimal prospective isolation may require both shared and patient-specific markers.

Given prior studies in AML implicating downregulation of major histocompatibility complex class II genes (MHC-II) as a mechanism of post-alloHCT immune escape^39,40^, we compared HLA-DR expression in MDS and donor HSPCs by Abseq. At the MRD time point, two cases contained sufficient residual host and donor HSPCs to make such a comparison (**Fig.5I**). In **UTSW 2810**, host MDS HSC/MPP/LMPPs exhibited increased HLA-DR relative to donor HSC/MPP/LMPPs (log2FC = 1.66, Benjamini–Hochberg [BH]-adjusted *P* = 0.0015), while progenitors exhibited no significant differences (log2FC = −0.95, BH-adjusted *P* = 0.077). In **SC85**, host MDS HSCs showed no difference in HLA-DR expression relative to donor HSCs (log2FC = 0.09, BH-adjusted *P* = 0.91), whereas host MDS progenitors exhibited significantly reduced HLA-DR expression relative to donor progenitors (log2FC = −0.67, BH-adjusted *P* < 1.0×10⁻^4^). Thus, unlike in AML, MDS HSCs did not exhibit reduced MHC-II expression following alloHCT, supporting a differentiation stage-specific model of post-transplant MHC-II regulation.

### Molecular characteristics of MDS HSCs persisting after transplant

To determine whether MDS HSCs persisting after alloHCT exhibit shared molecular features, we performed single-cell RNA sequencing (scRNA-seq) with paired mitochondrial transcriptome profiling. This enabled distinction of donor from host cells using germline mitochondrial variants and delineation of disease subclones as previously described^41,42^. BM specimens were multiplexed using germline mitochondrial variants, analogous to our scDNA-seq approach (**Supp.Fig.10**, **Supp.Methods**). We profiled 21,906 cells from serial CD34^+^ cell-enriched BM specimens obtained from five patients with residual MDS cells detected prior to post-transplant relapse (**Fig.5A**). Pre-transplant, MRD time point, and relapse specimens were available for three patients (**UTSW 2810**, **UTSW 2155**, and **UTSW 1706**), MRD time point and relapse specimens for one patient (**UTSW 431**), and an MRD time point specimen for one patient (**SC79**).

Residual host cells consistent with persistent MDS were detected at the MRD time point in all cases, albeit at low frequencies (**Fig.6A**). We used somatic mitochondrial variants to define MDS subclones and trace their evolution over the course of alloHCT in four patients (**Fig.6B**, **Supp.Fig.11A**). In **UTSW 431**, characterized by t(1;3)(p36.1;q21) and *FLT3*-ITD, progression from MRD to secondary AML occurred without new driver mutations, but mitochondrial variants (ChrM:3643 G>C, ChrM:9591 G>C) identified an expanding LMPP-like subclone with AML and stem cell programs, indicating progression via expansion of leukemia stem cell-like populations driven by pre-existing lesions (**Supp.Fig.11B**). In the pre-transplant biospecimen from **UTSW 1706**, mitochondrial variants distinguished MDS HSCs from wild-type HSCs, identifying highly distinct features (**Supp.Fig.11C-F**) such as downregulation of genes encoding AP-1 transcription factors and ribosomal proteins, as well as dysregulation of long non-coding RNAs (lncRNAs) and regulators of venetoclax sensitivity. We found that relapse was oligoclonal in all cases, with most dominant relapse subclones detectable before transplant, as has been described^43^, and in three of four patients, already present at the MRD time point (**Fig.6B**, **Supp.Fig.11A**).

**Figure 6.**
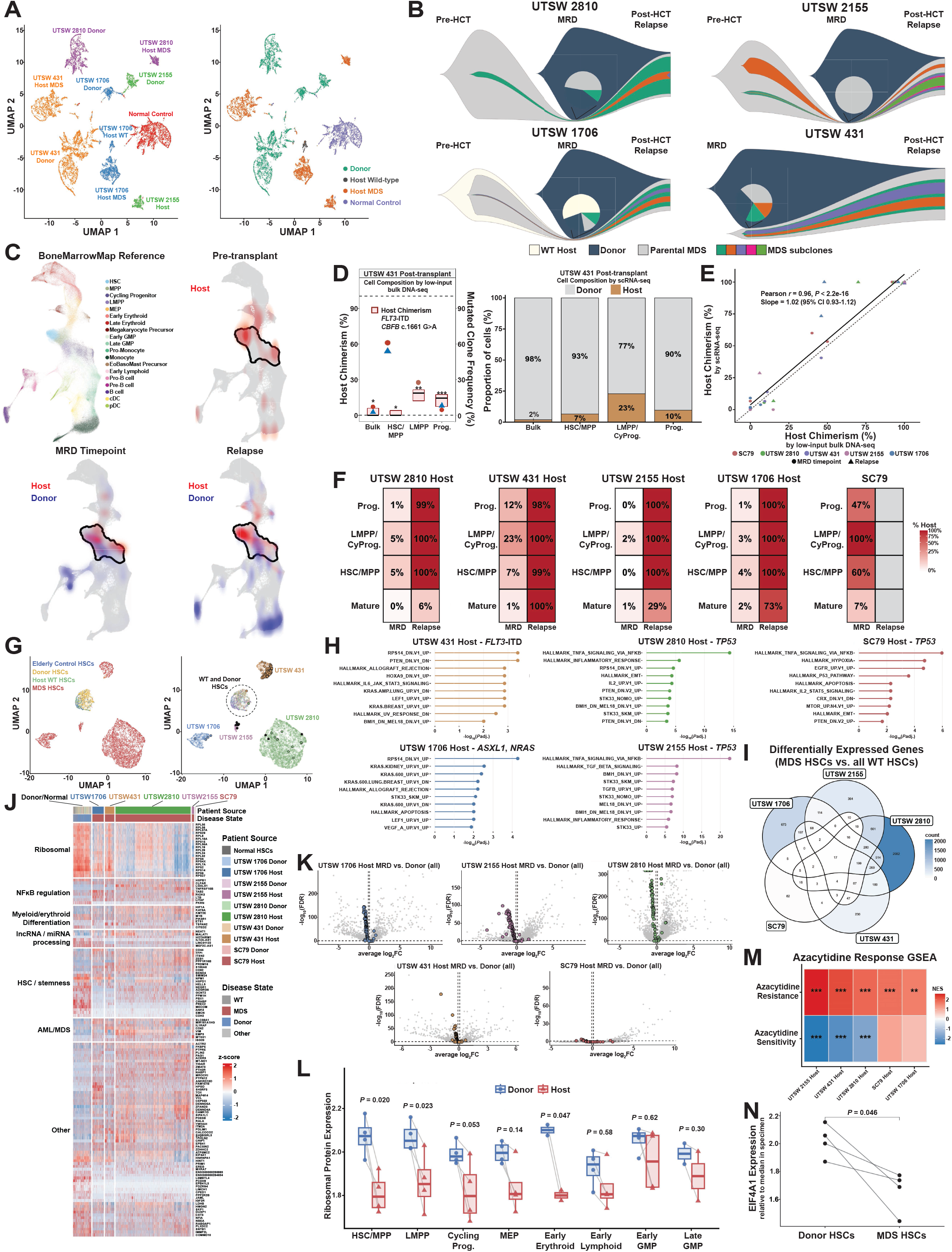
Molecular characteristics of MDS HSCs persisting after transplant. (**A**) UMAP projection of 10,639 cells from five individuals (four MDS, one healthy control), clustered based on mitochondrial variant allelic frequencies. Cells are colored by patient origin (left) and by donor versus host identity (right). (**B**) Fish plots depicting longitudinal changes in donor (dark blue) and host (gray) contributions across pre-transplant, measurable residual disease (MRD), and relapse time points for each patient. Mitochondrially defined subclones are shown as different colors nested within the host compartment. Pie charts show subclonal composition at the MRD time point. (**C**) Kernel density estimates of donor (blue) and host (red) cells projected onto a reference map of hematopoietic cells (BoneMarrowMap) for representative patient **UTSW 2810** across pre-HCT, MRD, and relapse time points. (**D**) Quantification of host chimerism in post-transplant bone marrow at the MRD time point (**UTSW 431**), measured by low-input DNA sequencing (left) and scRNA-seq (right). Boxplots represent 95% confidence intervals. Black lines indicate inferred chimerism from host-versus donor-specific variants. Points denote the frequency of cells with each variant (*FLT3*-ITD, *CBFB*). (**E**) Correlation between host chimerism estimates derived from low-input DNA sequencing and scRNA-seq across HSPC populations. Colors and shapes represent patient cell populations and time points, respectively. Pearson correlation and regression slope are shown. (**F**) Heatmaps showing the proportion of host-derived cells across hematopoietic compartments (HSC/MPP, LMPP and cycling progenitors [CyProg.], lineage-restricted progenitors [Prog.], and mature cells) at MRD and relapse time points for each patient. (**G**) UMAP of HSC/MPP/LMPP populations stratified by cNMF-derived transcriptional modules. Cells are colored by origin and disease state (left) and by patient with MRD time point cells highlighted (right). (**H**) Enrichment of Hallmark and C6 MSigDB gene sets in MDS HSC/MPP/LMPPs relative to WT HSC/MPP/LMPPs for each patient. (**I**) Venn diagram summarizing overlap of significantly differentially expressed genes (adj. *P*-value < 0.05, log₂FC > 0.25) in MDS HSC/MPP/LMPPs compared to WT HSC/MPP/LMPPs across patients. (**J**) Heatmap of scaled expression for genes dysregulated in ≥4 of 5 MDS cases relative to WT HSC/MPP/LMPPs. Columns represent single cells grouped by patient; genes are organized by functional categories. (**K**) Volcano plots of differential gene expression between MDS and WT HSC/MPP/LMPPs for each patient, with ribosomal genes highlighted. (**L**) Boxplots of median ribosomal gene module scores (RPS/RPL/RPLP genes) in matched donor (blue) and host/MDS (red) samples across hematopoietic populations. Points represent patient-level medians, with lines connecting matched donor-host pairs. *P*-values were determined by a paired two-sided *t*-test within each population. (**M**) Gene set enrichment analysis (GSEA) of azacytidine response and non-response gene signatures in host-derived HSPCs versus WT donor controls; normalized enrichment scores (NES) shown, with significance denoted by adjusted P values (*P<0.05, **P<0.01, ***P<0.001). (**N**) Paired comparison of mean EIF4A1 expression in donor versus MDS HSC/MPP and LMPP cells. Each point represents the patient-level mean, with lines connecting matched donor–MDS pairs. *P*-values were calculated using a two-sided paired *t*-test.

Cells were projected onto the BoneMarrowMap reference atlas^38^, derived from comparable scRNA-seq datasets and validated against immunophenotypically-defined hematopoietic populations (**Fig.6C**, **Supp.Fig.12A-B**). Host-cell frequencies across transcriptionally-defined populations closely tracked with disease burden estimates derived from low-input DNA sequencing of FACS-purified populations (Pearson *r* = 0.96, P < 2.2 × 10⁻¹⁶; **Fig.6D-E**). At the MRD time points, host cells were most enriched within HSC/MPP/LMPPs (median 3.61-fold enrichment, 95% CI 1.67–6.93; **Fig.6F**, **Supp.Fig.12C-D**), and in four of the five cases host cells comprised less than 2% of mature cells. At relapse, host cells increased in frequency to comprise nearly all CD34⁺ HSPCs, also becoming detectable in mature cells (**Fig.6F**, **Supp.Fig.12D**), consistent with dropout of normal donor hematopoiesis. These results closely mirror our immunophenotypic data from Abseq (**Fig.5C-F**), independently demonstrating that MDS selectively persists after alloHCT in transcriptionally-defined HSCs that expand to high frequencies at the time of relapse to erode normal donor hematopoiesis.

To identify transcriptional programs associated with transplant-refractory MDS HSCs, we applied consensus non-negative matrix factorization (cNMF) to HSC/MPP/LMPPs from all five patients together with age-matched healthy control HSPCs (**Fig.6G**, **Supp.Fig.13A-B**). MDS cells clustered by patient, revealing substantial interpatient transcriptional heterogeneity. Donor-derived cells clustered tightly with healthy controls and showed little evidence of interpatient heterogeneity, suggesting little effect of the post-transplant microenvironment on the transcriptional profiles of these cells. MDS cells present at MRD and relapse time points from the same patient clustered together, indicating that the transcriptional state observed at relapse was already established early after alloHCT.

In line with the known heterogeneity of MDS, transcriptional programs varied across patients and reflected underlying driver mutations (**Fig.6H**, **Supp.Fig.13C**). For example, *TP53*-mutant cases demonstrated enrichment of TNF-α signaling via NF-κB^44^, whereas cases harboring mutations in signaling effectors (*NRAS* Q61R and *FLT3*-ITD) showed activation of RAS-associated programs, as well as STAT5-associated programs in the *FLT3*-ITD case. We identified a subset of 192 genes highly specific to individual MDS cases, as defined by expression in >50% of cells from one patient and absence in >75% of cells from any other individual (**Supp.Fig.13D**). Expression of patient-specific transcripts was maintained from HSCs through multiple hematopoietic progenitor populations, indicating that differentiated MDS cells retain distinct transcriptional programs established in HSCs (**Supp.Fig.13E**). Patient-specific transcripts included several lncRNAs, such as *LINC01819*, *LINC02476*, *LINC00355*, *TEX41*, and *LINC01619* (**Supp.Fig.13F**). Additionally, in **UTSW 1706** we found that such lncRNAs were specific to MDS cells in a non-transplant setting (**Supp.Fig.11E**).

Despite marked interpatient heterogeneity, we identified 170 genes that were recurrently dysregulated in HSPCs from at least four out of the five patients relative to healthy control HSPCs (**Fig.6I**). These were enriched for genes associated with NF-κB regulation, myeloid/erythroid differentiation, maintenance of HSC function, and AML/MDS (**Fig.6J**). The most prominent shared program was a coordinated reduction in ribosomal protein gene expression (**Fig.6K**). Ribosomal insufficiency is a well-established feature of specific MDS subtypes, including del(5q)^45^ and *DDX41*- or *RUNX1*-mutant disease^46,47^, as well as BM failure syndromes such as Diamond-Blackfan anemia^48^, but has not previously been linked to the mutational profiles observed in our patients. The reduction in ribosomal protein gene expression was not dependent on cell-cycle state (**Supp.Fig.14A**). It showed marked differentiation-stage specificity, with a paired analysis against donor cells demonstrating the strongest downregulation in HSC/MPPs (**Fig.6L**). In the pre-transplant setting, MDS HSCs also exhibited a differentiation stage-specific decrease in ribosomal protein gene expression compared with wild-type HSCs (**Supp.Fig.14B**). Restriction of this ribosomal insufficiency program to primitive HSC/MPPs may explain why it has gone unrecognized in most prior studies of MDS. Together, these findings identify coordinated ribosomal gene downregulation as a shared feature of persistent MDS HSCs, suggesting that alloHCT may select for HSCs with lower protein synthesis capacity.

We next evaluated MDS HSCs for inflammatory gene signatures upregulated in wild-type HSCs in patients with clonal hematopoiesis. Recent studies indicate that such inflammation is selectively suppressed in mutant HSCs relative to wild-type HSCs in clonal hematopoiesis, which may serve as a mechanism for their clonal selection^18^. MDS HSCs exhibited an increase in these signatures compared with donor HSCs (**Supp.Fig.14C**). This is consistent with the fact that donor HSCs are derived from young healthy individuals and thus not previously exposed to the inflamed BM microenvironment of clonal hematopoiesis or MDS. Consistent with this, MDS HSCs also exhibited enrichment of aged HSC gene signatures (**Supp.Fig.14C**). These results suggest that MDS HSCs may not gain a clonal advantage after transplant by suppressing inflammation relative to donor cells.

MDS HSCs also exhibited shared transcriptional programs linked to therapeutic response. Consistent with the resistance of MDS HSCs to non-transplant therapies^5,6^ and uniform treatment exposure before alloHCT, MDS HSCs from every patient were enriched for azacytidine resistance signatures^49^ (**Fig.6M**), suggesting the presence of shared mechanisms of therapy resistance, and in line with the minimal benefit of post-transplant azacytidine^50^. MDS HSCs also upregulated the lncRNAs *MALAT1* and *NEAT1*, which can promote BCL2 expression^51,52^ (**Supp.Fig.14D**). This may reflect a dependence on BCL2 consistent with the robust responses to venetoclax we observed (**Fig.4J**, **Supp.Fig.5**). We also observed markedly reduced expression of *EIF4A1* (**Fig.6N**), which encodes a translation initiation factor whose inhibition synergizes with venetoclax^53^. Finally, we observed upregulation of potential therapeutic targets such as *KAT6A*^54^ and *HIF1A*^55^ (**Supp.Fig.14E**), the latter of which has been broadly implicated in MDS pathogenesis^56^. Thus, MDS HSCs persisting after transplant exhibit shared features that may explain resistance to azacytidine and sensitivity to venetoclax, while identifying candidate therapeutic vulnerabilities.

## Discussion

Evaluating MDS HSCs is critical for assessing therapeutic response^5,6^ and understanding disease biology^1–4^, but existing approaches either lack broad applicability or require patient-specific customization. Fluorescence *in situ* hybridization is applicable to only 43% of MDS cases, whereas 91% harbor NGS-detectable mutations^57^; ddPCR requires patient-specific assays and cannot detect *de novo* variants^14^; and single-cell approaches remain constrained by input requirements, capture efficiency, and cost. Our ultra-low-input error-corrected DNA sequencing assay addresses this gap, detecting the full spectrum of MDS-associated mutations in rare cells and generating interpretable data from 98% of BM specimens yielding more than 25 HSC/MPP/LMPPs. It requires no patient-specific customization and enables *de novo* variant discovery, including mutations absent from clinical panels, unmasked by post-transplant clonal expansion^43^, or restricted to immature cells. Finally, this assay provides a broadly applicable platform for genomic interrogation of rare purified cell populations across a wide range of biologically and clinically relevant contexts.

Our analysis demonstrates selective MDS HSC persistence after transplant, with orthogonal confirmation by scDNA-seq/Abseq immunophenotyping and scRNA-seq transcriptional profiling. Persistence was nearly invariably associated with relapse, whereas unfractionated BM sequencing added no predictive information and mutations restricted to more mature cells were not associated with relapse. Thus, the cellular compartment harboring residual mutations determines their prognostic significance, providing a biological explanation for the high false-positive rates and limited positive predictive value reported for studies using ultra-deep sequencing^58^ or chimerism analyses^59,60^ of unfractionated BM. Prior ddPCR-based work detected persistent MDS HSCs before relapse^14^. Our study defines the biological context in which these residual mutations are clinically meaningful by demonstrating that they predict relapse only when they persist within the HSC compartment. Because isolated *DNMT3A*, *TET2*, and *ASXL1* (DTA) mutations might reflect clonal hematopoiesis, the prior study required another corroborating mutation^14^. We included isolated DTA mutations without loss of specificity, substantially broadening eligibility for HSC-based disease monitoring and suggesting host-derived clonal hematopoiesis is uncommon after alloHCT.

An additional strength of our ultra-low-input DNA sequencing approach is the ability to precisely quantify donor:host HSC chimerism after alloHCT without patient-specific customization, using hundreds to thousands of informative SNVs to overcome the limited precision of clinical STR assays^61^. Estimates were consistent across donor types and independently supported by mtDNA variants in our single-cell data. Detection of residual host HSCs improved sensitivity for relapse prediction compared with mutation-based approaches, albeit with reduced specificity, consistent with persistence of wild-type host HSCs in some cases. Host chimerism closely paralleled driver mutation burden in most cases, suggesting preferential persistence of MDS HSCs after alloHCT, although paired pre- and post-transplant analyses will be required to determine the extent of this selective persistence. HSC chimerism analyses also extend MRD assessment to patients lacking known driver mutations, where elevated host chimerism may serve as a surrogate for residual disease. Finally, this approach provides a broadly applicable platform for studying human HSC engraftment dynamics and lineage output after transplantation, enabling direct investigation of human HSC biology *in vivo* without the need for genetic labeling^62^.

We present a mutation-resolved single-cell multi-omics approach in which multiplexing specimens for scDNA- and scRNA-sequencing overcomes input constraints and enables efficient characterization of rare cell populations. We leveraged nuclear and mitochondrial DNA variants captured across each modality to resolve patient identity and MDS subclonal structure, enabling direct comparison of aberrant HSCs with their wild-type counterparts and delineation of molecularly distinct subclones. MDS HSCs exhibited substantial patient-specific variation in surface protein abundance, lncRNA expression, and transcriptional programs, consistent with the genetic heterogeneity of MDS and the influence of underlying driver mutations. For example, *TP53*-mutant disease was associated with NF-κB signaling, consistent with its role in LSC survival^63^ and repression by TP53^44^. Despite this diversity, recurrent dysregulation of 170 genes defined a shared transcriptional architecture.

A predominant shared feature was a coordinated reduction in ribosomal protein gene expression, most pronounced in HSC/MPP/LMPPs and absent in myeloid progenitors. Since normal HSCs require low protein synthesis rates^64^ and are sensitive to proteotoxic stress^65^, ribosomal insufficiency may confer a selective advantage to HSCs by reducing proteotoxic stress during hematopoietic reconstitution. Furthermore, ribosomal insufficiency is a general hallmark of MDS^66^ that is particularly pronounced in MDS HSCs^66,67^. In such pathologic states of ribosomal insufficiency, clonally selected HSCs may exhibit impaired differentiation^46^ due to the high protein synthesis rates required in restricted progenitor cells^64^. Together, these findings suggest that alloHCT selects for ribosomal insufficiency as a shared feature of persistent MDS HSCs, and that selection for this phenotype may contribute to the marked impairment in differentiation observed after transplant.

In the two evaluable cases, MDS HSCs did not downregulate MHC-II after alloHCT, unlike in AML^39,40^, although progenitors in one case did show downregulation. This may reflect cell-of-origin context, as AML can arise from progenitors^20,68^ and MDS from HSCs^1–4^, suggesting stage-specific MHC-II regulation. Notably, MHC-II expression has been associated with promotion of a stress-resistant quiescent state in HSCs^69^. Unlike in clonal hematopoiesis, where mutant HSCs gain fitness by attenuating inflammatory responses^18,70^, persistent MDS HSCs exhibited increased inflammatory signatures relative to donor HSCs. Thus, intrinsic MDS HSC properties, rather than MHC-II downregulation or inflammation suppression, may underlie post-transplant persistence.

Our data identify persistent MDS HSCs as the critical target for preventing post-transplant relapse. The high specificity of MDS HSC detection and the dismal prognosis of post-transplant relapse^25^ provide a strong rationale for preemptive intervention trials. In contrast, the limited specificity of mutation detection in unfractionated BM suggests that therapeutic decisions based solely on such testing may expose patients unlikely to relapse to unnecessary post-transplant therapies. Our ultra-low-input DNA sequencing assay enables identification of patients with persistent MDS HSCs in whom clinical trials could evaluate Ven/Aza^24^, donor lymphocyte infusion, or a second alloHCT. In such patients, therapy could be initiated when MDS HSC burden remains low and donor hematopoiesis remains sufficient to support treatment tolerance. Our assay also provides a stem cell-based biomarker for longitudinal monitoring, enabling early treatment switching in non-responders and de-escalation after MDS HSC eradication to limit toxicity.

Although the phase 3 VERONA trial testing Ven/Aza in MDS patients in the non-transplant setting did not demonstrate an overall survival benefit^71^, recent studies have found an unusual degree of response to venetoclax-based regimens in the post-transplant setting^24,72,73^. The effect of Ven/Aza on MDS HSCs has not previously been directly assessed in any context. Despite enhanced azacytidine resistance signatures, we found that Ven/Aza reduced or eliminated detectable MDS HSCs in all longitudinally evaluated cases, enabling re-expansion of donor-derived hematopoiesis. Our multi-omics analyses identify candidate mediators of venetoclax sensitivity such as *MALAT1*^51^, *NEAT1*^52^, and *EIF4A1*^53^, which may serve as biomarkers to predict response and guide therapy. We also identified upregulation of potential therapeutic targets in MDS HSCs, including *KAT6A* and *HIF1A*, for which clinical-stage inhibitors are available^54,55^. Finally, ribosomal insufficiency may sensitize MDS HSCs to inhibitors of ribosome biogenesis or translation.

Despite its high specificity, HSC mutation detection had imperfect sensitivity, with 30.3% of patients without detectable HSC mutations subsequently relapsing. This likely reflects limited detection of low-VAF mutations in the small numbers of HSCs recovered from archived cryopreserved specimens. Sensitivity may improve with prospectively collected fresh samples yielding more HSCs. Nevertheless, relapse occurred much later in patients without detectable mutations or with host HSC chimerism alone (**Fig.3K**, **Fig.4I**), suggesting that serial BM assessments in these patients could provide additional opportunities to detect MDS HSCs before overt clinical relapse. Prospective studies are needed to evaluate these approaches, as well as to determine whether circulating HSCs can be used for less invasive disease monitoring.

Limitations of our study include its retrospective design and cohort heterogeneity; prospective validation in larger risk-and treatment-stratified cohorts is needed. Paired pre- and post-transplant analyses are required to define the impact of pre-transplant therapies, conditioning, and baseline MDS HSC burden on relapse risk. Further mechanistic studies are needed to define the drivers of ribosomal insufficiency, the functional impact of aberrant lncRNAs, and the contribution of NF-κB signaling to transplant resistance in *TP53*-mutant disease – questions that the HSC-resolved sequencing approaches described here now allow us to investigate directly in primary patient HSCs. Validation of MDS HSC-specific surface markers across disease subtypes promises to enable flow cytometry-based detection of MDS HSCs for diagnostic studies, as well as their prospective isolation for functional studies. Together, our work establishes the feasibility and clinical relevance of universal, patient-agnostic MDS HSC detection after alloHCT while providing a robust platform to investigate and therapeutically target relapse-driving MDS HSCs.

## Methods

### Study Design and Patient Samples

Institutional review board approval for biospecimen accrual was obtained at the University of Texas Southwestern Medical Center, the Cleveland Clinic, Moffitt Cancer Center, and Memorial Sloan Kettering Cancer Center (MSKCC). Studies were conducted according to the principles embodied in the Declaration of Helsinki, and all biospecimens were obtained with informed consent. De-identified biospecimens and clinical data were collected from 91 patients (n=37 from UTSW, n=26 from the Cleveland Clinic, n=13 from Moffitt Cancer Center, and n=15 from MSKCC) who underwent alloHCT for MDS or acute myeloid leukemia, myelodysplasia-related (AML-MR). Clinical eligibility required availability of a BM specimen obtained between D+30 and D+150, before relapse. Patients were otherwise not excluded based on treatment regimen or any other clinical risk characteristics. Of these, 81 had known relapse status, and 76 of those patients had available BM biospecimens obtained between D+30 and D+150 after transplant. Of these 76 patients, 67 had known driver mutations based on clinical sequencing. Ten patients lacked sufficient sorted cells to meet the >25-cell criterion, and one failed the >10× mean read-depth criterion, leaving 56 patients in the primary analysis. Of these, 23 had paired pre- and post-transplant specimens for chimerism analyses. Four patients were evaluated longitudinally during post-transplant Ven/Aza therapy.

### Relapse Criteria and Endpoints

Relapse was defined as the presence of any of the following: >5% myeloblasts in the BM, reemergence of pre-transplant cytogenetic abnormalities, or clinical intervention due to evidence of systemic relapse, such as loss of donor chimerism or recurrence of pre-transplant morphological features. Isolated extramedullary relapse (including the central nervous system) was not counted as relapse. Patients receiving prophylactic post-transplant azacytidine (n=14) were included. As biospecimens were de-identified, treating clinicians were not aware of the sequencing data presented in this manuscript, and these data were not used to make treatment decisions.

### Purification of HSPCs

Hematopoietic stem and progenitor cells were identified using the following antibodies: anti-CD34 (581), anti-CD90 (5E10), and anti-CD45RA (HI100) from BioLegend, and anti-CD38 (HIT2) from eBioscience. We additionally used a lineage cocktail containing antibodies against CD2 (RPA-2.10), CD3 (HIT3a), CD4 (RPA-T4), CD7 (M-T701), CD8 (RPA-T8), CD10 (HI10a), CD11b (ICRF44), CD14 (TuK4), CD19 (CC2C6), CD20 (2H7), GPA (HIR2), and CD56 (B159), all from BD Biosciences. Cells were purified using a FACSAria II flow cytometer, and data were analyzed using FlowJo v10 software (Tree Star). Stem and progenitor cell populations were sorted using fluorescence-activated cell sorting (FACS), employing established, functionally validated gating strategies, as shown in **Supp. Table 1**, with representative flow cytometry gates shown in **Fig.2A-D** and **Fig.5B**. CD38 expression thresholds were determined by evaluating expression of CD90, which is typically absent or present at very low levels on CD38⁺ restricted progenitors.

### Low-Input Targeted DNA Sequencing

Targeted DNA sequencing of 261 genes associated with myeloid malignancies (**Supp. Table 2**) was performed using unique dual index-corrected next-generation sequencing. Library preparation was conducted using the xGen cfDNA and FFPE and xGen Hybrid Capture Core Reagent kits (IDT), following the manufacturer’s guidelines. See **Supplementary Methods** for sequencing and analysis methods.

### Statistical Analysis

The primary endpoint was cumulative incidence of relapse. Clinical, cytogenetic, and molecular characteristics of relapsed and non-relapsed patients were compared (**Table 1**). Relapse incidence was assessed using the Gray model for competing risks. Host chimerism was considered detectable when the 95% confidence interval for donor chimerism excluded 100%. Statistical significance was defined as a two-sided *P*-value ≤ 0.05.

### Mission Bio Tapestri single-cell DNA + Abseq library preparation and sequencing

Single-cell DNA sequencing (scDNA-seq) and cell surface protein (Abseq) profiling were performed using the Mission Bio Tapestri platform. Live cells were FACS-purified from Ficoll-separated BM mononuclear cell preparations, stained with a custom panel of oligonucleotide-conjugated antibodies (**Supp. Table 6**), washed, and loaded onto the Tapestri instrument for droplet-based single-cell encapsulation and barcoding at a target concentration of 3,500 cells/µL. See **Supplementary Methods** for sequencing and analysis methods.

## Data availability statement

All single-cell multi-omics data are available at GEO under accession number GSE345192. Processed data and source data supporting the findings of this study, as well as custom code used for data processing and analysis, are available from the corresponding author upon request.

## Acknowledgements

This work was supported by grants from the National Cancer Institute (K08CA194275, R37CA273232), the Department of Defense (W81XWH-22-1-0857), the American Society of Hematology (Scholar Award), the Edward P. Evans Foundation (Discovery Research Grant), the Henry & Marilyn Taub Foundation (Pilot Award in MDS Research), the Dresner Foundation (MDS Research Fund), and the Cancer Prevention and Research Institute of Texas (RR180046, RP250208), as well as philanthropic funding from Irwin Gordon and Malcolm Louden to S.S.C. B.K. was supported by grants from the National Cancer Institute (R37CA273232-02S1) and the National Heart, Lung, and Blood Institute (F30HL178281).

## Authorship contributions

S.S.C., B.K., and R.T. (Tamari) conceived the project and S.S.C. supervised all aspects of the project. B.K. designed and performed all experiments and analyzed the resulting data. Y.H., T.T., A.S., and P.C. contributed to cell sorting experiments. Y.H. and P.C. contributed to low-input DNA sequencing library preparation. Y.J. generated single-cell RNA-sequencing libraries. R.T. (Tamari), C.G., V.V., D.A.S., A.M. (Mishra), and J.M. provided curated patient specimens and accompanying de-identified clinical data from MSKCC, Cleveland Clinic, and Moffitt Cancer Center. Y.M., R.C., C.J., V.I., A.M. (Marinos), P.P., M.V., and J.A. provided de-identified clinical data from UTSW. R.T. (Tachibana) contributed to development of the chimerism analysis pipeline. Z.B., A.M. (McLemore), A.C., J.Y., B.D.A.H., and R.V. assisted with biospecimen collection, processing, and annotation. B.K. and S.S.C. wrote the manuscript with input from all authors. Current affiliations: Y.H., The University of Texas at Dallas, Dallas, TX; P.P., Servier BioInnovation, Boston, MA; M.V., Actinium Pharmaceuticals, New York, NY; J.A., Department of Hematology and Hematopoietic Cell Transplantation, City of Hope, Duarte, CA.

