## Supplemental Figures 1-14 for "Persistent Malignant Stem Cells Drive Relapse After Transplantation in the Myelodysplastic Neoplasms"

**Supplementary Data**

#### Supplementary Figure 1

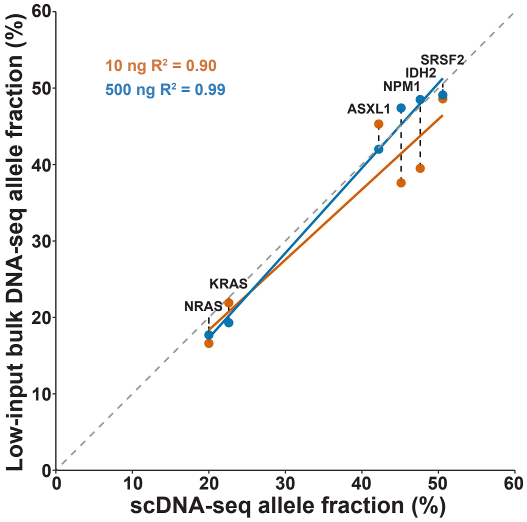


**Supplementary Figure 1. Concordance of ultra-low input bulk DNA-sequencing with single cell DNA-sequencing.** An AML sample with six pathologic somatic variants was sequenced using low-input bulk DNA sequencing with input of 10 ng or 500 ng of DNA, as well as by single-cell DNA-sequencing. The allele fraction (%) of each of the six variants is shown, as estimated using both methods. Correlation between allele fractions estimated using both methods is shown (orange line for 10 ng input, blue line for 500 ng input). Pearson correlation is shown.

### Supplementary Figure 2

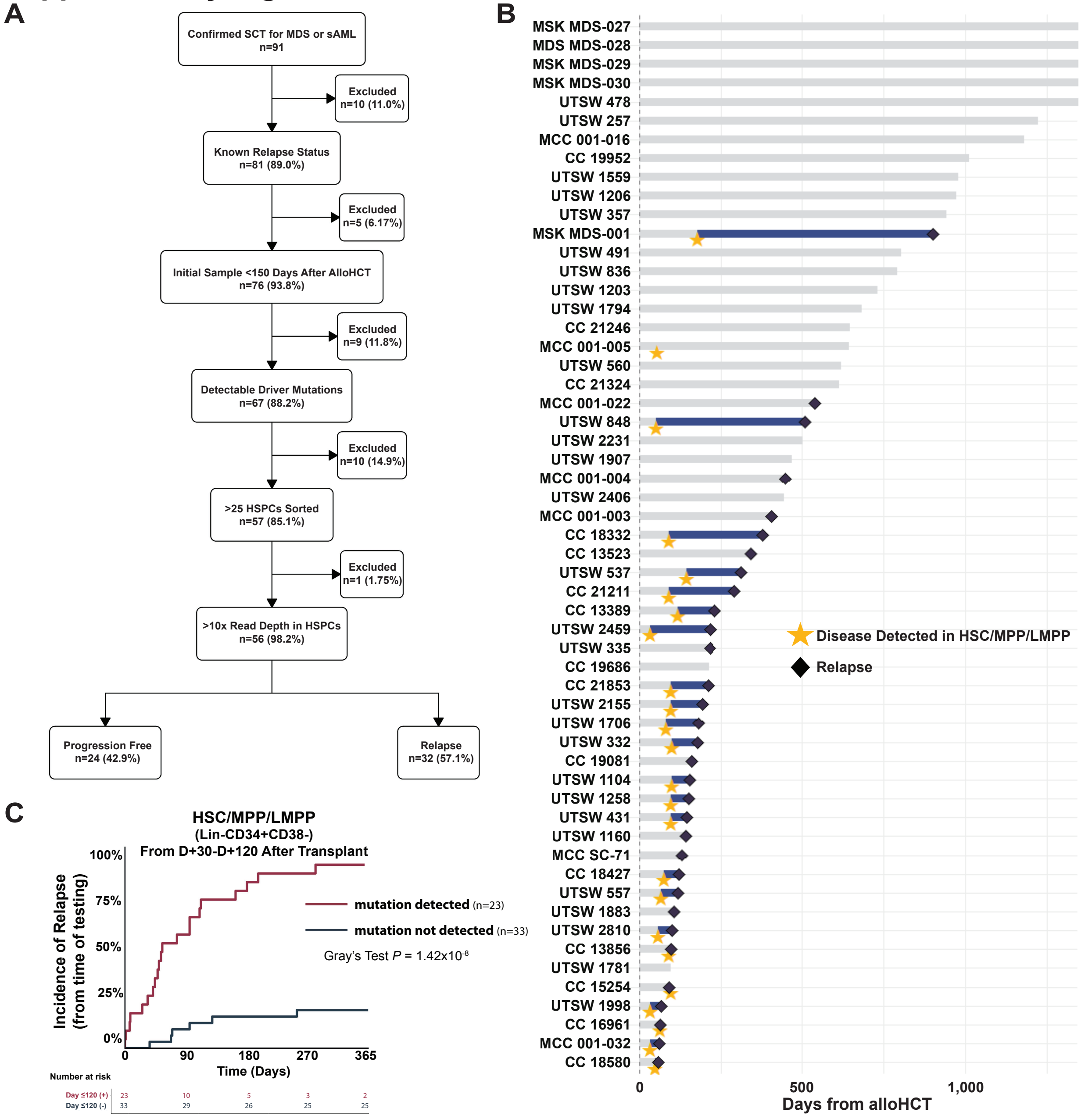


**Supplementary Figure 2. Multi-institutional patient cohort.** (**A**) CONSORT diagram of patient inclusion and exclusion criteria. (**B**) Swimmer's plot depiction of patient follow-up, indicating timepoint of relapse and death. (**C**) Cumulative incidence of relapse within the first year after testing, based on detection of any persistent pathogenic variant in hematopoietic stem cells (HSCs), multipotent progenitors (MPPs), or lymphoid primed multipotent progenitor cells (LMPPs); Statistical significance was assessed using Gray's test.

### Supplementary Figure 3

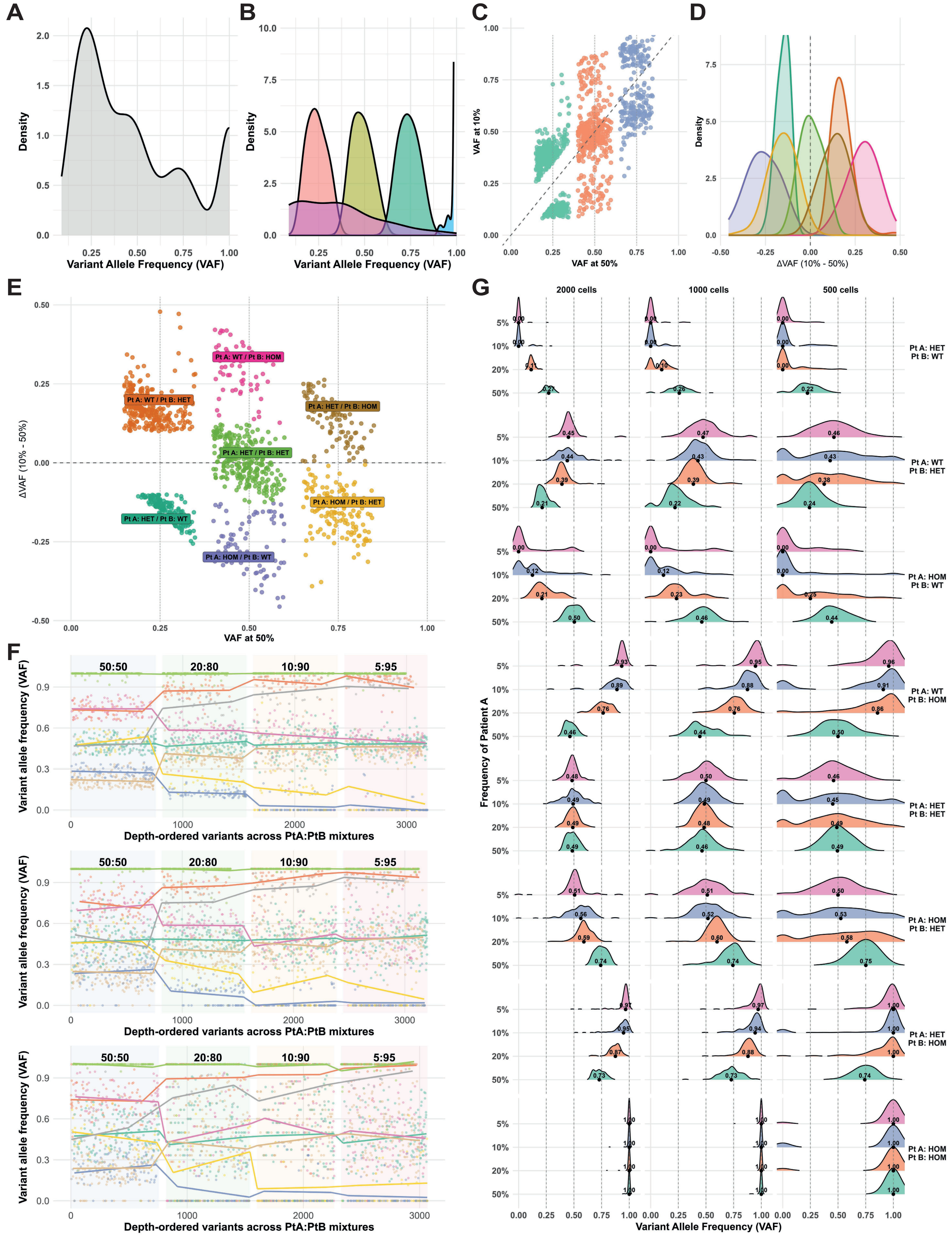

**Supplementary Figure 3. Identification of single nucleotide variants that distinguish patients.** (**A-B**) Density plots showing the distribution of variant allele frequencies (VAFs) for shared variants identified in the 50:50 PtA:PtB mixture (**A**), and the same variants grouped within ±0.1 of 0.25, 0.5, 0.75, and 1.0 (**B**), corresponding to expected allelic configurations in a balanced two-patient mixture. (**C**) Scatter plot of VAFs of binned germline variants in the 50:50 versus 10:90 PtA:PtB mixtures. The dashed line indicates the identity line (y = x). (**D**) Density plot of VAF shifts between the 50:50 and 10:90 mixtures (ΔVAF), colored by direction of change and initial 50:50 VAF group. (**E**) Scatter plot of VAF in the 50:50 mixture versus ΔVAF (10:90 – 50:50), colored by inferred PtA/PtB zygosity combination for each variant. (**F**) Scatter plots of VAF versus read depth for germline variants, stratified by PtA:PtB mixture (50:50, 20:80, 10:90, 5:95) and cellular input (2000, 1000, and 500 cells). Lines represent linear regression fits summarizing zygosity-specific VAF behavior across conditions. (**G**) Ridgeline plots showing VAF distributions for each inferred zygosity combination across PtA:PtB mixtures and cellular input levels.

### Supplementary Figure 4

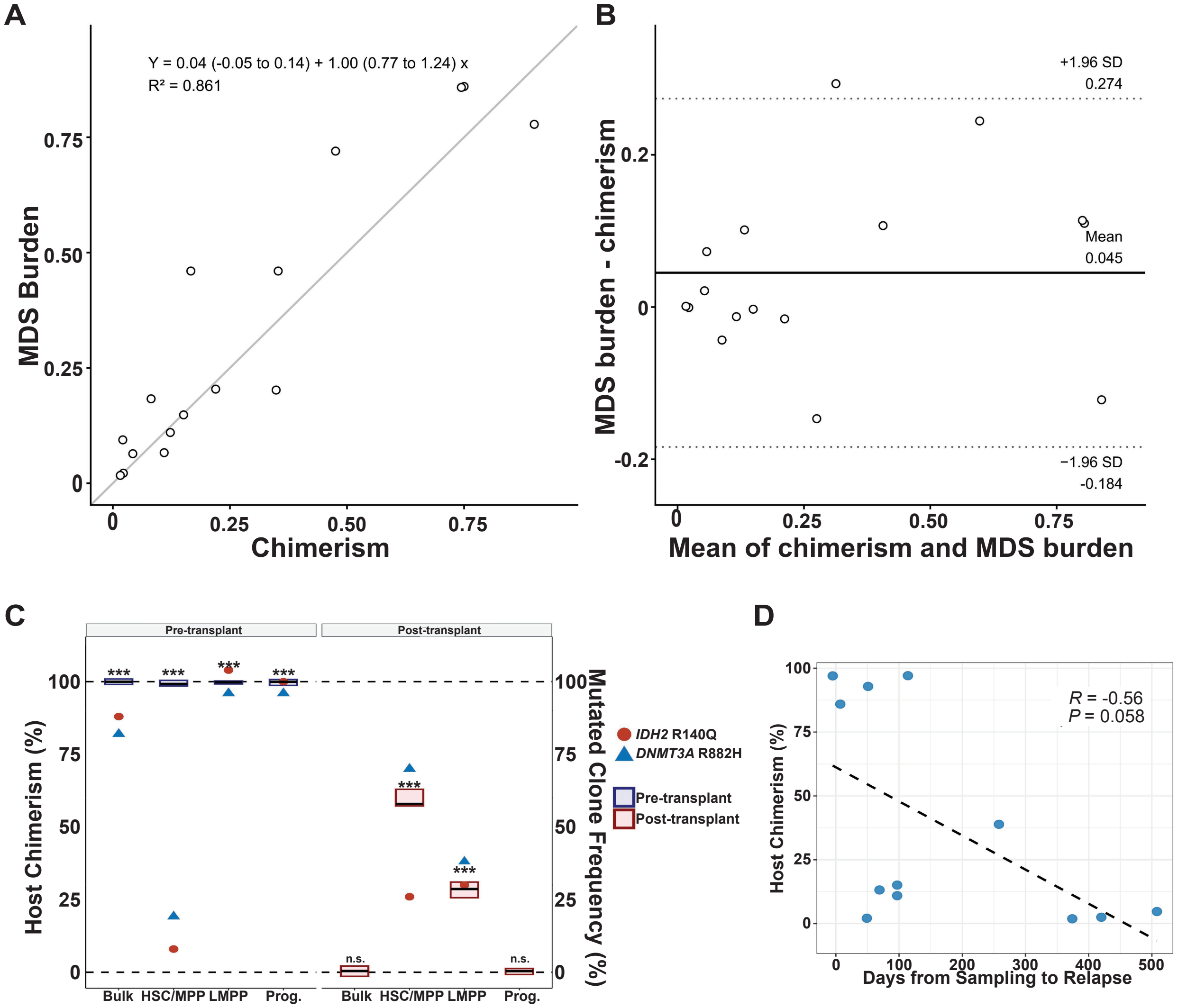


**Supplementary Figure 4. Concordance between MDS disease burden and host chimerism estimates.** (**A**) Correlation between MDS disease burden as estimated by driver mutation detection and estimated host chimerism. (**B**) Bland-Altman plot shows the difference between MDS burden and host chimerism plotted against the mean of their abundance. Solid and dashed lines indicate the mean bias and 95% confidence interval respectively. (**C**) Low-input sequencing-based chimerism estimates for **MSK MDS-001** across unfractionated bone marrow (bulk), hematopoietic stem cells and multipotent progenitors (HSC/MPP), lymphoid-primed multipotent progenitors (LMPP), and restricted progenitors (Prog.). Boxes indicate 95% confidence intervals of host chimerism, with shading denoting pre- (blue) and post-transplant (red) time points. Black lines represent chimerism estimates derived from variants with host:donor homozygous:wild-type genotypes. Symbols denote the estimated clonal fraction of cells harboring each driver mutation. (**D**) Correlation between host chimerism in HSC/MPP/LMPPs and time from testing to relapse.

### Supplementary Figure 5

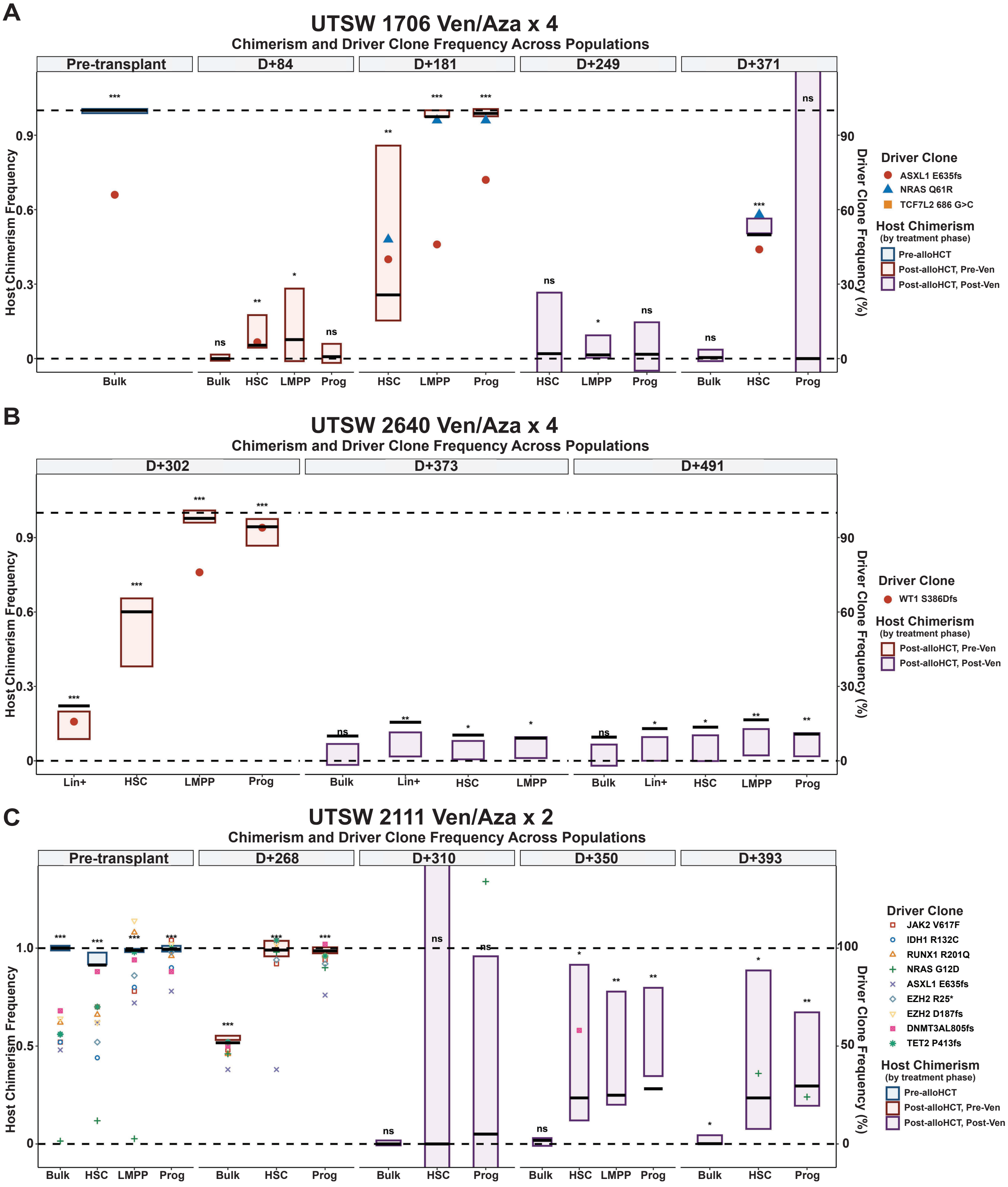

D

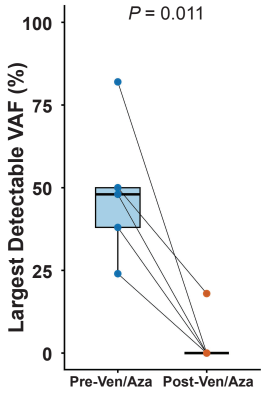


**Supplementary Figure 5. Venetoclax and azacitidine can deplete residual MDS HSCs after transplant.** (**A-D**) Low-input DNA-sequencing results for **UTSW 1706** (**A**), **UTSW 2640** (**B**), and **UTSW 2111** (**C**). Cell populations evaluated include unfractionated bone marrow (Bulk), hematopoietic stem cells (HSC), lymphoid primed multipotent progenitors (LMPP), and restricted progenitors (Prog). Boxes depict the 95% confidence interval of host chimerism, shaded to indicate pre-alloHCT (blue), post-alloHCT / pre-Ven/Aza (red), and post-alloHCT / post-Ven/Aza (purple). Black lines represent chimerism estimation based on host:homozygous donor:WT germline variants. The estimated clonal frequency of cells harboring driver mutations is shown by their corresponding symbols (see legend on right). Statistical significance was assessed using a one-tailed one-sample *t*-test comparing the mean host chimerism to zero; $*P<0.05$, $**P<0.01$, $***P<0.001$, not significant (n.s.). (**D**) In five patients who received post-transplant Ven/Aza with available driver mutation data, the largest detectable mutation variant allele frequency (VAF) prior to the first cycle and after the last cycle of Ven/Aza. Statistical significance was assessed using a a two-sided paired Student's *t*-test.

### Supplementary Figure 6

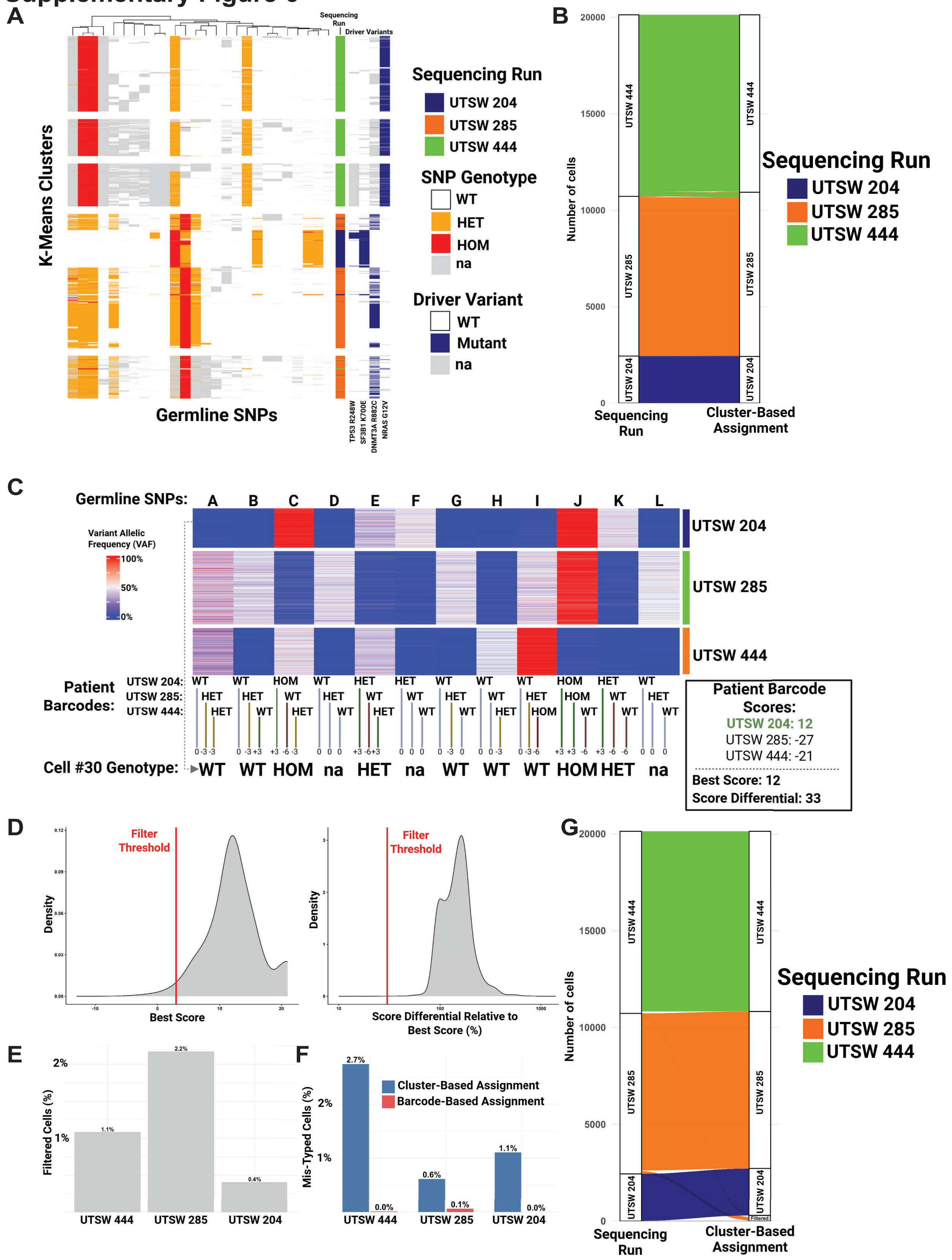

**H**

**Per patient score density with patient-specific threshold**

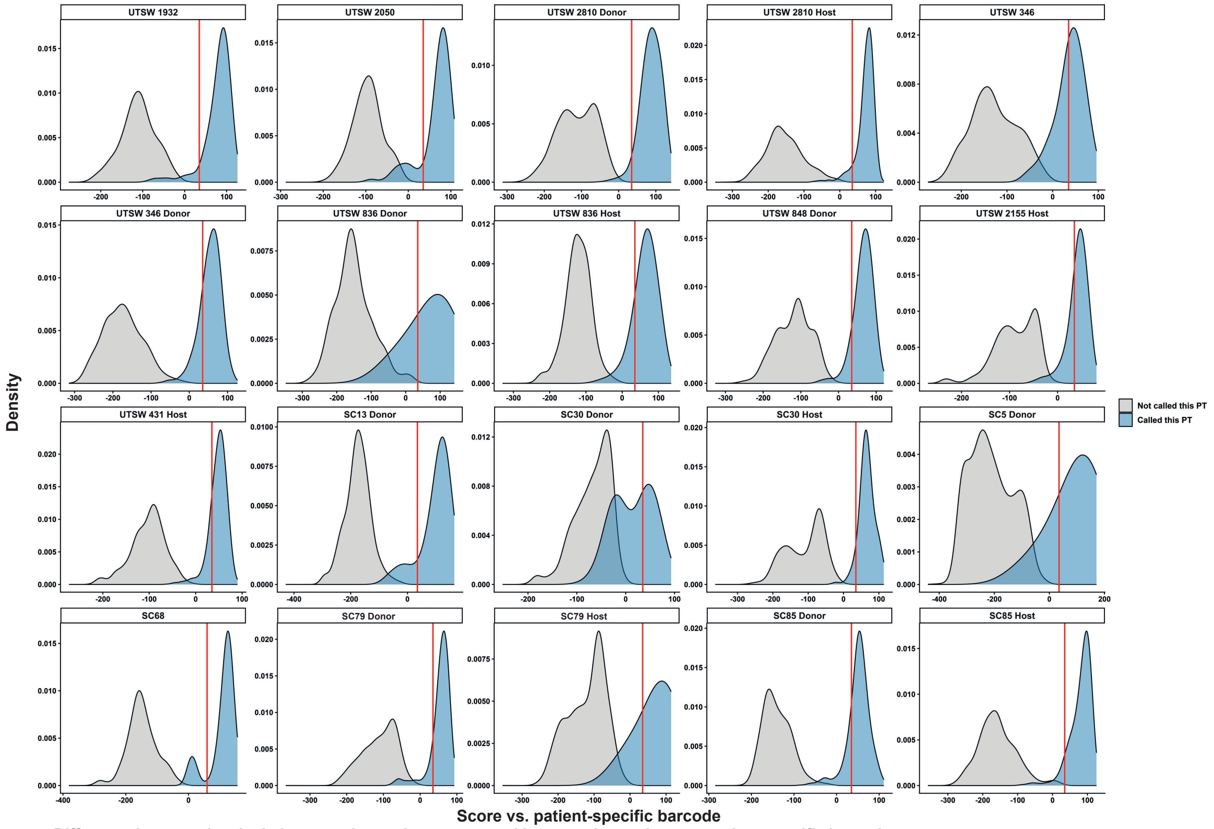


**I**

**Difference in score density in best scoring patient vs. second best scoring patient per patient-specific barcode**

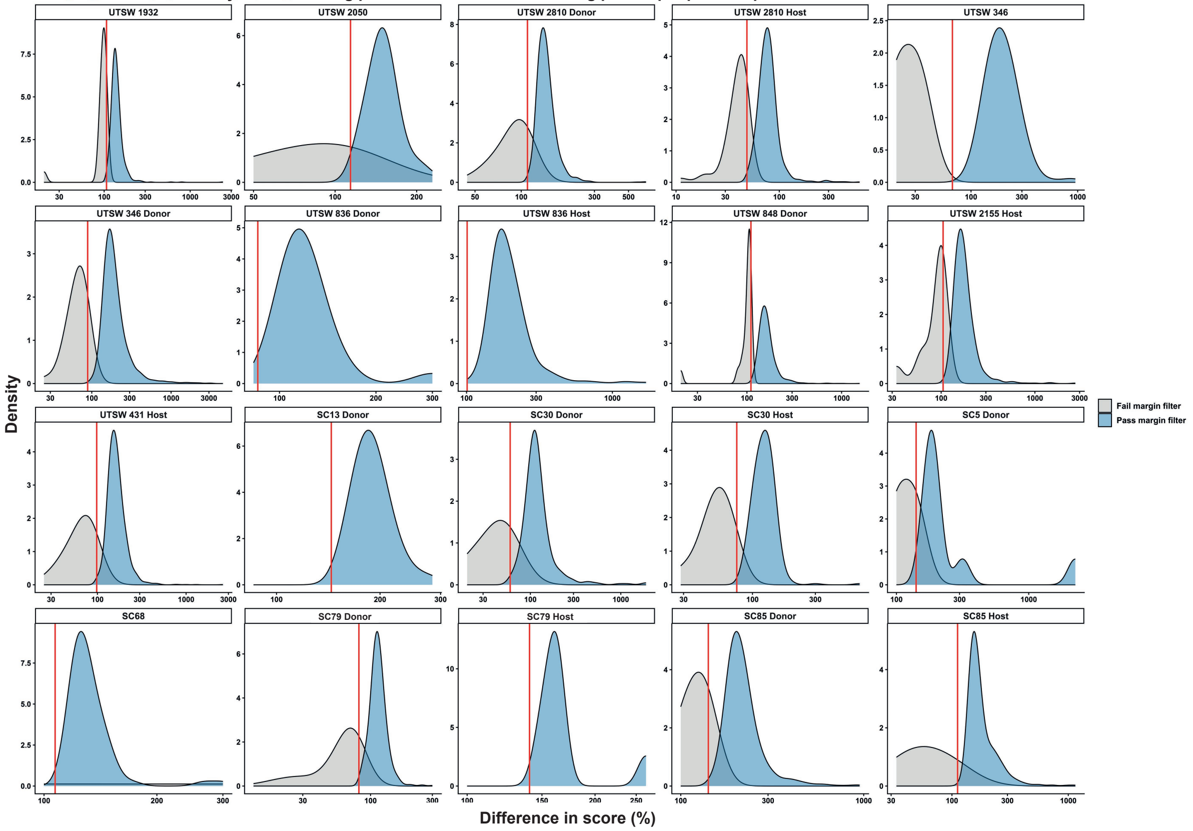

J

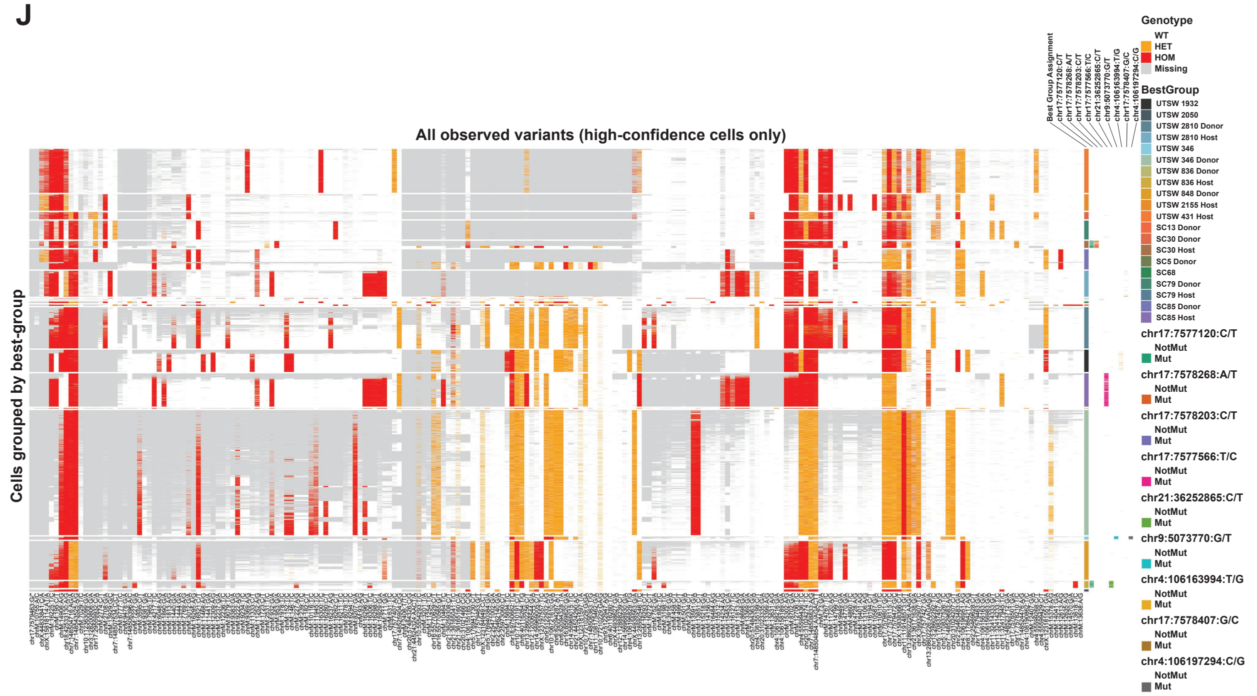


**Supplementary Figure 6. Demultiplexing Strategy for pooled cells in single-cell DNA-seq data.** (**A**) Heatmap of germline variant genotypes across individual cells captured in distinct sequencing runs from three individuals with AML. Cells are grouped by k-means clustering (k = 5). Annotation to the right show ground-truth sequencing run and mutation status for known pathogenic variants. Sample identification was initially performed agnostic to ground truth, and utilized pathologic variants to determine the origins of each genotype cluster. (**B**) Alluvial plot comparing ground-truth sample identity, defined by the sequencing run from which each cell originated, with cluster-based cell assignment. (**C**) Schematic of the germline barcode approach to cell multiplexing. Patient-specific barcodes are derived from cluster-level consensus genotypes. Individual cells are scored against each barcode for assignment on a per variant basis. Cells that match poorly to any patient barcode or exhibit insufficient separation between the top two matching scores are filtered. (**D**) Distribution of top barcode scores (left) and the differential between the 1$^{st}$ and 2$^{nd}$ highest barcode scores relative to the top score (right). Red lines indicating filtering thresholds. (**E**) Bar plot showing the percentage of cells filtered for each patient. (**F**) Bar plot showing the percentage of cells misassigned to each patient using cluster-based assignment (blue) and barcode-based assignment (red). (**G**) Alluvial plot comparing ground-truth sample identity, defined by the sequencing run from which each cell originated, with barcode-based cell assignment after filtering. (**H**) Density distributions of top barcode scores for cells assigned to each patient (blue) compared with all cells not assigned to that patient (gray). Red lines indicate patient-specific filtering thresholds. (**I**) Density distributions of score differentials between the best- and second-best-matching patient barcodes relative to the top match score for cells that passed filtering (blue) and cells that failed filtering (gray). Red lines indicate patient-specific filtering thresholds. (**J**) Heatmap of genotypes across the panel of nuclear and mitochondrial germline variants for all cells that passed filtering. Rows represent individual cells and columns represent variants; cells are grouped by final patient assignment. Genotypes are shown as wild-type (WT, white), heterozygous (HET, yellow), homozygous (HOM, red), or missing (gray). Annotations indicate final patient assignment and mutation status for pathogenic variants.

### Supplementary Figure 7

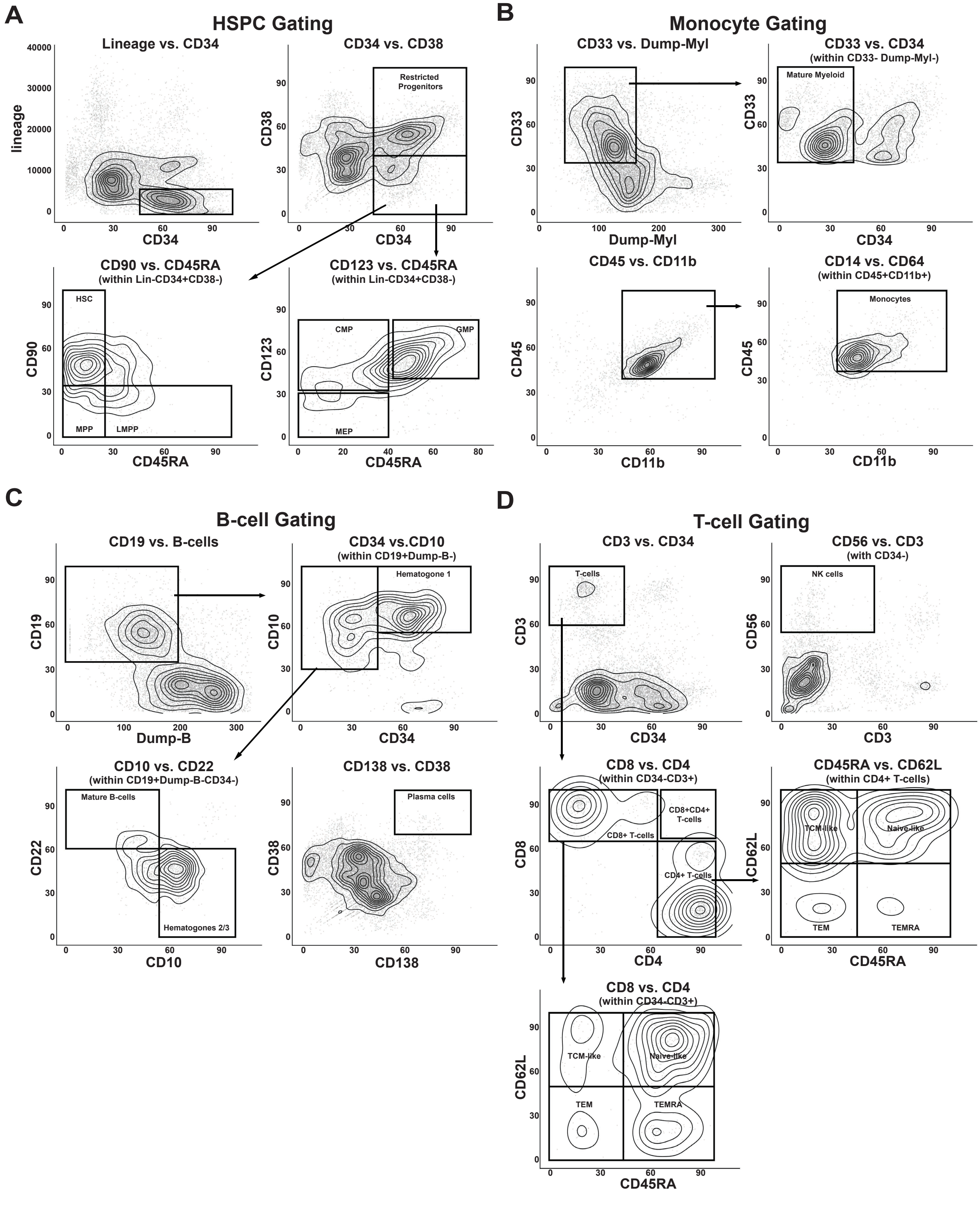

E

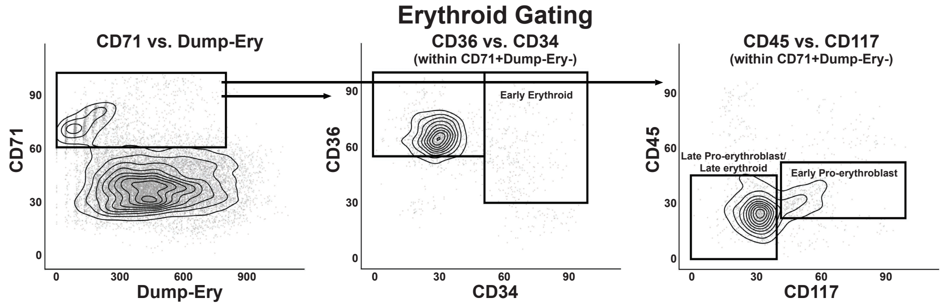
Erythroid Gating
CD71 vs. Dump-Ery
CD71
Dump-Ery
CD36 vs. CD34
(within CD71+Dump-Ery-)
Early Erythroid
CD36
CD34
CD45 vs. CD117
(within CD71+Dump-Ery-)
Late Pro-erythroblast/
Late erythroid
Early Pro-erythroblast
CD45
CD117

F

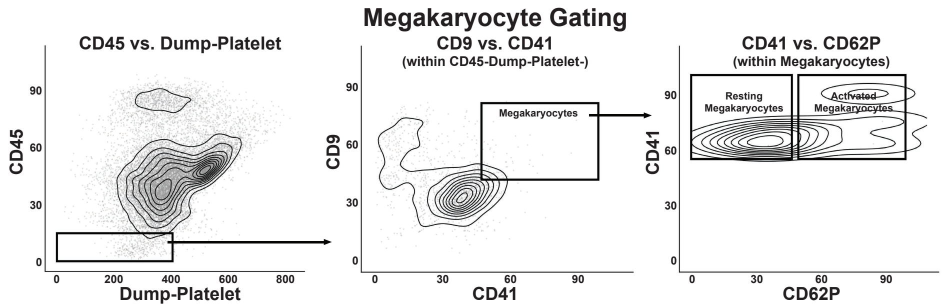
Megakaryocyte Gating
CD45 vs. Dump-Platelet
CD45
Dump-Platelet
CD9 vs. CD41
(within CD45-Dump-Platelet-)
Megakaryocytes
CD9
CD41
CD41 vs. CD62P
(within Megakaryocytes)
Resting
Megakaryocytes
Activated
Megakaryocytes
CD41
CD62P

G

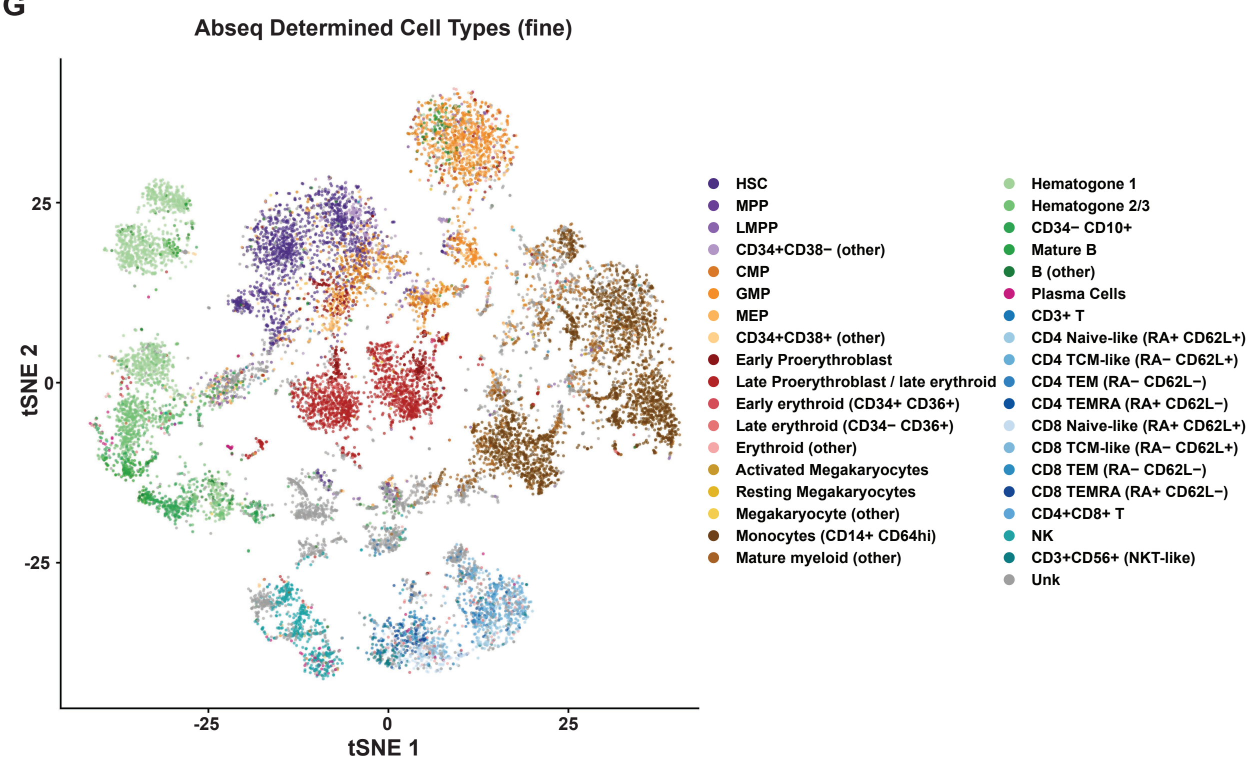
Abseq Determined Cell Types (fine)
tSNE 2
tSNE 1
HSC
MPP
LMPP
CD34+CD38− (other)
CMP
GMP
MEP
CD34+CD38+ (other)
Early Proerythroblast
Late Proerythroblast / late erythroid
Early erythroid (CD34+ CD36+)
Late erythroid (CD34− CD36+)
Erythroid (other)
Activated Megakaryocytes
Resting Megakaryocytes
Megakaryocyte (other)
Monocytes (CD14+ CD64hi)
Mature myeloid (other)
Hematogone 1
Hematogone 2/3
CD34− CD10+
Mature B
B (other)
Plasma Cells
CD3+ T
CD4 Naive-like (RA+ CD62L+)
CD4 TCM-like (RA− CD62L+)
CD4 TEM (RA− CD62L−)
CD4 TEMRA (RA+ CD62L−)
CD8 Naive-like (RA+ CD62L+)
CD8 TCM-like (RA− CD62L+)
CD8 TEM (RA− CD62L−)
CD8 TEMRA (RA+ CD62L−)
CD4+CD8+ T
NK
CD3+CD56+ (NKT-like)
Unk

**Supplementary Figure 7. Cell surface protein measurements by Abseq allow for discrimination of cell types.** Cell types (color-coded) were assigned by flow cytometry-like gating (**A-F**). Fine-grained populations are shown here; broader groupings are presented in **Fig.5b**. Bi-axial flow cytometry-like cell surface protein expression plots from scDNA-sequencing data are shown for identification of hematopoietic stem and progenitors cells (HSPCs)(**A**), monocytes (**B**), B cells and plasma cells (**C**), T cell subsets (**D**), and cells from erythroid (**E**) and megakaryocyte (**F**) lineages. **Lineage** - CD3, CD4, CD5, CD7, CD8, CD10, CD11b, CD14, CD19, CD22, CD56, CD71; **Dump-Myl** - CD3, CD56, CD4, CD8, CD71, CD19; Dump-B - CD3, CD56, CD14, CD33, CD11b, CD4, CD8, CD71; **Dump-Ery** - CD3, CD4, CD8, CD2, CD5, CD7, CD19, CD22, CD10, CD56, CD244, CD11b, CD14, CD16, CD33, CD64, CD163, CD11c, CD13, CD18, CD32, HLA.DR, CD74, CD123, CD1c, CD303, CD304; **Dump-Platelet** - CD45, CD71, CD14, CD64, CD33, CD11b, CD16, CD163, CD11c, CD13, HLA.DR, CD3, CD19, CD56. (**G**) Cell surface protein-based clustering of 12,032 cells from 14 patients (12 MDS, one wild-type control, one clonal hematopoiesis control).

### Supplementary Figure 8

A

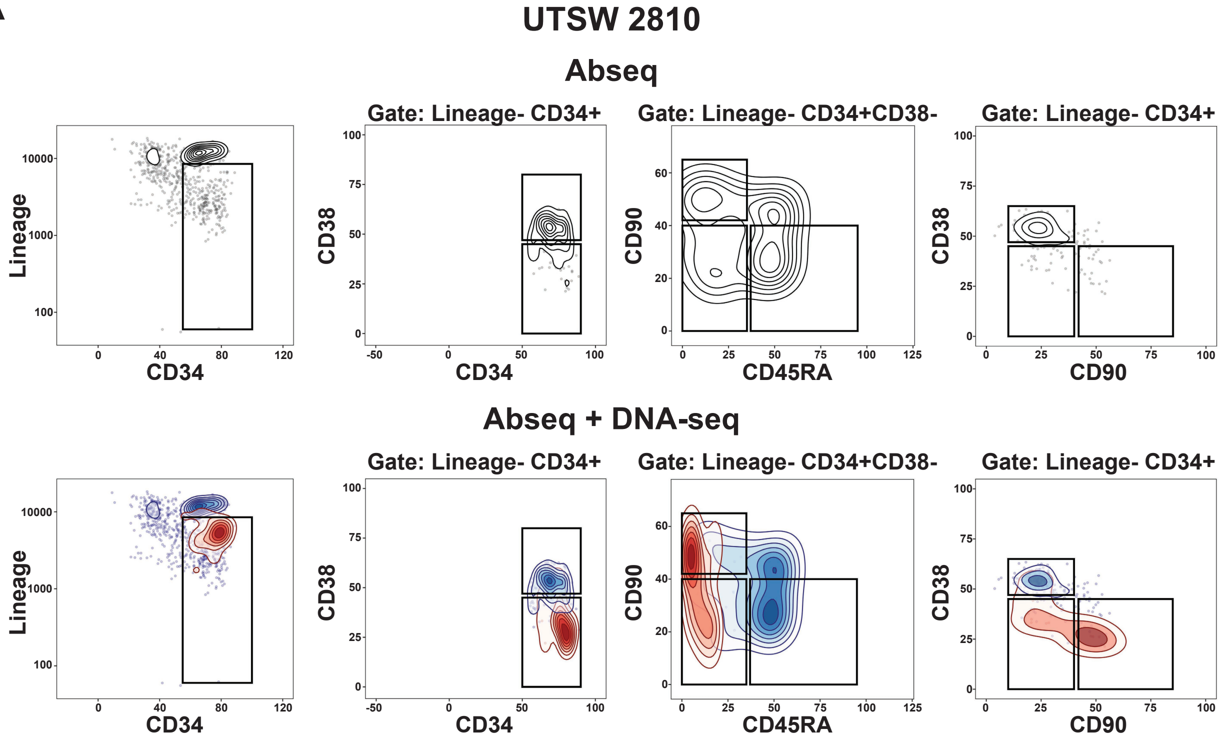


B

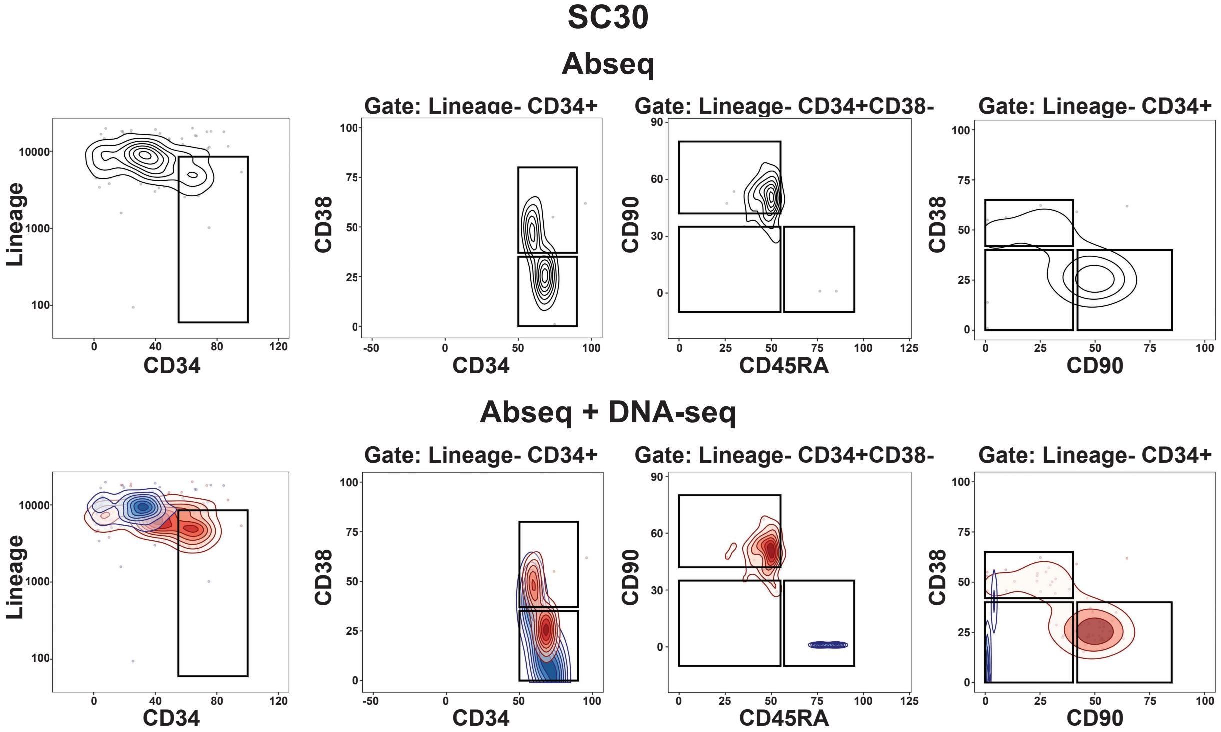

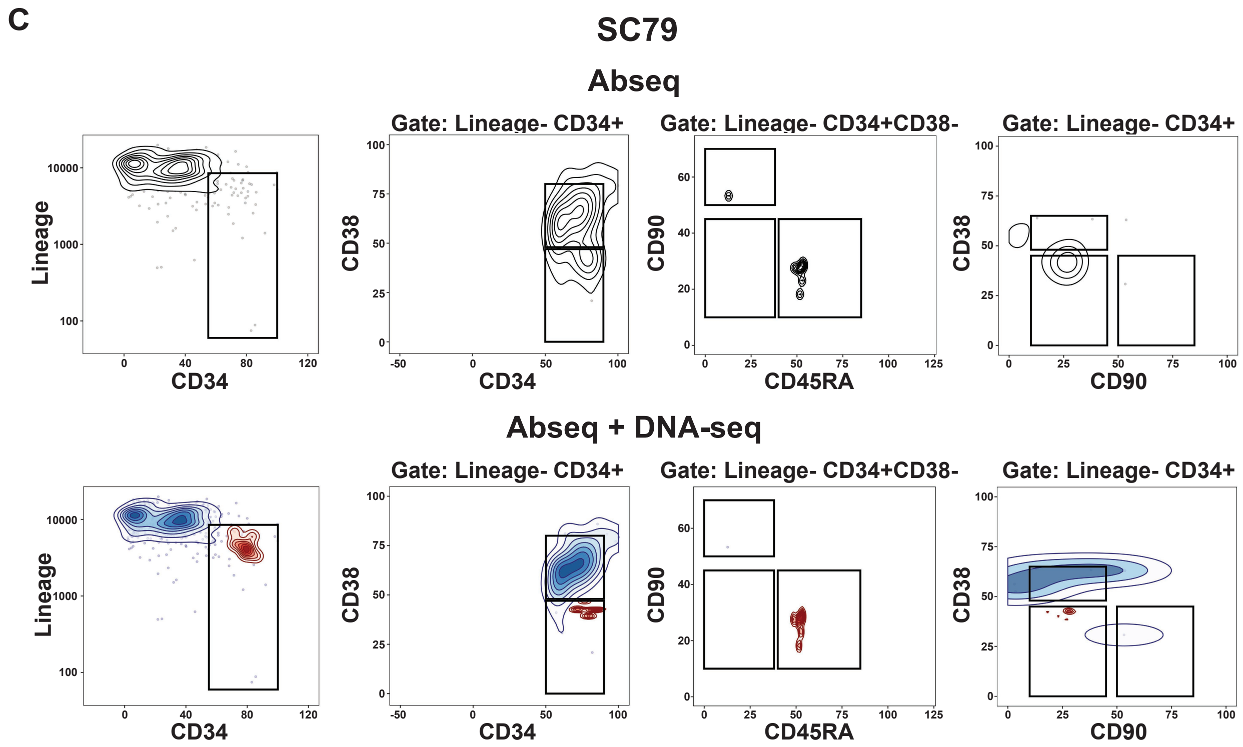


**Supplementary Figure 8. Single-cell DNA-sequencing and Abseq show enrichment of MDS cells in immature cell populations.** (**A-C**) Bi-axial flow cytometry-like cell surface protein expression (Abseq) plots from scDNA-sequencing data are shown for identification of hematopoietic stem cells (HSC), multipotent progenitors (MPP), lymphoid primed multipotent progenitors (LMPP), and restricted progenitors (Prog) from **UTSW 2810** (**A**), **SC30** (**B**), and **SC79** (**C**). Top rows show Abseq plots and bottom rows show Abseq plots with separate densities for host MDS (red) and donor (blue) cells.

#### Supplementary Figure 9

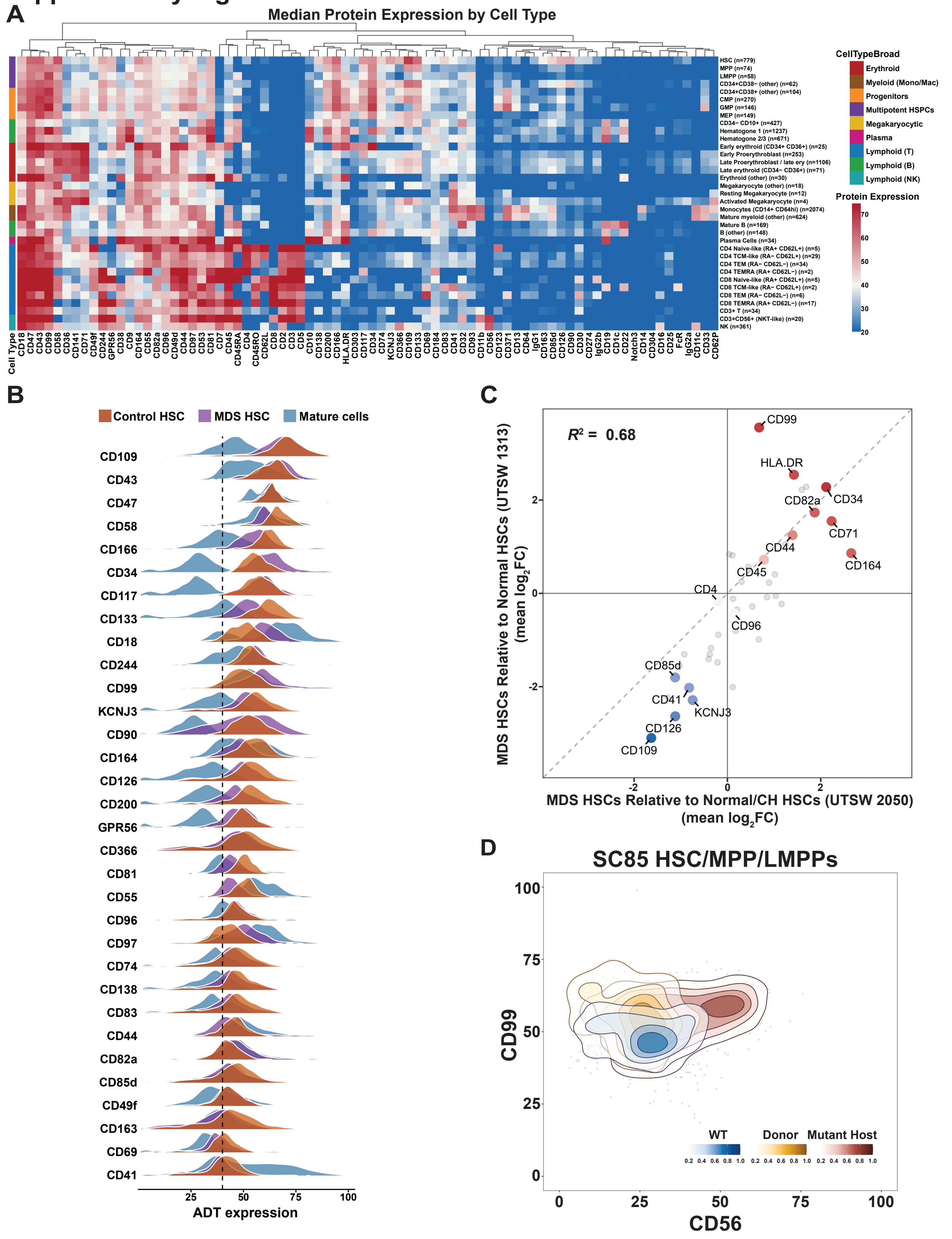

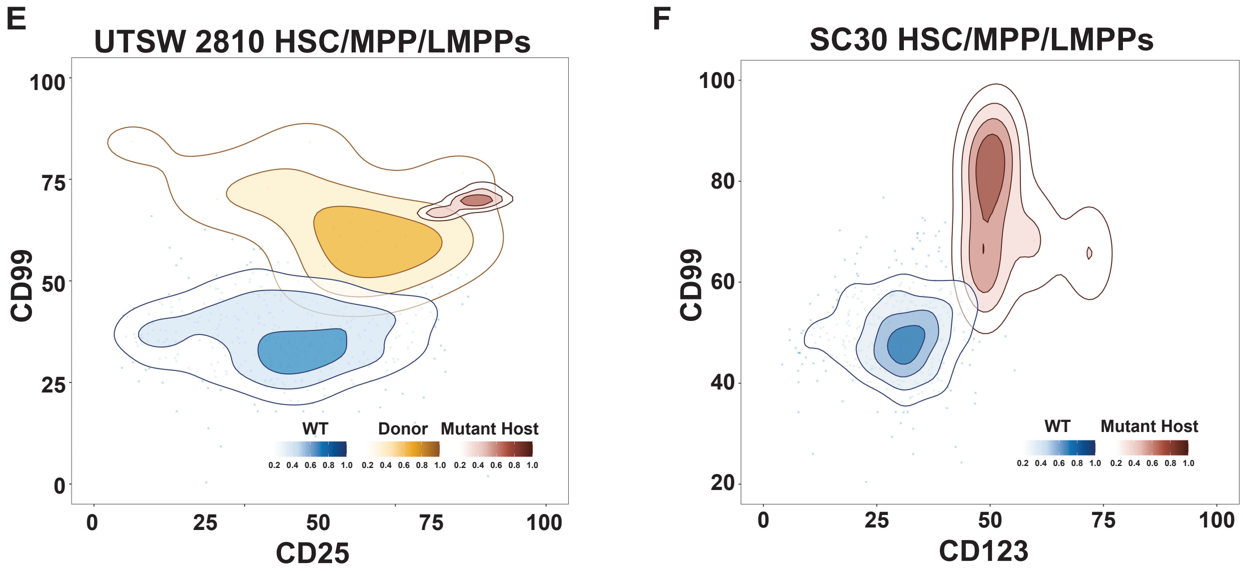


**Supplementary Figure 9. Aberrantly expressed cell surface proteins distinguish MDS from donor and normal control stem cells.** (**A**) Cell surface expression (Abseq) of 78 candidate cell surface proteins (columns) in scDNA-seq of wild-type cells (both donor and normal controls). Rows indicate cell types as identified by flow cytometry-like subsetting (shown in **Supp.Fig.7**), with the number of cells evaluated indicated. (**B**) 32 cell surface proteins robustly expressed on normal control HSCs were selected, and their relative expression on normal, donor, and MDS HSCs is shown. (**C**) Correlation between log-fold change in expression of candidate MDS HSC cell surface proteins relative to HSCs from two normal controls (one with CH [x-axis] and one without CH [y-axis]). (**D-F**) Bi-axial flow cytometry-like plot of candidate MDS HSC cell surface proteins on HSC/MPP/LMPPs from MDS patients (red) as compared with matched donor cells present in the same specimen (yellow) or normal controls (blue). Markers best distinguishing MDS HSC/MPP/LMPPs are shown for SC85 (**D**), **UTSW 2810** (**E**), and **SC30** (**F**).

#### Supplementary Figure 10

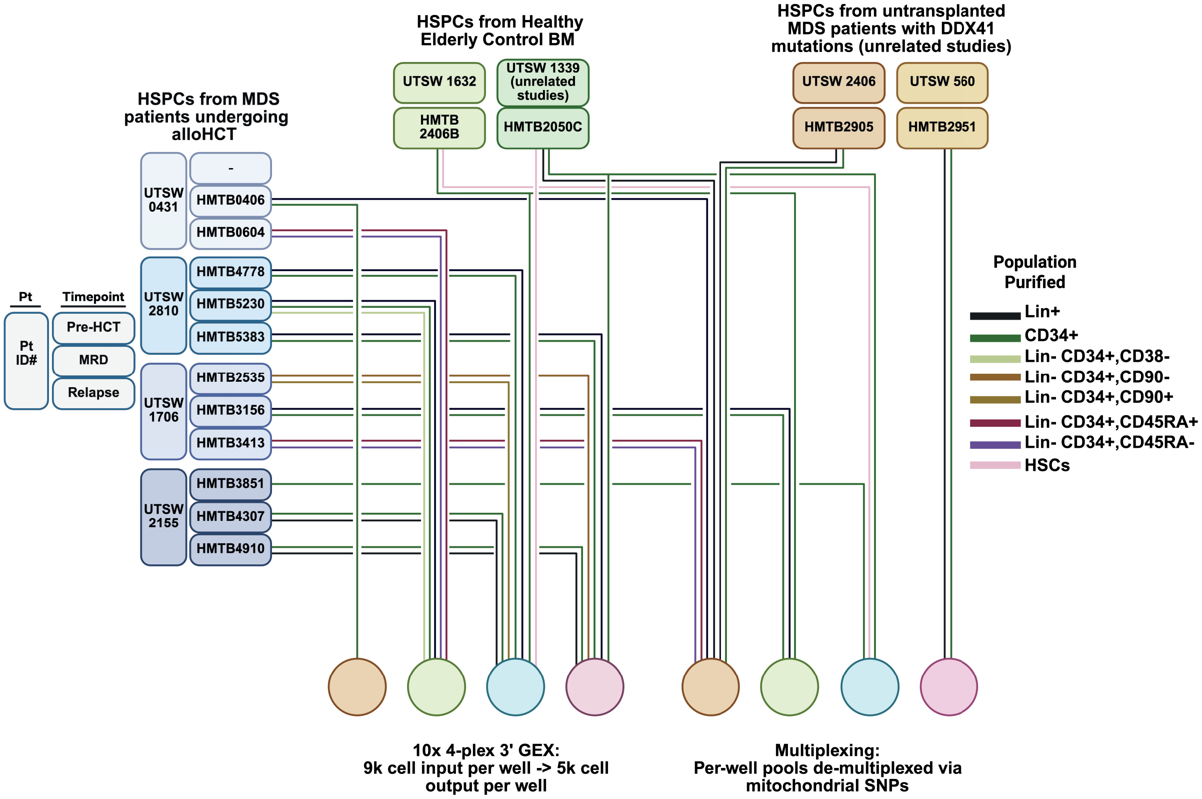


**Supplementary Figure 10. Subway map diagram illustrating multiplexing strategy for single-cell transcriptional profiling of HSPCs from multiple patients.** Primary bone marrow (BM) samples are shown in the left column and top row, grouped by patient. Each track represents a distinct immunophenotypic population isolated from a given sample, colored by population (lineage+ [Lin+], black; Lin-CD34+, dark green; Lin-CD34+CD38-, light green; Lin-CD34+CD90-, dark brown; Lin-CD34+CD90+, light brown; Lin-CD34+CD45RA+, red; Lin-CD34+CD45RA-, purple; HSCs [Lin-CD34+CD38-CD90+CD45RA-], pink). Tracks converge into multiplexed pools corresponding to 10x Genomics 4-plex 3′ wells. Each well contains a unique set of cell barcodes, enabling assignment of cells to their capture well, while individual patient identities within each pool were resolved post hoc using mitochondrial variant-based de-multiplexing.

### Supplementary Figure 11

A

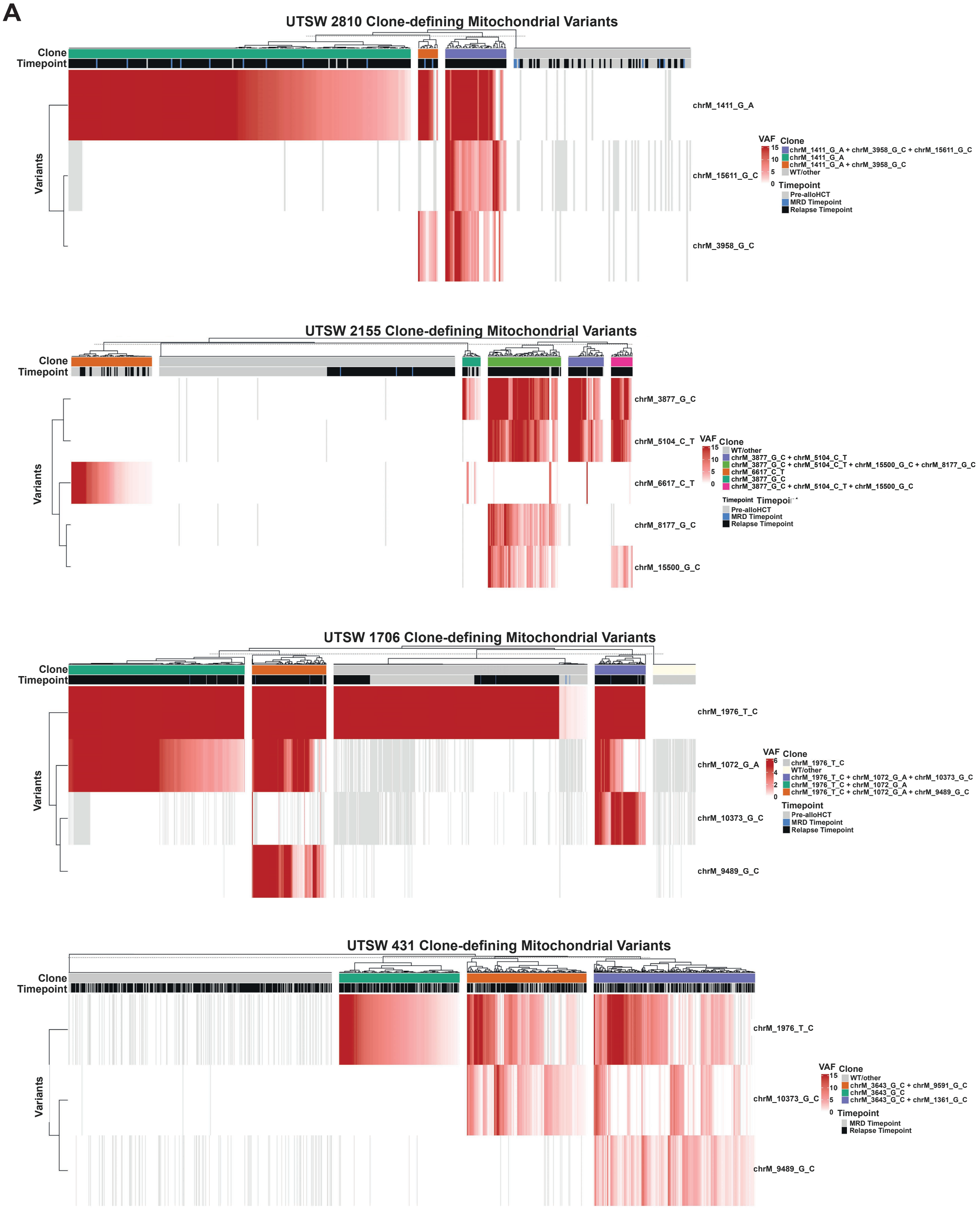

**B**

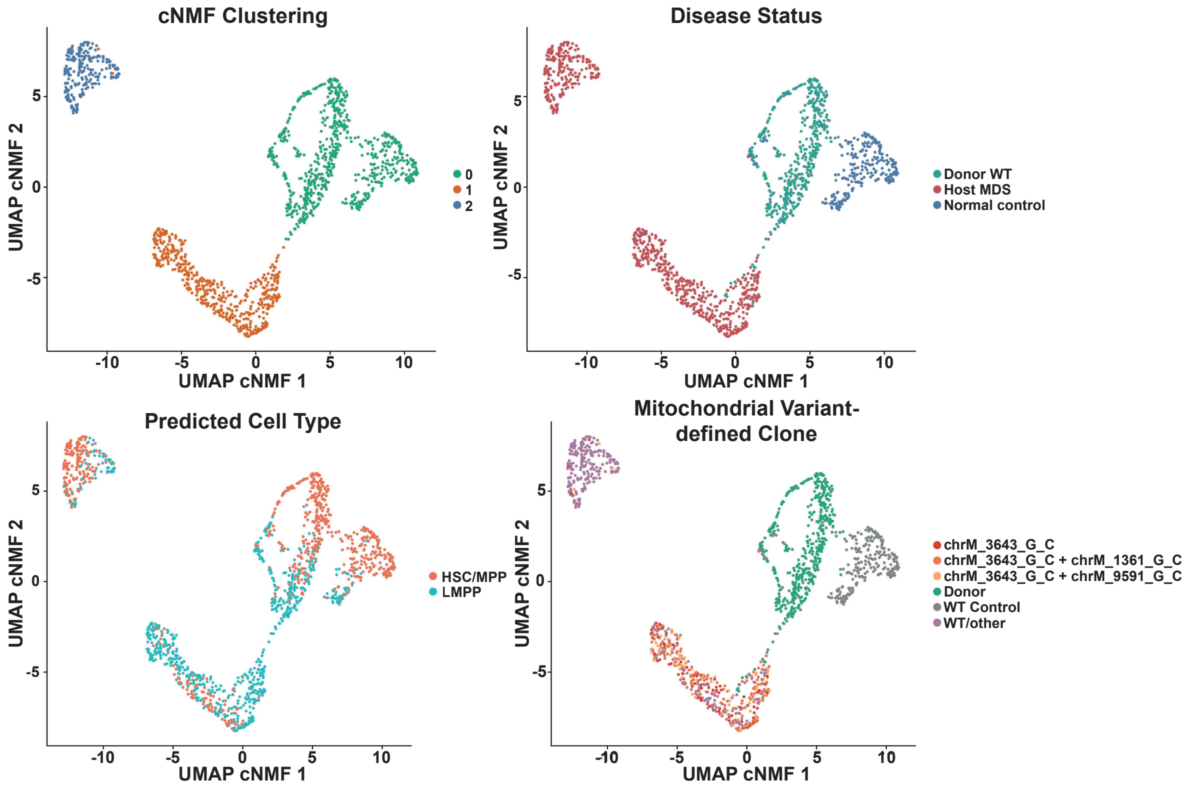
cNMF Clustering
Disease Status
Predicted Cell Type
Mitochondrial Variant-defined Clone
UMAP cNMF 1
UMAP cNMF 2
0
1
2
Donor WT
Host MDS
Normal control
HSC/MPP
LMPP
chrM_3643_G_C
chrM_3643_G_C + chrM_1361_G_C
chrM_3643_G_C + chrM_9591_G_C
Donor
WT Control
WT/other

**C**

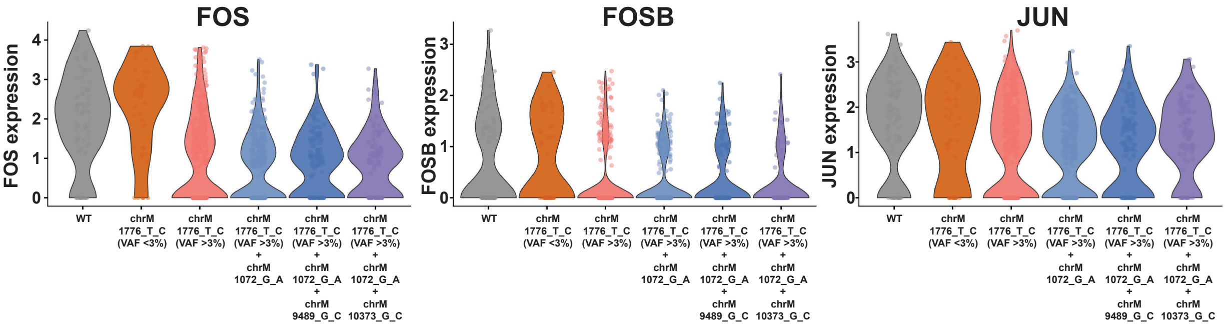
FOS
FOSB
JUN
FOS expression
FOSB expression
JUN expression
WT
chrM 1776_T_C (VAF <3%)
chrM 1776_T_C (VAF >3%)
chrM 1776_T_C (VAF >3%) + chrM 1072_G_A
chrM 1776_T_C (VAF >3%) + chrM 1072_G_A + chrM 9489_G_C
chrM 1776_T_C (VAF >3%) + chrM 1072_G_A + chrM 10373_G_C

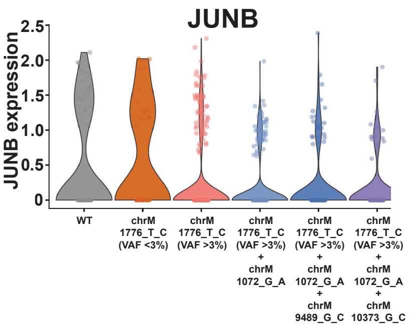
JUNB
JUNB expression
WT
chrM 1776_T_C (VAF <3%)
chrM 1776_T_C (VAF >3%)
chrM 1776_T_C (VAF >3%) + chrM 1072_G_A
chrM 1776_T_C (VAF >3%) + chrM 1072_G_A + chrM 9489_G_C
chrM 1776_T_C (VAF >3%) + chrM 1072_G_A + chrM 10373_G_C

**D**

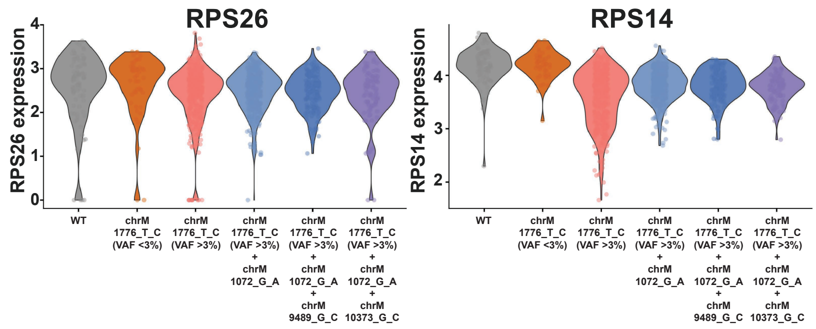
RPS26
RPS14
RPS26 expression
RPS14 expression
WT
chrM 1776_T_C (VAF <3%)
chrM 1776_T_C (VAF >3%)
chrM 1776_T_C (VAF >3%) + chrM 1072_G_A
chrM 1776_T_C (VAF >3%) + chrM 1072_G_A + chrM 9489_G_C
chrM 1776_T_C (VAF >3%) + chrM 1072_G_A + chrM 10373_G_C

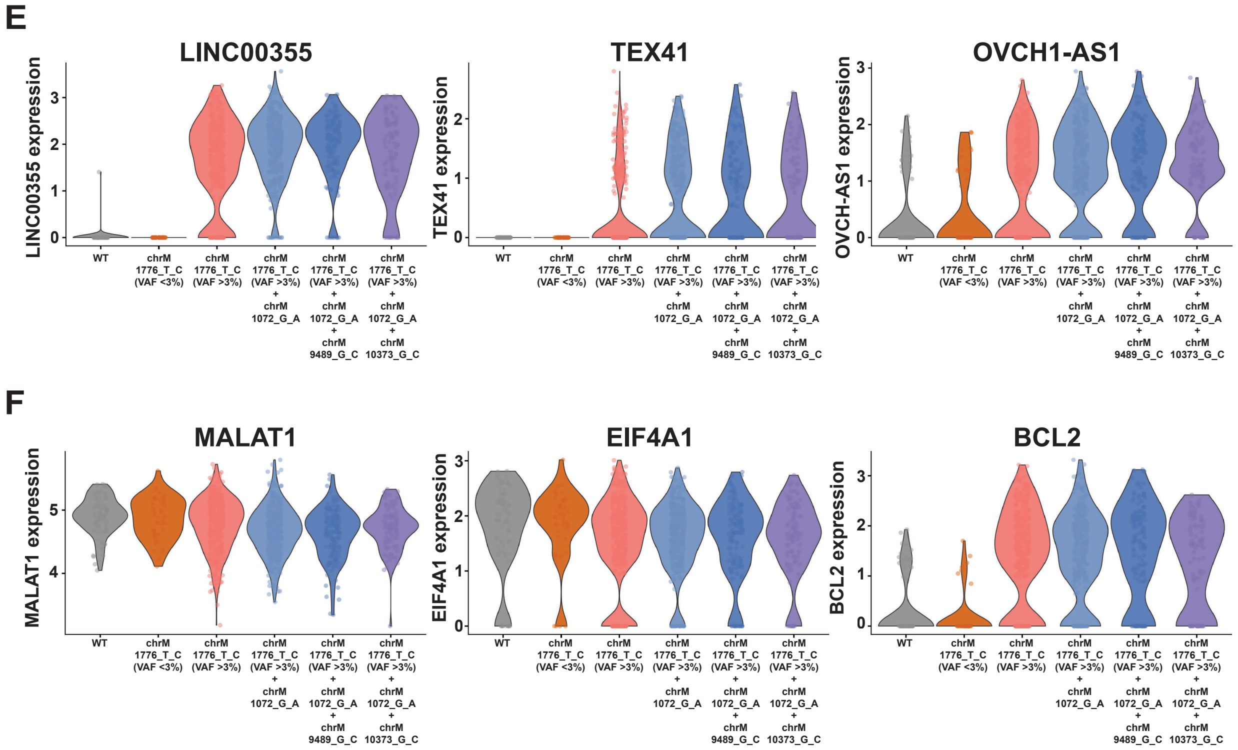


**Supplementary Figure 11. Somatic mitochondrial variants distinguish disease subclones.** (**A**) Heatmaps showing per-cell variant allelic frequencies (VAFs) for patient-specific mitochondrial variants exhibiting significant changes between pre-HCT and relapse, defined by >10% absolute difference in frequency, ≥1.25-fold change, and FDR < 0.05 (Fisher's exact test). Rows represent variants and columns represent individual cells, hierarchically clustered by VAF profile. VAFs were filtered by minimum coverage (>20 reads for UTSW 2810 and >50 reads for all others), with missing values shown in gray. Columns are annotated by inferred clone and timepoint. (**B**) UMAP embedding of HSC/MPP/LMPPs from UTSW 431 and normal control HSCs based on consensus non-negative matrix factorization (cNMF) modules. Cells are colored by cNMF-based clustering (top left), origin/disease status (top right), predicted cell type (bottom left), or mitochondrial variant-defined subclone (bottom right). MDS clusters segregate by clone identity, with expanded subclones clones largely confined to the cluster with predominant LMPP-like transcriptional features. (**C-F**) Violin plots showing normalized expression of selected gene classes across mitochondrial variant-defined subclones in HSC/MPP/LMPPs from UTSW 1706. Genes include AP-1 transcription factors (FOS, FOSB, JUN, JUNB)(**C**), ribosomal proteins (RPS26, RPS14)(**D**), MDS-associated lncRNAs (LINC00355, TEX41, OVCH1-AS1)(**E**), and potential modulators of venetoclax sensitivity (MALAT1, EIF4A1, and BCL2)(**F**).

### Supplementary Figure 12

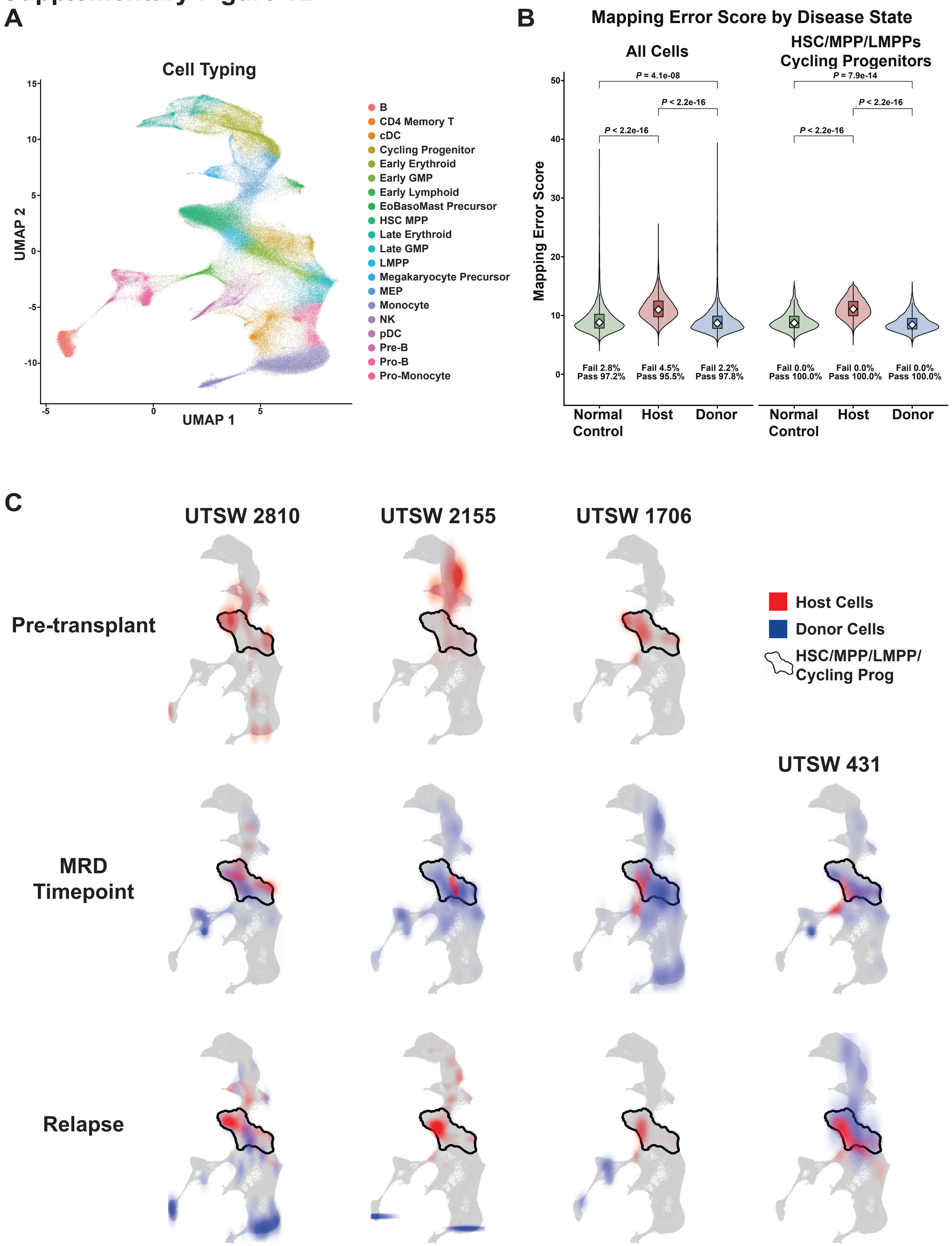

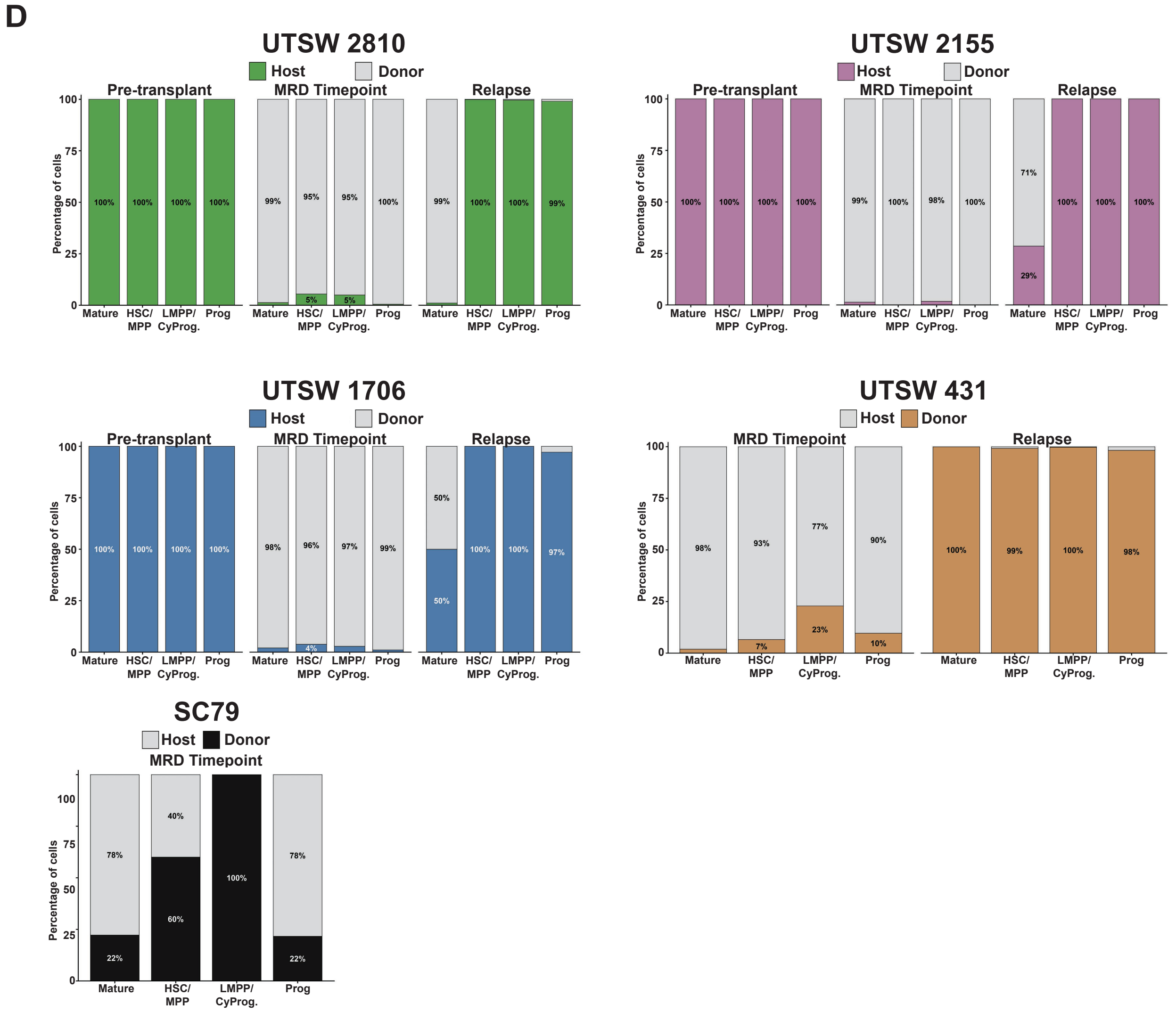


**Supplementary Figure 12. Distribution of donor and host cells along hematopoietic differentiation.** (**A**) Hematopoietic reference (BoneMarrowMap) with annotation for cell types. (**B**) Violin plots of per-cell mapping error scores for WT (green), MDS (red), and donor WT (blue) cells following projection onto the BoneMarrowMap reference, calculated as the weighted Mahalanobis distance to reference centroids (higher = lower confidence). Shown for all cells (left) and primitive compartments (HSC, MPP, LMPP, cycling progenitors; right). Cells were classified as failing the mapping error filter if their scores exceeded 2.5 median absolute deviations (MAD) from the median. Group differences were assessed using a Kruskal–Wallis test, with pairwise comparisons performed using two-sided Wilcoxon rank-sum tests and Benjamini–Hochberg correction for multiple testing. (**C**) Kernel density estimates of donor (blue) and host (red) cells projected onto the hematopoietic reference for **UTSW 2810**, **UTSW 2155**, **UTSW 1706**, and **UTSW 431** at pre-transplant, MRD timepoint, and relapse. (**D**) Stacked bar plots showing the proportion of donor (gray) and host (colored by patient) cells across hematopoietic compartments (mature cells, HSC/MPP, LMPP/cycling progenitors [CyProg.], progenitors [Prog]) for each bone marrow sample across patients and timepoints. Percentages are indicated within each bar.

#### Supplementary Figure 13

A

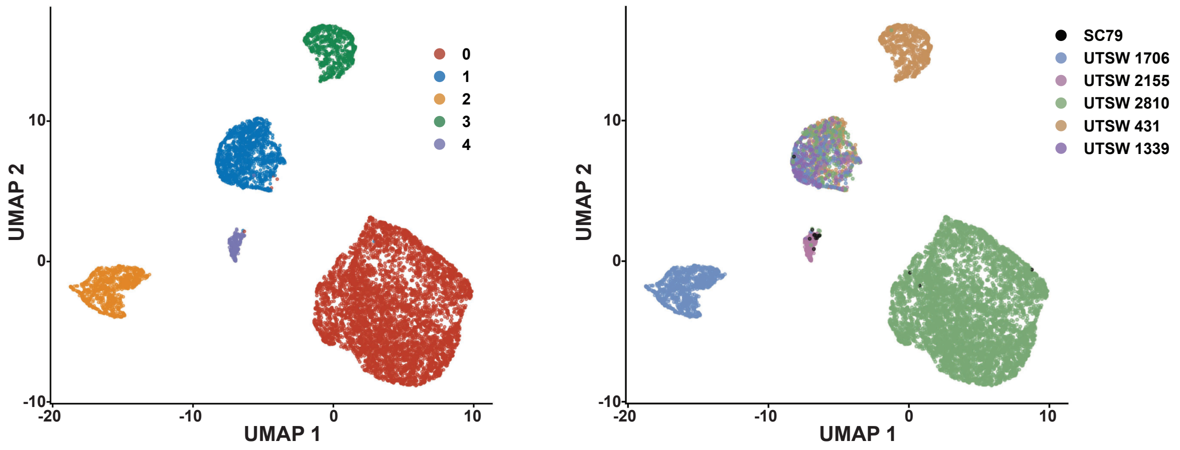


B

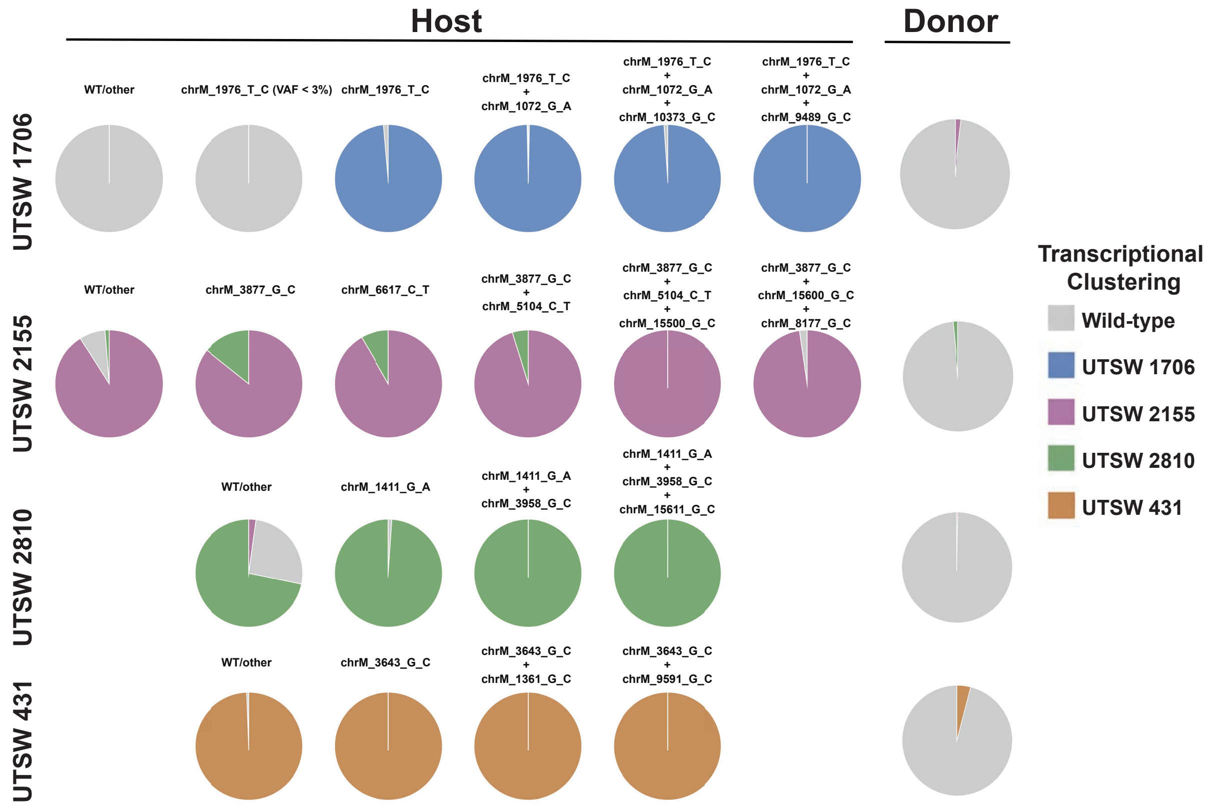

C

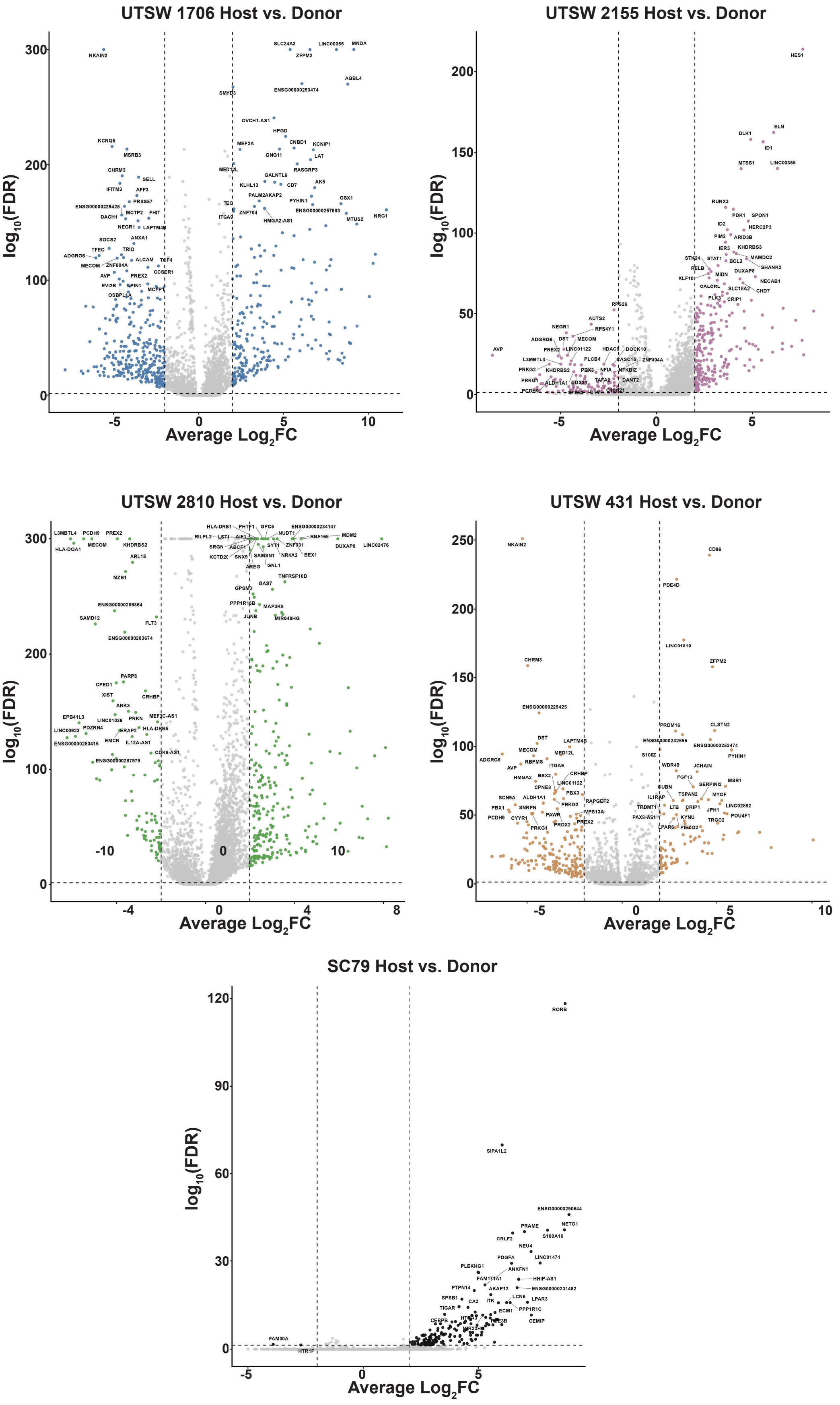

UTSW 1706 Host vs. Donor
UTSW 2155 Host vs. Donor
UTSW 2810 Host vs. Donor
UTSW 431 Host vs. Donor
SC79 Host vs. Donor
log10(FDR)
Average Log2FC

D

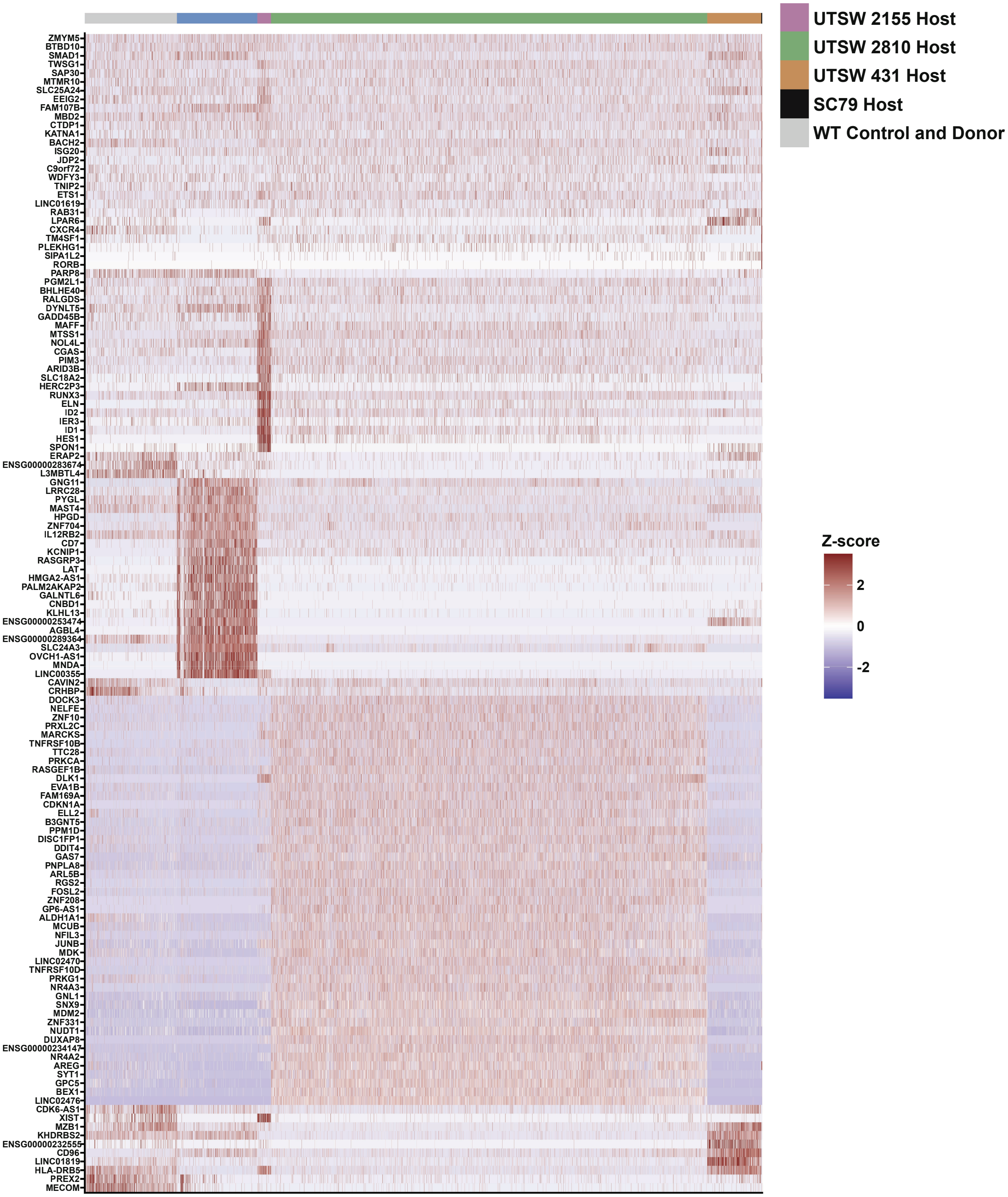

UTSW 2155 Host
UTSW 2810 Host
UTSW 431 Host
SC79 Host
WT Control and Donor
ZMYM5
BTBD10
SMAD1
TWSG1
SAP30
MTMR10
SLC25A24
EEIG2
FAM107B
MBD2
CTDP1
KATNA1
BACH2
ISG20
JDP2
C9orf72
WDFY3
TNIP2
ETS1
LINC01619
RAB31
LPAR6
CXCR4
TM4SF1
PLEKHG1
SIPA1L2
RORB
PARP8
PGM2L1
BHLHE40
RALGDS
DYNLT5
GADD45B
MAFF
MTSS1
NOL4L
CGAS
PIM3
ARID3B
SLC18A2
HERC2P3
RUNX3
ELN
ID2
IER3
ID1
HES1
SPON1
ERAP2
ENSG00000283674
L3MBTL4
GNG11
LRRC28
PYGL
MAST4
HPGD
ZNF704
IL12RB2
CD7
KCNIP1
RASGRP3
LAT
HMGA2-AS1
PALM2AKAP2
GALNTL6
CNBD1
KLHL13
ENSG00000253474
AGBL4
ENSG00000289364
SLC24A3
OVCH1-AS1
MNDA
LINC00355
CAVIN2
CRHBP
DOCK3
NELFE
ZNF10
PRXL2C
MARCKS
TNFRSF10B
TTC28
PRKCA
RASGEF1B
DLK1
EVA1B
FAM169A
CDKN1A
ELL2
B3GNT5
PPM1D
DISC1FP1
DDIT4
GAS7
PNPLA8
ARL5B
RGS2
FOSL2
ZNF208
GP6-AS1
ALDH1A1
MCUB
NFIL3
JUNB
MDK
LINC02470
TNFRSF10D
PRKG1
NR4A3
GNL1
SNX9
MDM2
ZNF331
NUDT1
DUXAP8
ENSG00000234147
NR4A2
AREG
SYT1
GPC5
BEX1
LINC02476
CDK6-AS1
XIST
MZB1
KHDRBS2
ENSG00000232555
CD96
LINC01819
HLA-DRB5
PREX2
MECOM
Z-score
2
0
-2

E

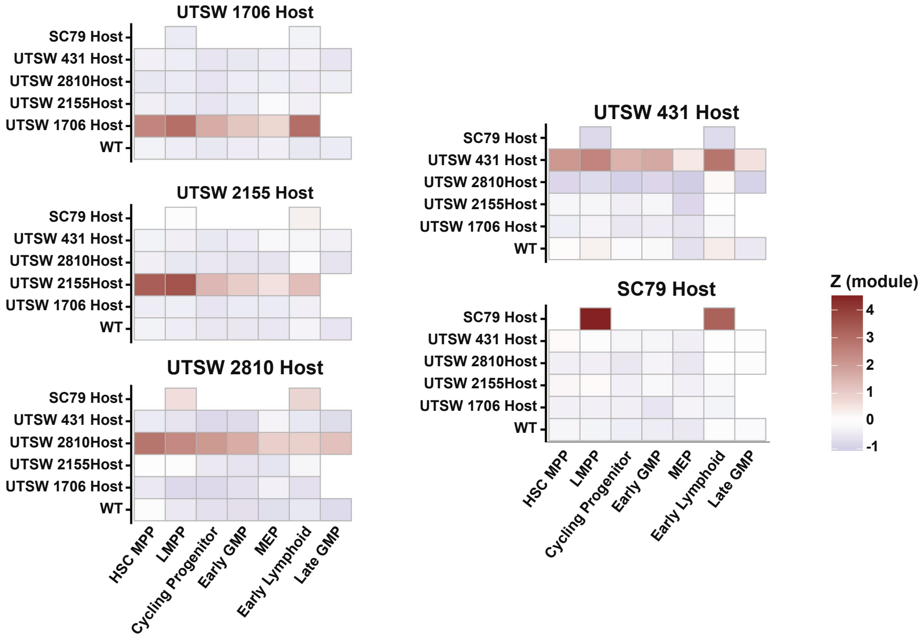
UTSW 1706 Host
UTSW 2155 Host
UTSW 2810 Host
UTSW 431 Host
SC79 Host
SC79 Host
UTSW 431 Host
UTSW 2810Host
UTSW 2155Host
UTSW 1706 Host
WT
HSC MPP
LMPP
Cycling Progenitor
Early GMP
MEP
Early Lymphoid
Late GMP
Z (module)
4
3
2
1
0
-1

F

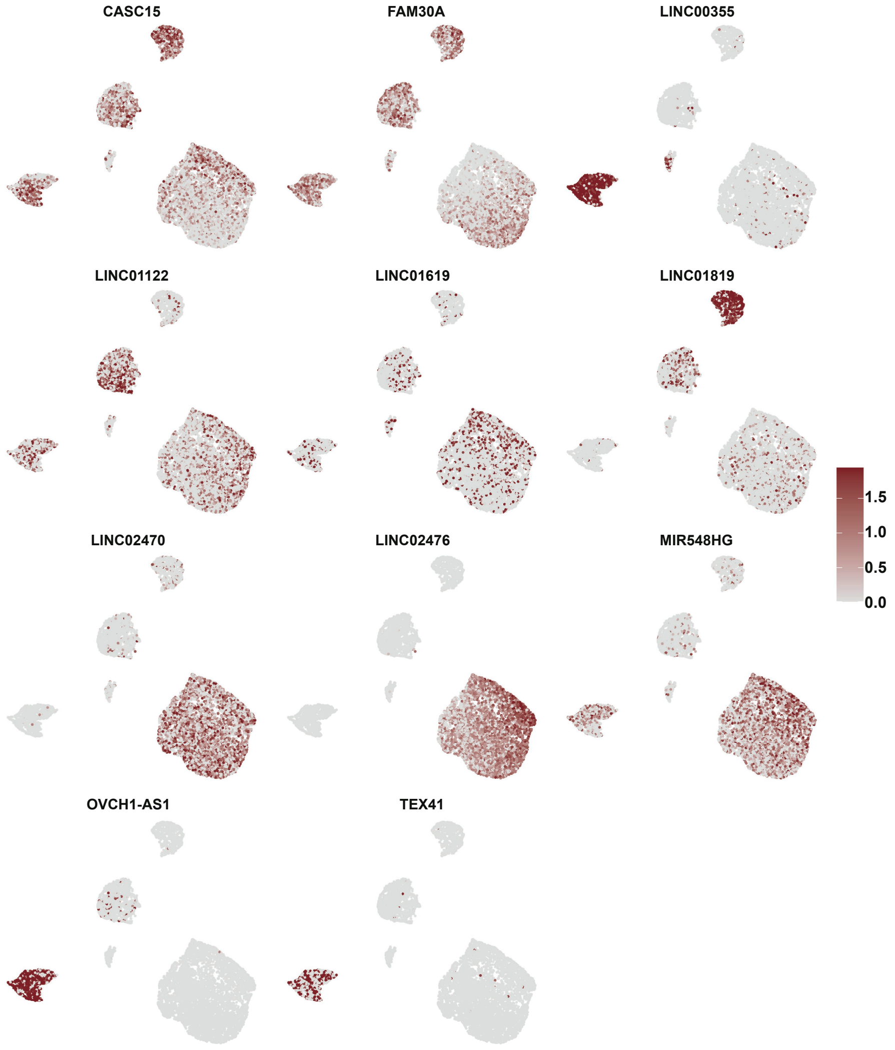
CASC15
FAM30A
LINC00355
LINC01122
LINC01619
LINC01819
LINC02470
LINC02476
MIR548HG
OVCH1-AS1
TEX41
1.5
1.0
0.5
0.0

**Supplementary Figure 13. Transcriptional programs distinguish MDS from wild-type control and donor HSCs.** (**A**) UMAP projection of HSC/MPP/LMPP cells computed from per-cell consensus non-negative matrix factorization (cNMF) program usage (k = 7). Cells are colored by cluster (left) and patient of origin (right). (**B**) Pie charts showing the distribution of patient-specific or wild-type donor or normal control transcriptional clusters within mitochondrially defined subclones for each patient. Each pie represents a distinct subclone, with slices indicating the percentage of HSC/MPP/LMPP cells assigned to each transcriptional cluster. (**C**) Volcano plots showing differential gene expression in patient-specific MDS HSC/MPP/LMPP clusters from each indicated patient relative to the corresponding WT cluster. Genes with FDR < 0.05 and |log2 fold-change| > 2 are highlighted in patient-specific colors. (**D**) Cell-level heatmap showing z-scored expression of HSC/MPP/LMPP cells for patient-specific gene modules. Genes were defined as cluster markers expressed in >50% of cells within the target population and <25% of cells in the comparison group. Columns represent individual cells, ordered by patient, with the top annotation bar indicating sample group; donor-derived and WT control and donor cells are shown in gray. (**E**) Scaled heatmap of mean module scores for patient-specific gene sets across hematopoietic populations (HSC/MPP, LMPP, cycling progenitor, MEP, early lymphoid, early GMP, late GMP). Rows represent distinct host-derived cells; WT includes donor-derived and wild-type control and donor cells. Scores are averaged per group and population and scaled within each module. Only group–population combinations with ≥5 cells were included. (**F**) UMAP of HSC/MPP/LMPP cells showing scaled expression of differentially expressed lncRNAs (defined by >40% absolute difference in detection frequency in MDS vs WT). Color intensity indicates z-scored expression, capped at the 95th percentile. XIST and TTTY14 were excluded due to confounding sex-dependent expression.

### Supplementary Figure 14

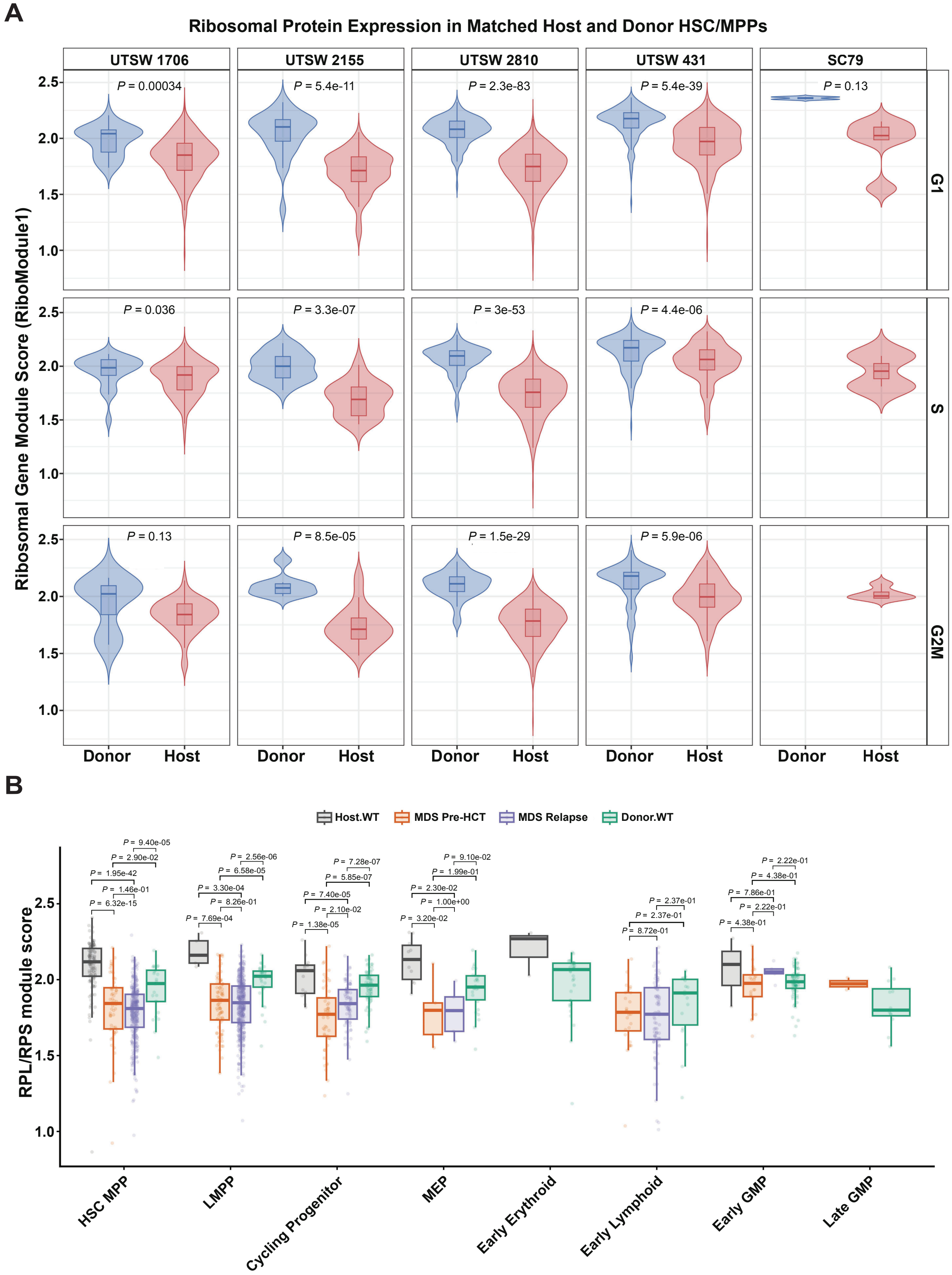

C

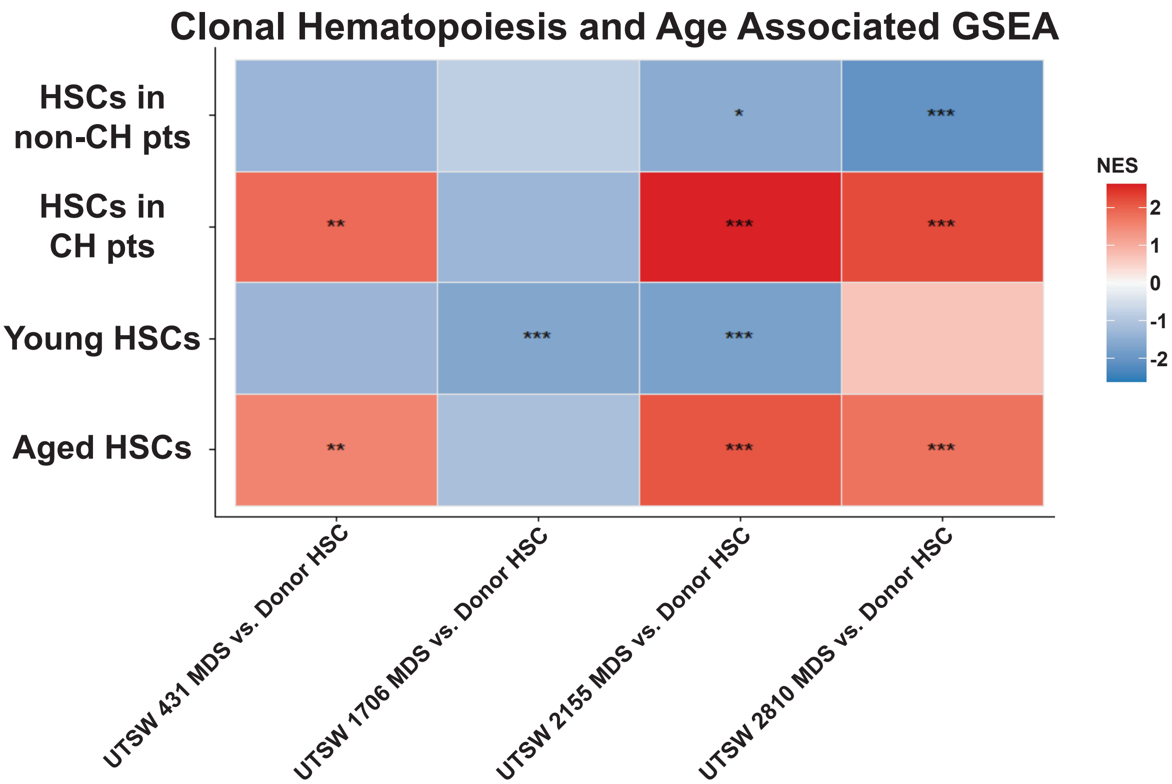
Clonal Hematopoiesis and Age Associated GSEA
HSCs in non-CH pts
HSCs in CH pts
Young HSCs
Aged HSCs
NES
2
1
0
-1
-2
*
**
***
UTSW 431 MDS vs. Donor HSC
UTSW 1706 MDS vs. Donor HSC
UTSW 2155 MDS vs. Donor HSC
UTSW 2810 MDS vs. Donor HSC

D

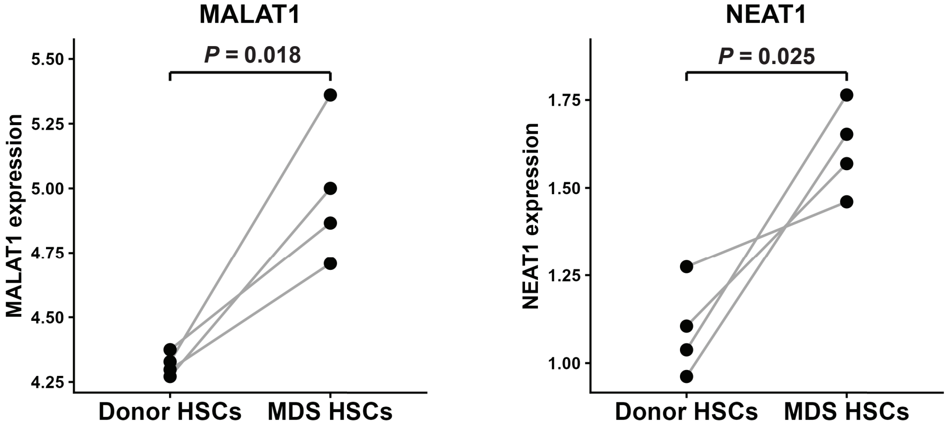
MALAT1
P = 0.018
MALAT1 expression
5.50
5.25
5.00
4.75
4.50
4.25
Donor HSCs
MDS HSCs
NEAT1
P = 0.025
NEAT1 expression
1.75
1.50
1.25
1.00
Donor HSCs
MDS HSCs

E

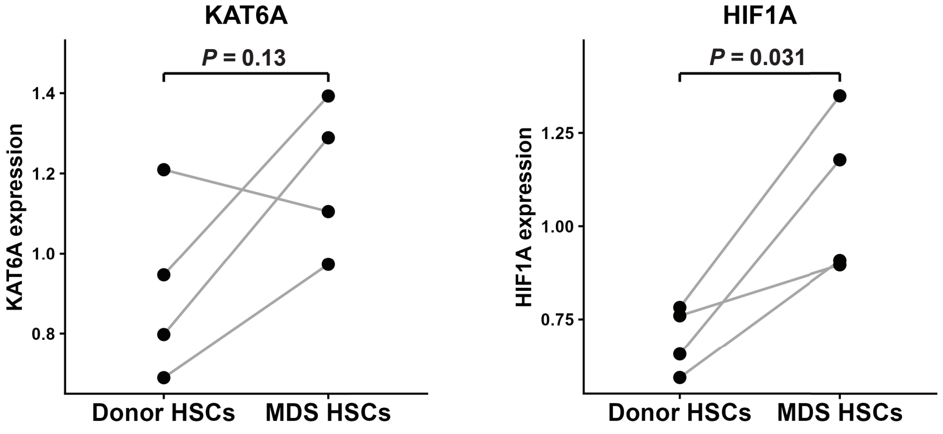
KAT6A
P = 0.13
KAT6A expression
1.4
1.2
1.0
0.8
Donor HSCs
MDS HSCs
HIF1A
P = 0.031
HIF1A expression
1.25
1.00
0.75
Donor HSCs
MDS HSCs

**Supplementary Figure 14. Genes and pathways commonly dysregulated in MDS HSCs following transplant.** (**A**) Violin plots showing the distribution of ribosomal protein expression module scores in matched donor (blue) and host (red) cells across cell-cycle phases (G1, S, G2M) for each patient. Scores were calculated using expression of ribosomal protein genes (RPS/RPL/RPLP). Significance was assessed using two-sided Wilcoxon rank-sum tests within each patient and phase ($P < 0.05$, $P < 0.01$, $P < 0.001$; ns, not significant). (**B**) Boxplots showing ribosomal module score based on expression of RPS/RPL/RPLP genes across hematopoietic populations (HSC/MPP, LMPP, cycling progenitor, MEP, early erythroid, early lymphoid, early GMP, late GMP) in **UTSW 1706**, stratified by cell origin/disease state (Host WT cells, MDS Pre-HCT, MDS Relapse, Donor WT). Each point represents a single cell; boxes indicate median and interquartile range. Statistical analysis comparing differences within each group was performed using Wilcoxon rank-sum tests with Benjamini–Hochberg correction, with adjusted p-values shown. (**C**) Gene set enrichment analysis (GSEA) of clonal hematopoiesis (CH)- and aging-associated gene signatures in MDS versus matched donor HSC/MPP/LMPPs; normalized enrichment score (NES) shown, with significance denoted by adjusted P values (*$P<0.05$, **$P<0.01$, ***$P<0.001$). (**D-E**) Paired comparison of mean MALAT1, NEAT1 (**D**), KAT6A, and HIF1A (**E**) expression in donor versus MDS HSC/MPP/LMPP cells. Each point represents the patient-level mean, with lines connecting matched donor–MDS pairs. P values were calculated using a two-sided paired t-test.
