## Supplemental Tables 1-8 for "Persistent Malignant Stem Cells Drive Relapse After Transplantation in the Myelodysplastic Neoplasms"

**Supplementary Tables**

**Supplementary Table 1**

**Flow cytometry gating:**

| Population | Gating Strategy |
|---|---|
| Restricted Progenitor Cells | $Lin^-CD34^+CD38^+$ |
| Lymphoid-Primed Multipotent Progenitor (LMPP) | $Lin^-CD34^+CD38^-CD45RA^+CD90^-$ |
| Multipotent Progenitor (MPP) | $Lin^-CD34^+CD38^-CD45RA^-CD90^-$ |
| Hematopoietic Stem Cell (HSC) | $Lin^-CD34^+CD38^-CD45RA^-CD90^+$ |

**Supplementary Table 2**

**Low Input DNA-sequencing Panels:**

**Myeloid Panel**
**Total genes:** 261
**Total probes:** 11736
**Genes:**
ADCY5, ANK2, APOB, SHROOM2, ATP2B3, DST, C5, CACNA1B, CACNA1E, CALR, RUNX1, RUNX1T1, CBFB, CBL, SCARB1, CD74, LRBA, CEBPA, CHD4, COL12A1, VCAN, DDX11, DNAH5, DNAH9, DNMT3A, DNMT3B, DOCK2, DRD2, DSCAM, GPR183, CELSR3, MEGF8, EGFR, ETV6, EZH2, FLG, FLT1, FLT3, GAS6, GATA2, GJB3, GRID1, GRIK2, GRIK4, GRM3, GRM8, HIVEP1, HNRNPK, TNC, IDH1, IDH2, ITPR3, JAK1, JAK2, JAK3, KCNA4, KCNH2, KCNQ2, KDR, KIT, KRAS, KRT19, MAP1B, MAP2, ADAM11, MEFV, MYC, MYH4, MYO5B, MYOC, NF1, NPM1, NRAS, NTRK3, DDR2, P2RY2, PKHD1, PPP1R3A, PTCH1, PTPN11, PTPRN, RAD21, RBBP4, RFC3, RYR1, RYR3, SCN1A, SRSF2, TRA2B, SHC1, SI, SLC12A3, STRN, NR2E1, TP53, TUBA3C, TYK2, U2AF1, KDM6A, WT1, USP9X, SMC1A, DYSF, CUL1, STC2, CADPS, EED, SYNGAP1, FCGBP, PRPF4B, CACNA1G, BSN, TOP3B, PKD2L1, SMC3, DCLK1, MTA2, NRXN3, ABCG2, CROCC, MAGI2, THRAP3, MED12, ZBTB33, EDIL3, SPEG, SEMA3A, SCML2, DLC1, PRPF8, STAG2, PTPRT, AKAP13, SCAF8, DIS3, RIMS1, SPEN, SMG1, HECW1, ATP10B, PSME4, MTUS2, SF3B1, KIAA0240, SUZ12, FLRT2, FKBP8, SETBP1, GIGYF2, LRIT1, DNAI1, FOXP1, C10orf28, CECR2, PCDHB1, COL5A3, PLCE1, WAC, DDX41, KDM3B, LRP1B, CNTN5, PCDHB18, HYDIN, TET2, DCHS2, TMEM104, BCOR, PHIP, ATG16L1, C10orf118, MTMR8, CACNA2D3, PCDHA13, PCDHA6, KCNK13, NMUR2, VARS2, FAM40B, PLEKHH1, ALPK3, KCNT1, ZNF687, KIAA1529, RNF213, NLRC4, MLL3, LRRC4, SEMA4A, IKZF4, CSMD1, PRAMEF2, FAM65A, DYNC2H1, E2F8, TRPM3, TET1, ABTB1, KIAA1683, EPPK1, CRISPLD1, FAM57B, SYT15, HMCN1, PHF6, PDCD2L, TTBK1, KIF2B, LNX1, TCEAL3, CNTNAP4, NAV1, PKHD1L1, MUC16, PKD1L2, CSMD3, GBP4, ARAP2, ZC3H18, MYOM3, GPR112, KCNU1, FAM47A, TCEAL6, CCDC67, XIRP1, BMPER, ASXL1, CMYA5, UNC5B, OR8B12, PHACTR1, CADM2, NALCN, BOD1L, SLC39A5, KSR2, FAM154B, KIAA1267, EPHA10, FRYL, ILDR1, KRT79, FAM5C, FREM2, OR13H1, FAM70B, GSTK1, GALNTL4, C17orf97, CUEDC1, OR11H12, CLEC18B, MUC5B, MROH5, SBF1P1, MIR142, EEF1A1P29, HSP90B3P, IGHG3, ST13P13

**MHC panel**
**Total Genes:** 11
**Total Probes:** 1181
**Genes:**
HLA-DPA1, HLA-DRB1, HLA-DRB4, HLA-DRB5, HLA-DPB1, HLA-DQA1, HLA-DRB3, HLA-A, HLA-B, HLA-C, HLA-DQB1

**Supplementary Table 3**

**Low Input Sensitivity Experiment:**

Germline Variants (Variants with population freq. > 25% in 1000g Project)

| CellCount | VAFbatch | # variants | Detected in 20% | Detected in 40% | Detected in 60% | Detected in 80% | Detected in 100% |
|---|---|---|---|---|---|---|---|
| 5000 | 2 to 5 | 3 | 100.0 | 100.0 | 100.0 | 100.0 | 100.0 |
| 5000 | 5 to 10 | 12 | 100.0 | 100.0 | 100.0 | 100.0 | 100.0 |
| 5000 | 10 to 20 | 15 | 100.0 | 100.0 | 100.0 | 100.0 | 100.0 |
| 5000 | 20 to 50 | 97 | 100.0 | 100.0 | 100.0 | 100.0 | 100.0 |
| 5000 | over 50 | 117 | 100.0 | 100.0 | 100.0 | 100.0 | 100.0 |
| 500 | 2 to 5 | 3 | 100.0 | 100.0 | 100.0 | 100.0 | 0.0 |
| 500 | 5 to 10 | 12 | 100.0 | 100.0 | 91.7 | 75.0 | 8.3 |
| 500 | 10 to 20 | 15 | 100.0 | 93.3 | 86.7 | 80.0 | 33.3 |
| 500 | 20 to 50 | 97 | 100.0 | 95.9 | 95.9 | 91.8 | 82.5 |
| 500 | over 50 | 117 | 100.0 | 100.0 | 98.3 | 90.6 | 82.1 |
| 200 | 2 to 5 | 3 | 100.0 | 100.0 | 100.0 | 33.3 | 0.0 |
| 200 | 5 to 10 | 12 | 100.0 | 83.3 | 66.7 | 58.3 | 16.7 |
| 200 | 10 to 20 | 15 | 100.0 | 100.0 | 93.3 | 80.0 | 73.3 |
| 200 | 20 to 50 | 97 | 96.9 | 96.9 | 94.8 | 88.7 | 74.2 |
| 200 | over 50 | 117 | 100.0 | 99.1 | 97.4 | 93.2 | 83.8 |
| 100 | 2 to 5 | 3 | 100.0 | 100.0 | 100.0 | 33.3 | 0.0 |
| 100 | 5 to 10 | 12 | 100.0 | 66.7 | 41.7 | 25.0 | 25.0 |
| 100 | 10 to 20 | 15 | 86.7 | 73.3 | 66.7 | 60.0 | 46.7 |
| 100 | 20 to 50 | 97 | 96.9 | 95.9 | 93.8 | 87.6 | 73.2 |
| 100 | over 50 | 117 | 99.1 | 96.6 | 94.0 | 88.9 | 84.6 |
| 50 | 2 to 5 | 3 | 100.0 | 100.0 | 0.0 | 0.0 | 0.0 |
| 50 | 5 to 10 | 12 | 100.0 | 83.3 | 33.3 | 8.3 | 0.0 |
| 50 | 10 to 20 | 15 | 100.0 | 86.7 | 60.0 | 33.3 | 13.3 |
| 50 | 20 to 50 | 97 | 100.0 | 96.9 | 90.7 | 69.1 | 38.1 |
| 50 | over 50 | 117 | 100.0 | 97.4 | 94.0 | 86.3 | 44.4 |
| 20 | 2 to 5 | 3 | 100.0 | 0.0 | 0.0 | 0.0 | 0.0 |
| 20 | 5 to 10 | 12 | 58.3 | 16.7 | 8.3 | 0.0 | 0.0 |
| 20 | 10 to 20 | 15 | 60.0 | 33.3 | 26.7 | 13.3 | 6.7 |
| 20 | 20 to 50 | 97 | 88.7 | 72.2 | 55.7 | 29.9 | 11.3 |
| 20 | over 50 | 117 | 94.9 | 82.1 | 67.5 | 57.3 | 42.7 |

Pathologic Variants

| CellCount | VAFbatch | # variants | Detected in 20% | Detected in 40% | Detected in 60% | Detected in 80% | Detected in 100% |
|---|---|---|---|---|---|---|---|
| 20 | 2 to 5 | 2 | 0 | 0 | 0 | 0 | 0 |
| 20 | 5 to 10 | 5 | 60 | 60 | 60 | 60 | 60 |

| | | | | | | | |
|---|---|---|---|---|---|---|---|
| 20 | 10 to 20 | 7 | 57.14286 | 57.14286 | 57.14286 | 28.57143 | 28.57143 |
| 20 | 20 to 50 | 3 | 100 | 66.66667 | 66.66667 | 0 | 0 |
| 50 | 2 to 5 | 2 | 100 | 0 | 0 | 0 | 0 |
| 50 | 5 to 10 | 5 | 100 | 100 | 60 | 60 | 60 |
| 50 | 10 to 20 | 7 | 100 | 71.42857 | 71.42857 | 42.85714 | 42.85714 |
| 50 | 20 to 50 | 3 | 100 | 100 | 100 | 33.33333 | 33.33333 |
| 100 | 2 to 5 | 2 | 50 | 50 | 0 | 0 | 0 |
| 100 | 5 to 10 | 5 | 100 | 100 | 80 | 80 | 60 |
| 100 | 10 to 20 | 7 | 100 | 100 | 100 | 71.42857 | 57.14286 |
| 100 | 20 to 50 | 3 | 100 | 100 | 100 | 100 | 100 |
| 200 | 2 to 5 | 2 | 100 | 100 | 50 | 0 | 0 |
| 200 | 5 to 10 | 5 | 100 | 100 | 100 | 100 | 80 |
| 200 | 10 to 20 | 7 | 100 | 100 | 100 | 100 | 85.71429 |
| 200 | 20 to 50 | 3 | 100 | 100 | 100 | 100 | 100 |
| 500 | 2 to 5 | 2 | 100 | 100 | 50 | 0 | 0 |
| 500 | 5 to 10 | 5 | 100 | 100 | 100 | 100 | 60 |
| 500 | 10 to 20 | 7 | 100 | 100 | 100 | 100 | 85.71429 |
| 500 | 20 to 50 | 3 | 100 | 100 | 100 | 100 | 100 |
| 5000 | 2 to 5 | 2 | 100 | 100 | 100 | 100 | 100 |
| 5000 | 5 to 10 | 5 | 100 | 100 | 100 | 100 | 100 |
| 5000 | 10 to 20 | 7 | 100 | 100 | 100 | 100 | 100 |
| 5000 | 20 to 50 | 3 | 100 | 100 | 100 | 100 | 100 |

**Supplementary Table 4**

**Specimen information (Figure 2):**

**MSK-001**
Tx Date: 6/26/2014
Sample Date: 12/22/2014 (Day +179)
Relapse Date: 12/12/2016 (Day +900)

**UTSW 1706**
Tx Date: 1/12/2023
Sample Date: 4/6/2023 (Day +84)
Relapse Date: 7/12/2023 (Day +181)

**UTSW 537**
Tx Date: 1/22/2021
Sample Date: 6/17/2021 (Day +146)
Relapse Date: 11/30/2021 (Day +312)

**UTSW 2155**
Tx Date: 12/8/2023
Sample Date: 3/15/2024 (Day +98)
Relapse Date: 6/20/2024 (Day +195)

**Supplementary Table 5**

**Specimen information (Figure 4):**

**UTSW 557**
Sample: (pre) HMTB 1216, (post) HMTB1487
Population in Figure: Bulk and CD34+CD38- cells

Tx Date: 7/7/2021
Post-transplant sample date: 9/14/2021
Relapse Date: 11/4/2021
Total distinct SNPs: 1335

**UTSW 332**
Sample: (pre) HMTB 0680, (post) HMTB1127
Population in Figure: Bulk and CD34+CD38- cells

Tx Date: 2/5/2021
Post-transplant sample date: 5/18/2021
Relapse Date: 8/4/2021
Total distinct SNPs: 1591

5/18/21 Clinical chimerism testing: 80% Donor, 20% Host

**Supplementary Table 6**

**Mission Bio Panels:**

**46 nuclear genes**
ASXL1, ASXL2, ATM, BCOR, BRAF, C11orf65, CALR, CBL, CHEK2, DDX41, DNMT3A, ETV6, EZH2, FLT3, GATA2, GNAS, GNB1, IDH1, IDH2, JAK2, KDM6A, KIT, KRAS, MFSD11, NPM1, NRAS, PHF6, PPM1D, PTPN11, RAB31, RAD21, RUNX1, SETBP1, SF3B1, SMC1A, SORCS3, SRSF2, STAG2, TET2, TP53, U2AF1, WT1, ZRSR2

**15 mitochondrial genes**
MT-ND1, MT-ND2, MT-ND3, MT-ND4, MT-ND4L, MT-ND5, MT-ND6, MT-CO1, MT-CO2, MT-CO3, MT-ATP6, MT-ATP8, MT-CYB, MT-RNR1, MT-RNR2

**total Genes: 61 genes**

**Three Custom Panels:**
MYE + Mito: 380 amplicons
CLU + Mito: 252 amplicons
CO-471 + Mito: 215 amplicons

**Mission Bio ADT Panel:**

| Antibody | Barcode | ID |
|---|---|---|
| CD30 | TCAGGGTGTGCTGTA | 0028 anti-human CD30 |
| CD3 | CTCATTGTAACTCCT | 0034 anti-human CD3 |
| CD8 | GCGCAACTTGATGAT | 0046 anti-human CD8 |
| CD56 | TCCTTTCCTGATAGG | 0047 anti-human CD56 (NCAM) |
| CD45 | TCCCTTGCGATTTAC | 0048 anti-human CD45 |
| CD19 | CTGGGCAATTACTCG | 0050 anti-human CD19 |
| CD33 | TAACTCAGGGCCTAT | 0052 anti-human CD33 |
| CD11c | TACGCCTATAACTTG | 0053 anti-human CD11c |
| CD34 | GCAGAAATCTCCCTT | 0054 anti-human CD34 |
| CD138 | ACTCTTTCGTTTACG | 0055 anti-human CD138 (Syndecan-1) |
| CD90 | GCATTGTACGATTCA | 0060 anti-human CD90 (Thy1) |
| CD117 | AGACTAATAGCTGAC | 0061 anti-human CD117 (c-kit) |
| CD10 | CAGCCATTCATTAGG | 0062 anti-human CD10 |
| CD45RA | TCAATCCTTCCGCTT | 0063 anti-human CD45RA |
| CD123 | CTTCACTCTGTCAGG | 0064 anti-human CD123 |
| CD7 | TGGATTCCCGGACTT | 0066 anti-human CD7 |
| CD4 | TGTTCCCGCTCAACT | 0072 anti-human CD4 |
| CD44 | TGGCTTCAGGTCCTA | 0073 anti-mouse/human CD44 |
| CD14 | TCTCAGACCTCCGTA | 0081 anti-human CD14 |
| CD16 | AAGTTCACTCTTTGC | 0083 anti-human CD16 |
| CD25 | TTTGTCCTGTACGCC | 0085 anti-human CD25 |
| CD45RO | CTCCGAATCATGTTG | 0087 anti-human CD45RO |

| | | |
|---|---|---|
| IgG1 | GCCGGACGACATTAA | 0090 Mouse IgG1 |
| IgG2a | CTCCTACCTAAACTG | 0091 Mouse IgG2a |
| IgG2b | ATATGTATCACGCGA | 0092 Mouse IgG2b |
| CD5 | CATTAACGGGATGCC | 0138 anti-human CD5 |
| CD69 | GTCTCTTGGCTTAAA | 0146 anti-human CD69 |
| CD62L | GTCCCTGCAACTTGA | 0147 anti-human CD62L |
| HLA-DR | AATAGCGAGCAAGTA | 0159 anti-human HLA-DR |
| CD1c | GAGCTACTTCACTCG | 0160 anti-human CD1c |
| CD11b | GACAAGTGATCTGCA | 0161 anti-human CD11b |
| CD64 | AAGTATGCCCTACGA | 0162 anti-human CD64 |
| CD141 | GGATAACCGCGCTTT | 0163 anti-human CD141 (Thrombomodulin) |
| CD62P | CCTTCCGTATCCCTT | 0218 anti-human CD62P (P-Selectin) |
| FcεRIα | CTCGTTTCCGTATCG | 0352 anti-human FcεRIα |
| CD163 | GCTTCTCCTTCCTTA | 0358 anti-human CD163 |
| CD83 | CCACTCATTTCCGGT | 0359 anti-human CD83 |
| CD13 | TTTCAACGCCCTTTC | 0364 anti-human CD13 |
| CD2 | TACGATTTGTCAGGG | 0367 anti-human CD2 |
| CD303 | GAGATGTCCGAATTT | 0370 anti-human CD303 (BDCA-2) |
| CD38 | TGTACCCGCTTGTGA | 0389 anti-human CD38 |
| CD22 | GGGTTGTTGTCTTTG | 0393 anti-human CD22 |
| CD71 | CCGTGTTCCTCATTA | 0394 anti-human CD71 |
| CD304 | GGACTAAGTTTCGTT | 0406 anti-human CD304 (Neuropilin-1) |
| CD49d | CCATTCAACTTCCGG | 0576 anti-human CD49d |
| CD81 | GTATCCTTCCTTGGC | anti-human CD81 (TAPA-1) |
| CD133 | GTAAGACGCCTATGC | anti-human CD133 |
| CD41 | ACGTTGTGGCCTTGT | anti-human CD41 |
| CD85d | GAGAGCTTTCACTGC | anti-human CD85d (ILT4) |
| CD164 | CCTTACGAGCATTTC | anti-human164 |
| GPR56 | GAGGTTCGTTGTGGT | anti-human GPR56 |
| CD97 | CAGTTGTGTATGGTC | anti-human CD97 |
| CD96 | ATTCCCTTTCACTCT | anti-human CD96 (TACTILE) |
| CD53 | GACTTGGGATTCAGC | anti-human CD53 |
| CD184 | TCAGGTCCTTTCAAC | anti-human CD184 (CXCR4) |
| KCNJ3 | TATCTTTGTAGCCCT | anti-human KCNJ3 |
| CD274 | GTTGTCCGACAATAC | anti-human anti-human CD274 (PD-L1) |
| CD244 | TCGCTTGGATGGTAG | anti-human CD244 |
| CD200 | CACGTAGACCTTTGC | anti-human CD200 |
| CD74 | CTGTAGCATTTCCCT | anti-human CD74 |
| CD9 | GAGTCACCAATCTGC | anti-human CD9 |
| CD49f | TTCCGAGGATGATCT | anti-human CD49f |
| CD109 | CACTTAACTCTGGGT | anti-human anti-human CD109 |

| | | |
|---|---|---|
| CD47 | GCATTCTGTCACCTA | anti-human CD47 |
| CD82a | TCTGTCACTGCCACT | anit-humanCD82a |
| Notch3 | TCACCCTCTGCCTAAA | anti-human Notch 3 |
| CD93 | GGGATTTACACGTAA | anti-human CD93 |
| CD32 | GCTTCCGAATTACCG | anti-human CD32 |
| CD366 | TGTCCTACCCAACTT | anti-human CD366 (Tim3) |
| CD166 | CATAAGATTCCGAGC | anti-human CD166 |
| CD99 | ACCCGTCCCTAAGAA | anti-human CD99 |
| CD371 | CATTAGAGTCTGCCA | anti-human CD371 (CLL-1) |
| CD36 | TTCTTTGCCTTGCCA | anti-human CD36 |
| CD58 | GTTCCTATGGACGAC | anti-human CD58 |
| CD43 | GATTAACCAGCTCAT | anti-human CD43 |
| CD126 | TGATGGGAGCTTATC | anti-human CD126 |
| CD55 | TGGATGTACGAGATG | anti-human CD55 |
| CD18 | AGGTGGCTAGAAGAC | anti-human CD18 |
| CD20 | TTCTGGGTCCCTAGA | anti-human CD20 |
| CD24 | AGATTCCTTCGTGTT | anti-human CD24 |
| CD135 | CAGTAGATGGAGCAT | anti-human CD135 |
| CD11a | TATATCCTTGTGAGC | anti-human CD11a |
| CD235a | AGAGTATGTATGGGA | anti-human CD235a |
| CD35 | ACTTCCGTCGATCTT | anti-human CD35 |

**Supplementary Table 7**

**Identification of cell types by Abseq:**

| **Population** | **Abseq Gating Strategy** |
|---|---|
| CD34+ Lin− | Lineage$^{lo}$, CD34$^{hi}$ |
| CD34+CD38− | Lineage$^{lo}$, CD34$^{hi}$, CD38$^{lo}$ |
| CD34+CD38+ | Lineage$^{lo}$, CD34$^{hi}$, CD38$^{hi}$ |
| HSC | Lineage$^{lo}$, CD34$^{hi}$, CD38$^{lo}$, CD45RA$^{lo}$, CD90$^{hi}$ |
| MPP | Lineage$^{lo}$, CD34$^{hi}$, CD38$^{lo}$, CD45RA$^{lo}$, CD90$^{lo}$ |
| LMPP | Lineage$^{lo}$, CD34$^{hi}$, CD38$^{lo}$, CD45RA$^{hi}$, CD90$^{lo}$ |
| CMP | Lineage$^{lo}$, CD34$^{hi}$, CD38$^{hi}$, CD45RA$^{lo}$, CD123$^{hi}$ |
| GMP | Lineage$^{lo}$, CD34$^{hi}$, CD38$^{hi}$, CD45RA$^{hi}$, CD123$^{hi}$ |
| MEP | Lineage$^{lo}$, CD34$^{hi}$, CD38$^{hi}$, CD45RA$^{lo}$, CD123$^{lo}$ |
| T cells | CD34$^{lo}$, CD3$^{hi}$ |
| NK cells | CD34$^{lo}$, CD3$^{lo,}$ CD56$^{hi}$ |
| CD3+CD56+ (NKT-like) Cells | CD34$^{lo}$, CD3$^{hi,}$ CD56$^{hi}$ |
| CD4+ T cells | CD34$^{lo}$, CD3$^{hi}$, CD4$^{hi}$, CD8$^{lo}$ |
| CD8+ T cells | CD34$^{lo}$, CD3$^{hi}$, CD4$^{lo}$, CD8$^{hi}$ |
| Double-Negative T cells | CD34$^{lo}$, CD3$^{hi}$, CD4$^{lo}$, CD8$^{lo}$ |
| CD4+CD8+ T cells | CD34$^{lo}$, CD3$^{hi}$, CD4$^{hi}$, CD8$^{hi}$ |
| CD4 naive-like | CD34$^{lo}$, CD3$^{hi}$, CD4$^{hi}$, CD45RA$^{hi}$, CD62L$^{hi}$ |
| CD4 TCM-like | CD34$^{lo}$, CD3$^{hi}$, CD4$^{hi}$, CD45RA$^{lo}$, CD62L$^{hi}$ |
| CD4 TEM | CD34$^{lo}$, CD3$^{hi}$, CD4$^{hi}$, CD45RA$^{lo}$, CD62L$^{lo}$ |
| CD4 TEMRA | CD34$^{lo}$, CD3$^{hi}$, CD4$^{hi}$, CD45RA$^{hi}$, CD62L$^{lo}$ |
| CD8 naive-like | CD34$^{lo}$, CD3$^{hi}$, CD8$^{hi}$, CD45RA$^{hi}$, CD62L$^{hi}$ |
| CD8 TCM-like | CD34$^{lo}$, CD3$^{hi}$, CD8$^{hi}$, CD45RA$^{lo}$, CD62L$^{hi}$ |
| CD8 TEM | CD34$^{lo}$, CD3$^{hi}$, CD8$^{hi}$, CD45RA$^{lo}$, CD62L$^{lo}$ |
| CD8 TEMRA | CD34$^{lo}$, CD3$^{hi}$, CD8$^{hi}$, CD45RA$^{hi}$, CD62L$^{lo}$ |
| Clean B cells | Dump-B$^{lo}$, CD19$^{hi}$, |
| Hematogone 1 | Dump-B$^{lo}$, CD19$^{hi}$, CD34$^{hi}$, CD10$^{hi}$ |
| CD34− CD10+ B-cell branch | Dump-B$^{lo}$, CD19$^{hi}$, CD34$^{lo}$, CD10$^{hi}$ |
| Hematogones 2/3 | Dump-B$^{lo}$, CD19$^{hi}$, CD34$^{lo}$, CD10$^{hi}$, CD22$^{lo}$ |
| Mature B region | Dump-B$^{lo}$, CD19$^{hi}$, CD34$^{lo}$, CD10$^{lo}$, CD22$^{hi}$ |
| Plasma cells | CD138$^{hi}$, CD38$^{hi}$ |
| Myeloid lineage cells | Dump-Myl$^{lo}$, CD33$^{hi}$ |
| Mature myeloid cells | Dump-Myl$^{lo}$, CD33$^{hi}$, CD34$^{lo}$ |
| Monocytes | Dump-Myl$^{lo}$, CD33$^{hi}$, CD34$^{lo}$, CD11b$^{hi}$, CD45$^{hi}$, CD14$^{hi}$, CD64$^{hi}$ |
| Erythroid parent | Dump-Ery$^{lo}$, CD71$^{hi}$ |
| Early erythroid | Dump-Ery$^{lo}$, CD71$^{hi}$, CD34$^{hi}$, CD36$^{hi}$ |
| Late erythroid | Dump-Ery$^{lo}$, CD71$^{hi}$, CD34$^{lo}$, CD36$^{hi}$ |
| Early proerythroblast | Dump-Ery$^{lo}$, CD71$^{hi}$, CD117$^{hi}$, CD45$^{lo}$ |
| Late proerythroblast / late erythroid | Dump-Ery$^{lo}$, CD71$^{hi}$, CD117$^{lo}$, CD45$^{lo}$ |
| Megakaryocytes | Dump-Plt$^{lo}$, CD45$^{lo}$, CD41$^{hi}$, CD9$^{hi}$ |

| | |
|---|---|
| Activated megakaryocytes | Dump-Plt$^{lo}$, CD45$^{lo}$, CD41$^{hi}$, CD9$^{hi}$, CD62P$^{hi}$ |
| Resting megakaryocytes | Dump-Plt$^{lo}$, CD45$^{lo}$, CD41$^{hi}$, CD9$^{hi}$, CD62P$^{lo}$ |

| ***In silico* pooled cocktails** | |
|---|---|
| Lineage | CD3, CD4, CD5, CD7, CD8, CD10, CD11b, CD14, CD19, CD22, CD56, CD71 |
| Dump-Myl | CD3, CD56, CD4, CD8, CD71, CD1 |
| Dump-B | CD3, CD56, CD14, CD33, CD11b, CD4, CD8, CD71 |
| Dump-Ery | CD3, CD4, CD8, CD2, CD5, CD7, CD19, CD22, CD10, CD56, CD244, CD11b, CD14, CD16, CD33, CD64, CD163, CD11c, CD13, CD18, CD32, HLA.DR, CD74, CD123, CD1c, CD303, CD304 |
| Dump-Plt | CD45, CD71, CD14, CD64, CD33, CD11b, CD16, CD163, CD11c, CD13, HLA.DR, CD3, CD19, CD56 |

**Supplementary Table 8**

**Summary of mutant cells identified by scDNA-seq (Figure 5):**

| **Patient** | **Variant(s)** | **Cell Count** | **Any mutation** | **All mutations** | **Per variant** |
|---|---|---|---|---|---|
| **SC85** | chr17:7577566:T/C | 1079 | 998/1079 (92.5%) | 998/1079 (92.5%) | chr17:7577566:T/C: 998/1079 (92.5%) |
| **SC30** | chr17:7577120:C/T chr17:7578268:A/T | 237 | 195/237 (82.3%) | 135/237 (57.0%) | chr17:7577120:C/T: 177/237 (74.7%); chr17:7578268:A/T: 153/237 (64.6%) |
| **UTSW 2810** | chr17:7578407:G/C | 11 | 8/11 (72.7%) | 8/11 (72.7%) | chr17:7578407:G/C: 8/11 (72.7%) |
| **SC79** | chr17:7578203:C/T | 7 | 7/7 (100.0%) | 7/7 (100.0%) | chr17:7578203:C/T: 7/7 (100.0%) |
| | Total: | 1334 | 1208/1334 (91%) | | |
